# A new role for lipoproteins LpqZ and FecB in orchestrating mycobacterial cell envelope biogenesis

**DOI:** 10.1101/2025.06.19.660376

**Authors:** Robin Lissner, Aaron Franklin, Samuel T. Benedict, Vicky Charitou, Alexander Speer, Jaco Knol, Connie Jimenez, Patrick J. Moynihan, Sergey Nejentsev, Coen Kuijl, Wilbert Bitter

**Affiliations:** Molecular Microbiology A-LIFE department, Vrije Universiteit Amsterdam, the Netherlands; Medical Microbiology and Infection Control, Amsterdam UMC, Amsterdam, the Netherlands; Department of Medicine, University of Cambridge, Cambridge, UK; Medical Oncology, Amsterdam UMC, Amsterdam, the Netherlands; Molecular Cell Biology and Immunology, Amsterdam UMC, Amsterdam, the Netherlands; Amsterdam Institute for Immunology & Infectious diseases, Amsterdam, the Netherlands; School of Biosciences, University of Birmingham, Birmingham, UK; Department of Microbiology and Immunology, University of Western Ontario, London, Ontario, Canada

## Abstract

Accounting for more deaths than any other bacterial species, *Mycobacterium tuberculosis* (Mtb) represents a critical threat to public health worldwide. A key factor contributing to Mtb’s virulence is its unique cell envelope, which acts as a protective barrier. Among the components of this envelope, lipoproteins represent a critical but understudied group of proteins. In this study, we focused on 79 conserved putative lipoproteins, shared between Mtb and the closely related *M. marinum*. Leveraging the CRISPR1-Cas9 (Sth1Cas9) gene editing system for Mycobacteria, we generated frameshift mutations, targeting one conserved lipoprotein-coding gene at a time. We identified two mutants, *lpqZ* and *fecB*, that exhibited increased susceptibility to all tested antibiotics, suggesting essential roles in cell envelope biogenesis. Interestingly, despite having homology to periplasmic substrate binding proteins (SBPs), neither protein is associated with any inner membrane transporter complex. Instead, co-immunoprecipitation experiments revealed that LpqZ interacts with AftA and FecB interacts with AftB. Both these interaction partners are essential enzymes involved in arabinogalactan and lipoarabinomannan synthesis. Accordingly, we observed alterations for both glycoconjugates in *lpqZ* and *fecB* mutants. Together, these findings show that orphaned SBP-like proteins have been neofunctionalized in mycobacteria to aid key enzymes involved in cell envelope biosynthesis.

## Introduction

Accounting for more deaths than any other bacterial species, *Mycobacterium tuberculosis* (Mtb) represents a critical threat to public health worldwide. Maybe the most crucial and distinctive feature of mycobacteria is their unusual cell envelope. Unlike other bacteria, mycobacteria possess a complex cell envelope with an additional, unique outer membrane, the so called mycomembrane (Fig. 1a). This extremely hydrophobic membrane is characterized by the presence of long-chain mycolic acids ^1^. Most intriguingly, the mycolic acids of the inner leaflet are covalently linked to the key polysaccharide arabinogalactan, which in turn is linked to the peptidoglycan, collectively forming an enormous macro polymer^2^. This core structure of the cell wall is also referred to as the mycolyl-arabinogalactan-peptidoglycan (mAGP) complex. The outer leaflet of the mycomembrane is composed of various other lipids, including lipoglycans and mycolic acids linked to trehalose ^1^. The two most abundant lipoglycans lipoarabinomannan (LAM) and its precursor lipomannan (LM) are synthesized at the plasma membrane and transported to the outer membrane and the capsule, where they can be exposed to the cell surface or released as lipid-free glycans by enzymatic cleavage ^3,4^. LM and LAM are not only important for mycobacterial pathogenesis inside the host ^5^, but have also been suggested to play critical roles in bacterial physiology by modulating cell envelope integrity and regulating cell division ^6,7^. Altogether, the complex mycobacterial cell wall does not only supply mechanical resistance but also protects the bacteria from harmful agents, such as antibiotics and effector molecules of the immune system ^8^. To overcome this barrier and to fight Mtb it is crucial to understand the physiology of the cell envelope and its proteins ^9^.

**Figure 1:**
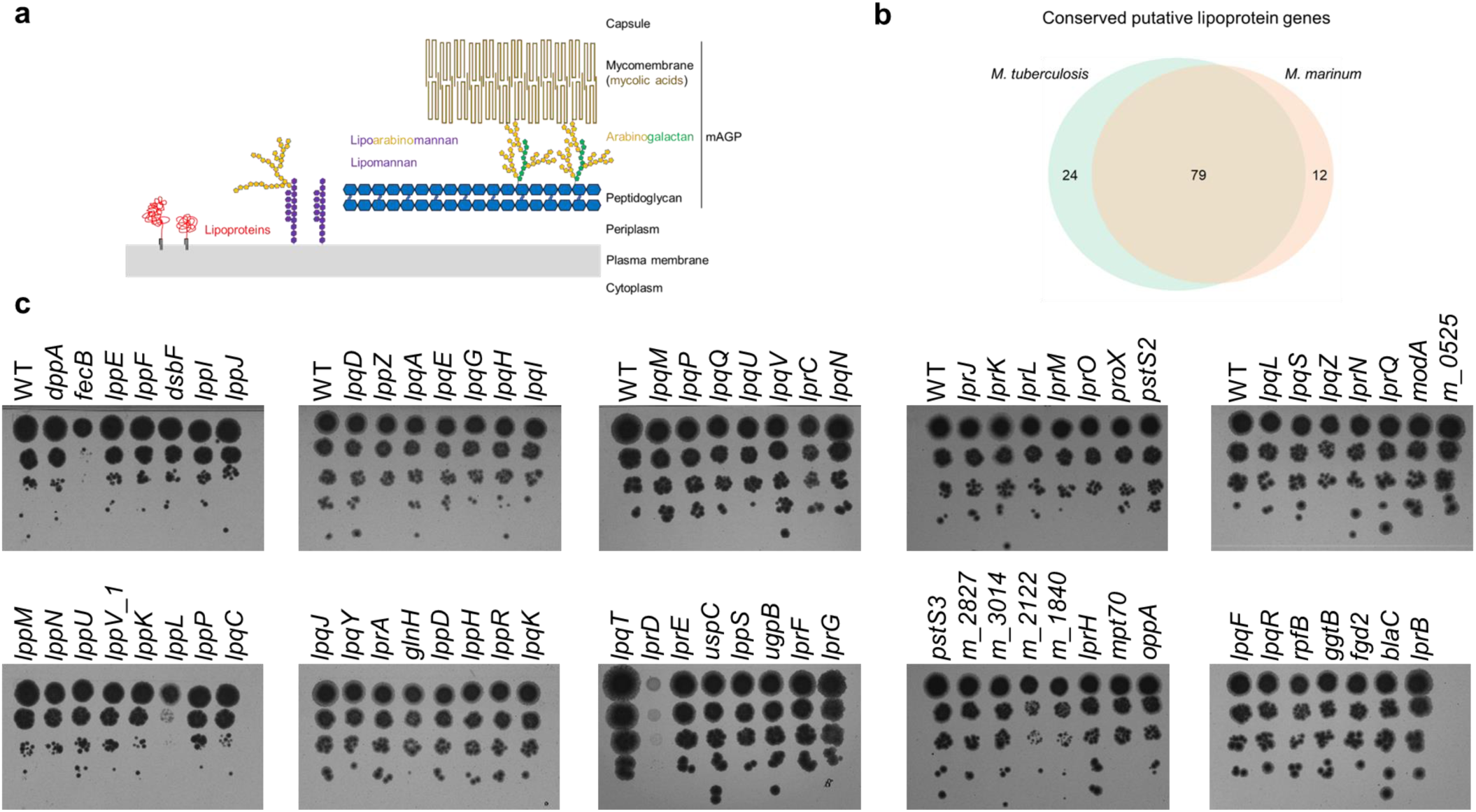
CRISPR/Cas9 mutants of conserved *M. marinum* lipoproteins. (**a**) the mycobacterial cell envelope and its main components: long-chain mycolic acids form the unique outer membrane (mycomembrane). Mycolic acids of the inner leaflet are covalently linked to a layer of arabinogalactan, which in turn is linked to the peptidoglycan. This macropolymer can be referred to as the mycolyl-arabinogalactan-peptidoglycan (mAGP) complex. The two most abundant glycolipids lipoarabinomannan and its precursor lipomannan are synthesized at the plasma membrane and can be transported to the outer membrane. Lipoproteins are a key group of membrane proteins that we believe to play important roles for the integrity of the cell envelope (**b**) putative lipoprotein genes in *M.tuberculosis* and *M.marinum* and their conserved fraction. (**c**) wild-type (*M. marinum* WT pCRISPRx-Sth1-Cas9 L5 empty) and 74 successfully sequence-verified *M. marinum* lipoprotein CRISPR/Cas9 frameshift mutants were adjusted to OD 0.1 and spotted on 7H10 kanamycin plates in 10-fold serial dilutions.

The highly organized and dynamic cell envelope needs to be build, maintained and controlled, which requires the function of many different enzymes and accessory proteins ^8^. One group of proteins that could play a role in this process are the lipoproteins, which are inserted into the membrane with their lipid anchor ^10^. They can be characterized by their N-terminal secretion signal peptide with the lipobox motif. The three enzymes Lgt, LspA and Lnt replace this signal sequence with a three fatty acid lipid anchor at a universally conserved cysteine residue within the lipobox ^11^. Based on signal peptide sequence pattern searches, the genome of Mtb harbors 99 genes that could potentially code for lipoproteins ^12^. Some of the mycobacterial lipoproteins are well characterized. For example, RpfB ^13,14^, LpqW ^15^ and LprG ^16,17^ have vital roles in cell envelope biogenesis, whereas LpqH ^18–21^ and the mce1 associated LprK lipoprotein ^22^ have been shown to be important virulence factors. However, the function of the majority of lipoproteins remains poorly understood. Further exploration is crucial for identifying additional lipoproteins with key roles in the functioning of the mycobacterial cell envelope or the interaction with the environment, including the human host. Identifying and understanding these lipoproteins could pave the way for novel antibiotic targets and advancements in vaccine development.

Here, we studied the 79 putative lipoproteins that are conserved between *M. tuberculosis* and the closely related model organism *Mycobacterium marinum*. Using the CRISPR1-Cas9 (Sth1Cas9) gene editing system for mycobacteria ^23^, we introduced frame shift mutations in all conserved lipoprotein-coding genes, targeting one gene at the time. We found two mutants, *lpqZ* and *fecB* that showed increased susceptibility to all tested antibiotics, pointing towards critical roles in cell envelope biogenesis. Using co-immunoprecipitation (Co-IP) followed by mass spectrometry (MS), we identified interacting partners of these two lipoproteins, indicating that they are key players in LAM and arabinogalactan synthesis, thus explaining the observed phenotypes.

## Results

### Construction of lipoprotein CRISPR/Cas9 frameshift mutants

Currently, the Mycobrowser database (https://mycobrowser.epfl.ch/), which serves as a genomic and proteomic resource for mycobacteria, lists 87 lipoproteins. In addition, through signal peptide sequence pattern searches conducted by Sutcliffe and Harrington (2004), we have compiled a list of 103 putative lipoproteins specific to *M. tuberculosis*, 79 of which are conserved in *M. marinum* (Fig. 1b, Supplementary Table 1). We targeted these conserved genes in *M. marinum,* because it is a BSL-2 pathogen with similar pathogenesis as *M. tuberculosis* in its natural host, poikilothermic vertebrates. Its close relationship to *M. tuberculosis* is reflected in its average amino acid identity of over 85% between orthologues, making it a useful model organism ^24^. We utilized the *Streptococcus thermophilus* CRISPR1- Cas9 (Sth1Cas9) gene editing system for mycobacteria, which has demonstrated high precision, no off-target effects, and strong efficiency - consistently yielding non-functional genes in all tested frameshift mutants ^23^. Targeting the genes with specific gRNAs one by one allowed us to individually select and verify clones with out-of-frame mutations using PCR and Sanger sequencing of the respective amplicon. Successful disruption of the ORF was achieved for 74 of the targeted genes. Three genes (*lprD, lppL and fecB*) showed a strong, visible growth defect on 7H10 agar plates (Fig. 1c). Additionally*, mmar_2122* and *mmar_1840* mutants showed slightly smaller colonies and the *lprG* mutant had a different colony morphology than the wild-type (WT) *M. marinum* strain, similar to the previously reported mutant in *M. tuberculosis* ^16,25^.

After multiple unsuccessful attempts to generate out-of-frame mutations in the five targeted lipoprotein genes *lpqB, lpqW, lppW, lppX* and *subI*, we decided to continue our research using a verified library of 74 lipoprotein mutants.

### Identification of key players in the cell envelope in a pooled antibiotic susceptibility assay

To identify lipoproteins involved in the mycobacterial cell envelope biogenesis and function, we screened our library of mutant strains by assessing their susceptibility to antibiotics that either target the cell envelope itself or depend on its permeability for activity. To accomplish this in a high-throughput manner, we employed a pooled library approach. Since our library contains only a limited set of verified mutants, we expect to have minimized bottle-neck issues and no bias towards big ORFs like in Tn-based methods. We exposed our library to a range of antibiotic concentrations, selecting the plates where bacterial growth was still evident, though at a noticeable reduction. After DNA extraction and amplification of the gRNA locus, next generation sequencing was used to quantify the presence of mutants in each individual pool using the integrated pCRISPRxSth1Cas9-L5 as unique barcodes (Fig. 2a). As a control for mutants with a general growth defect we also compared read counts from the calibrated input pool and compared them to the read counts from the mixture of bacterial mutants grown on 7H10 plates without antibiotics. To assess antibiotic susceptibility, the read counts from the non-antibiotic plates were compared to read counts of plates containing the respective antibiotics. As observed earlier, the *lprD, lppL* and *fecB* mutant showed a strong general growth defect on 7H10 plates (Fig. 2b), confirming the validity of the approach. Interestingly, while antibiotic treatment did not further impair the growth of *lprD* and *lppL* mutants, the *fecB* mutant exhibited increased susceptibility across all antibiotics tested. This observed cross-sensitivity aligns with previous findings in a *fecB* mutant of *M. tuberculosis* ^26^. Apart from *fecB*, also the *rpfB*, *mmar_2122, mmar_1840, lprG,* and *lpqZ* mutants were strongly depleted after exposure to the antibiotics (Fig. 2b). In contrast, the *lprQ, lprD, lprC, lprB, lpqI* and *lppL* mutants were significantly more resistant to augmentin than the wild-type. We confirmed the observations of our pooled screen by testing individual strains on agar plates containing different antibiotic dilutions (Fig. 2c, Supplementary Fig. 1).

**Figure 2:**
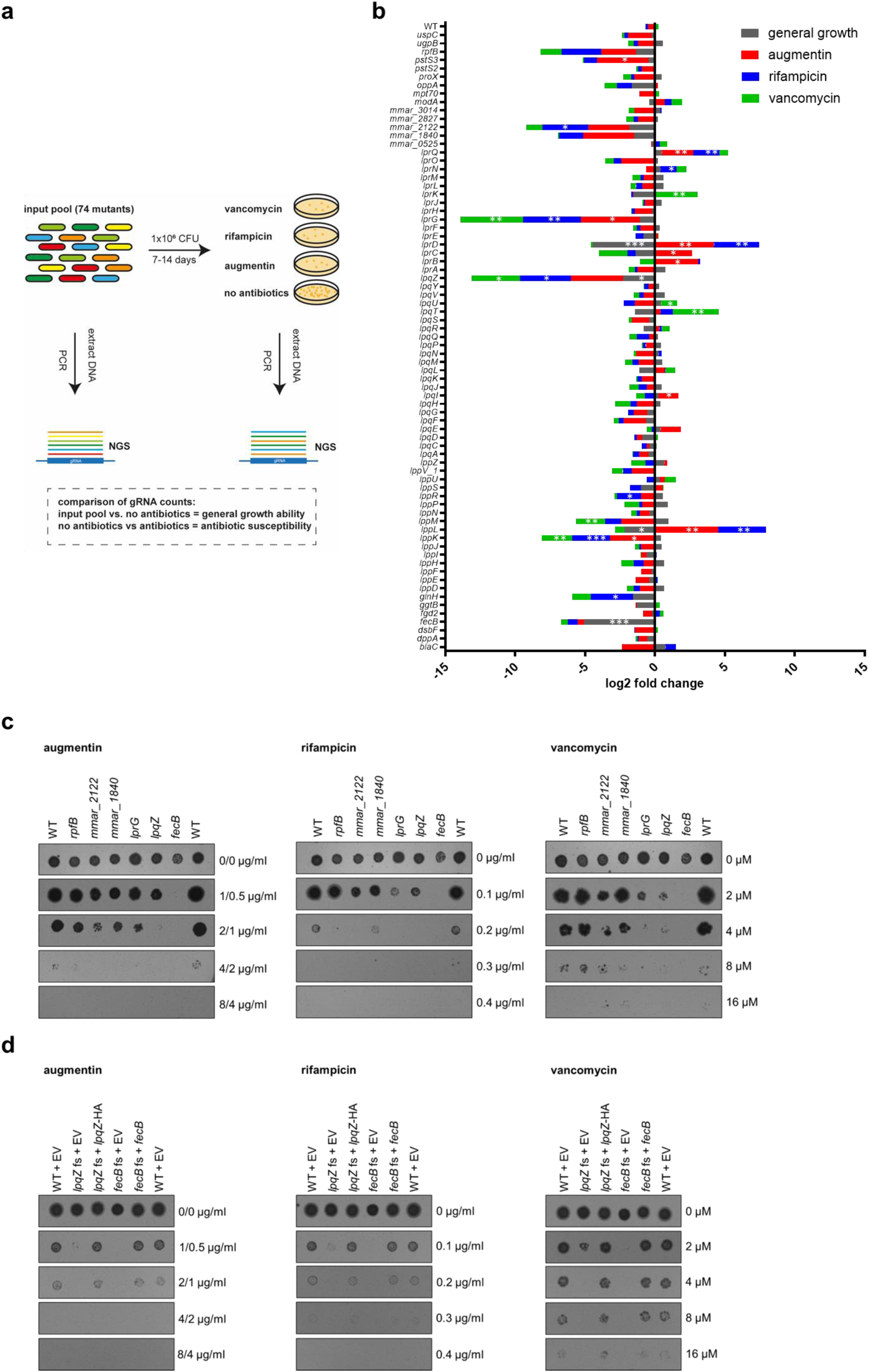
pooled susceptibility testing of lipoprotein mutants. (**a**)74 lipoprotein CRISPR/Cas9 frameshift mutants and *M. marinum* wild-type strain (WT pCRISPRx-Sth1- Cas9-L5 empty) were pooled together and incubated for 7-14 days on 7H10 plates with vancomycin, rifampicin, augmentin or without antibiotics. Bacterial DNA was extracted and the gRNA locus was amplified by PCR for next generation sequencing (NGS). General growth ability was assessed by comparison of gRNA counts of input pool and 7H10 plates without antibiotics. Antibiotic susceptibility was assessed by comparison of gRNA counts from 7H10 plates and plates with the respective antibiotics. (**b**) relative gRNA counts (log2 fold change) of 74 lipoprotein mutants and the WT strain (alphabetical order). Data shows the mean of two independent experiments. Asterisks indicate p-values as determined by a drugZ test (*p< 0.05, ** p< 0.01, ***p< 0.001). (**c**) individual strains from (b) were spotted on 7H10 plates containing different concentrations of the respective antibiotics. (**d**) genetically complemented *lpqZ* fs mutant (*lpqZ* fs + *lpqZ*-HA) and complemented *fecB* fs mutant (*fecB* fs + *fecB*) were compared to WT and fs mutants containing the empty pSMT3 vector (EV) for antibiotic susceptibility as in (c).

RpfB, LprG and LpqI are of known function and have been associated with roles that are crucial for cell envelope integrity, such as the resuscitation from the dormant state ^13,14^, triacylglyceride and lipoarabinomannan transport ^16,17^ and peptidoglycan recycling ^27^, respectively. However, such a functional link with cell envelope integrity was not known for *lpqZ* and *fecB*. To explore possible functions of these genes, we first employed structural similarity searches using Foldseek. We found that both LpqZ and FecB have homology to periplasmic substrate binding proteins (SBPs), which are often linked to ABC transporters. LpqZ shares 23.7% amino acid identity with ProX (*M. tuberculosis*) (Supplementary Table 2). ProX is part of the ProXVWZ operon, which is suggested to be responsible for osmoregulation in *M. tuberculosis* ^28,29^. On the other hand, FecB shares 20% identity with FecB of E.coli and 20.8% sequence identity with SirA of *Staphylococcus aureus* (Supplementary Table 2). Both *E.coli* FecB and SirA are involved in iron acquisition ^30,31^, however, in mycobacteria, the role of FecB is more controversial. While one study suggests that FecB has heme- and mycobactin binding properties ^32^, Xu et al. showed that FecB is not required for iron acquisition in *M. tuberculosis*^26^. In contrast to *proX*, *fecB* (*E.coli*) and *sirA*, both *lpqZ* and *fecB* are not part of a gene cluster/operon coding for an ABC transporter and are therefore considered orphan genes ^33^. This could mean that they have evolved novel functions, e.g. in the organization of the mycobacterial cell envelope. We therefore focused on LpqZ and FecB for further functional investigation.

Expression of *lpqZ* and *fecB* under the control of the HSP60 promotor successfully restored antibiotic susceptibility in the respective mutant strains to wild-type levels (Fig. 2d).

Previously, a *fecB* mutant has been demonstrated to exhibit increased cell wall permeability, potentially accounting for its increased susceptibility to antibiotics ^26^. To test this, we assessed the permeability of the *fecB* and *lpqZ* mutants to ethidium bromide (EtBr). Both mutants displayed only a slightly elevated permeability compared to the wild-type strain (Supplementary Fig. 2). While this observation supports a role in cell envelope organization, the modest increase in EtBr uptake suggests that changes in general permeability alone are unlikely to fully account for the pronounced antibiotic susceptibility. This implies that LpqZ and FecB might play more specific roles in the correct assembly, localization, or functionality of particular cell envelope components.

### LpqZ and FecB are interacting with arabinosyltransferases AftA and AftB, respectively

To elucidate the function of LpqZ and FecB, we aimed to identify their binding partner proteins by co-immunoprecipitation (co-IP) from n-dodecyl-β-D-maltoside (DDM)- solubilized membrane proteins, followed by mass spectrometry (MS). Comparative analysis of LFQ intensities revealed that LpqZ was found to be enriched over 5000-fold in the purified LpqZ- HA samples as compared to the wild-type control. Notably, the most enriched co-precipitated protein with LpqZ was AftA (MMAR_5354), which exhibited a 20,000-fold enrichment relative to the wild-type samples (Fig. 3a). AftA of *M. tuberculosis* encodes an arabinosyltransferase (Rv3792), which is essential for *in-vitro* growth and responsible for the priming step of arabinogalactan synthesis by transferring the first arabinosyl residue to the galactan chain ^34^.

**Figure 3:**
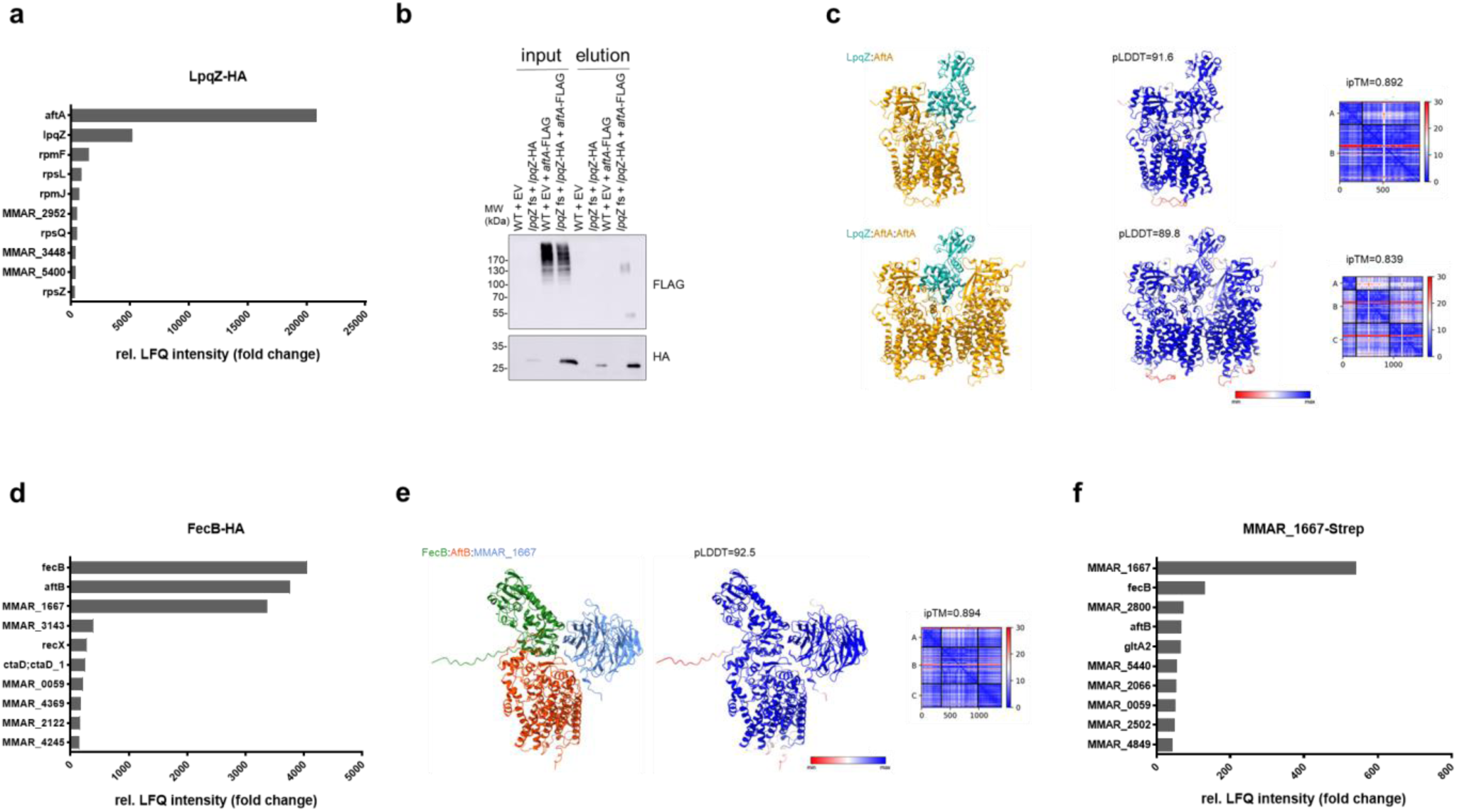
finding putative protein interaction partners of LpqZ and FecB. (**a,d,f**) Top 10 hits of mass spectrometry (MS) relative LFQ intensity values from *M. marinum* LpqZ-HA, FecB-HA or MMAR_1667-Strep co-immunoprecipitation (co-IP) samples compared to wild- type (WT) samples. FecB-HA and MMAR_1667-Strep co-IP relative LFQ intensity is the average of two biological replicates. (**b**) Co-IP of *M. marinum* AftA-FLAG from LpqZ-HA, showing the input and elutions of the genetically complemented *lpqZ* fs mutant (*lpqZ* fs +*lpqZ*-HA) with and without expression of *aftA*-FLAG, compared to the WT with the empty psMT3 vector (EV). Co-IP samples were separated by SDS page and western blot was stained with anti-HA and anti-FLAG antibodies. (**c,e**) AlphaFold complex prediction of the putative *M. marinum* interaction partners, showing the complex colored by chain (left), colored by pLDDT (middle) and respective PAE plots with ipTM values (right).

The structure of AftA has been determined by cryo-EM and shown to form a homodimer with two cavities that converge at the active site ^35^. We confirmed the putative interaction of the two proteins by co-immunoprecipitation from membrane fractions of cell lysates expressing LpqZ-HA and AftA-FLAG (Fig. 3b). HA-purified LpqZ-HA successfully co-purified AftA-FLAG in both its monomeric and homodimeric forms, which confirms the observations by Gong *et al*.^35^. AlphaFold predictions further suggested that both monomeric and dimeric AftA can bind LpqZ in a complex, with overall a slightly better model confidence for the monomeric (ipTM=0.892) than for the dimeric (ipTM= 0.839) state (Fig. 3c). AlphaFold predictions of the *M. tuberculosis* orthologs of LpqZ (Rv1244) and AftA (Rv3792) showed similar confident scores for a dimer (ipTM=0.873) and a trimer (ipTM=0.881) (Supplementary Fig. 3), whereas a negative control of AftA:FecB showed a nonsignificant ipTM score of 0.41 (Supplementary Fig. 4).

The MS analysis of proteins that co-immunoprecipitated with FecB-HA resulted in a 4000- fold enrichment of FecB itself, closely followed by AftB (MMAR_5369) and MMAR_1667 with over 3000-fold enrichment (Fig. 3d). The arabinosyltransferase AftB is, just like AftA, essential for *in-vitro* growth ^36^ and is thought to be responsible for the terminal β(1 → 2) linkage of arabinosyl to both arabinogalactan ^37^ and lipoarabinomannan ^38^. MMAR_1667 has a *M. tuberculosis* orthologue Rv3035 with unknown function. AlphaFold predictions suggested with high confidence (ipTM=0.894) that FecB, AftB and MMAR_1667 form a trimeric complex (Fig. 3e). Similarly, the *M. tuberculosis* orthologs FecB (Rv3044), AftB (Rv3805c) and Rv3035 showed confident scores (ipTM=0.864) for a trimeric complex (Supplementary Fig. 3), whereas the negative control of AftB:LpqZ showed a low confidence ipTM score of 0.312 (Supplementary Fig. 4). However, our attempts to co-purify AftB-FLAG from FecB-HA expressing cell lysates were unsuccessful. This could be due to steric hindrance of the C-terminal tags, which are predicted to be close to the protein interaction sites in the center of the trimeric complex (Supplementary Fig. 5a,b). This is supported by our observation that expression of FecB-HA was not able to complement the antibiotic susceptibility phenotype of the *fecB* mutant (Supplementary Fig. 5c), contrasting our earlier positive observations with the untagged protein (Fig. 2d). To address this technical issue, we decided to verify these interactions by repeating the co-IP with streptavidin-tagged MMAR_1667, followed by MS analysis. Given that the C-terminus of MMAR_1667 is located farther from the predicted interaction site with its partner proteins, we anticipated minimal interference from the tag. In confirmation with our previous findings, MMAR_1667 was enriched over 500-fold in our pulldowns, closely followed by FecB and AftB with 130-fold and 67-fold enrichment, respectively (Fig. 3f).

Next, we aimed to investigate the mutant phenotype of *mmar_1667* and compare it to those of its interaction partners. The disruption of this gene resulted in impaired growth on 7H10 agar plate, similar to the *fecB* mutant and to the *aftB* CRISPRi silencing (Supplementary Fig. 6a). The expression of *mmar_1667* under the control of an HSP60 promoter was able to complement the observed phenotype (Supplementary Fig. 6b). The similarity in growth disruption aligns with our co-IP data, further supporting the idea that MMAR_1667, FecB and AftB function together as interaction partners.

### FecB and LpqZ are involved in arabinogalactan synthesis

The interaction of LpqZ with AftA, as well as the interaction of FecB with AftB suggest that both lipoproteins support the function of these two enzymes in arabinogalactan synthesis. We cannot make frameshift mutants in these essential genes, but reducing the amount of the essential partner proteins using CRISPRi could lead to synergistic growth defects. To assess this, we silenced the expression of *aftA* or *aftB* in the respective *lpqZ* or *fecB* mutant backgrounds. We selected ATc concentrations that allowed residual growth, as higher levels resulted in complete growth arrest. Under these conditions, silencing *aftA* or *aftB* in the mutant backgrounds led to a slightly stronger growth reduction compared to silencing these arabinosyltransferases in the wild-type background, or silencing an unrelated essential lipoprotein gene (*lpqW*) in the mutant backgrounds (Supplementary Fig. 7a,b). The observed synergistic effects in growth reduction in the mutants therefore indicate a supportive function of the lipoproteins in arabinogalactan synthesis.

To directly assess alterations in arabinogalactan biosynthesis in the mutant strains, we extracted and hydrolyzed arabinogalactan from bacterial cell envelopes and analyzed the monosaccharide composition using high-performance anion-exchange chromatography (HPAEC). The relative abundance of arabinose and galactose in these hydrolysates serves as a proxy for arabinogalactan structure, with changes in the arabinose-to-galactose (ara:gal) ratio indicating possible disruptions in biosynthetic pathways. Both *fecB* and *lpqZ* mutants exhibited a reduced ara:gal ratio compared to wild-type *M. marinum* (Fig. 4a). This reduction suggests that both lipoproteins are involved in the incorporation or regulation of arabinose residues during arabinogalactan assembly.

**Figure 4:**
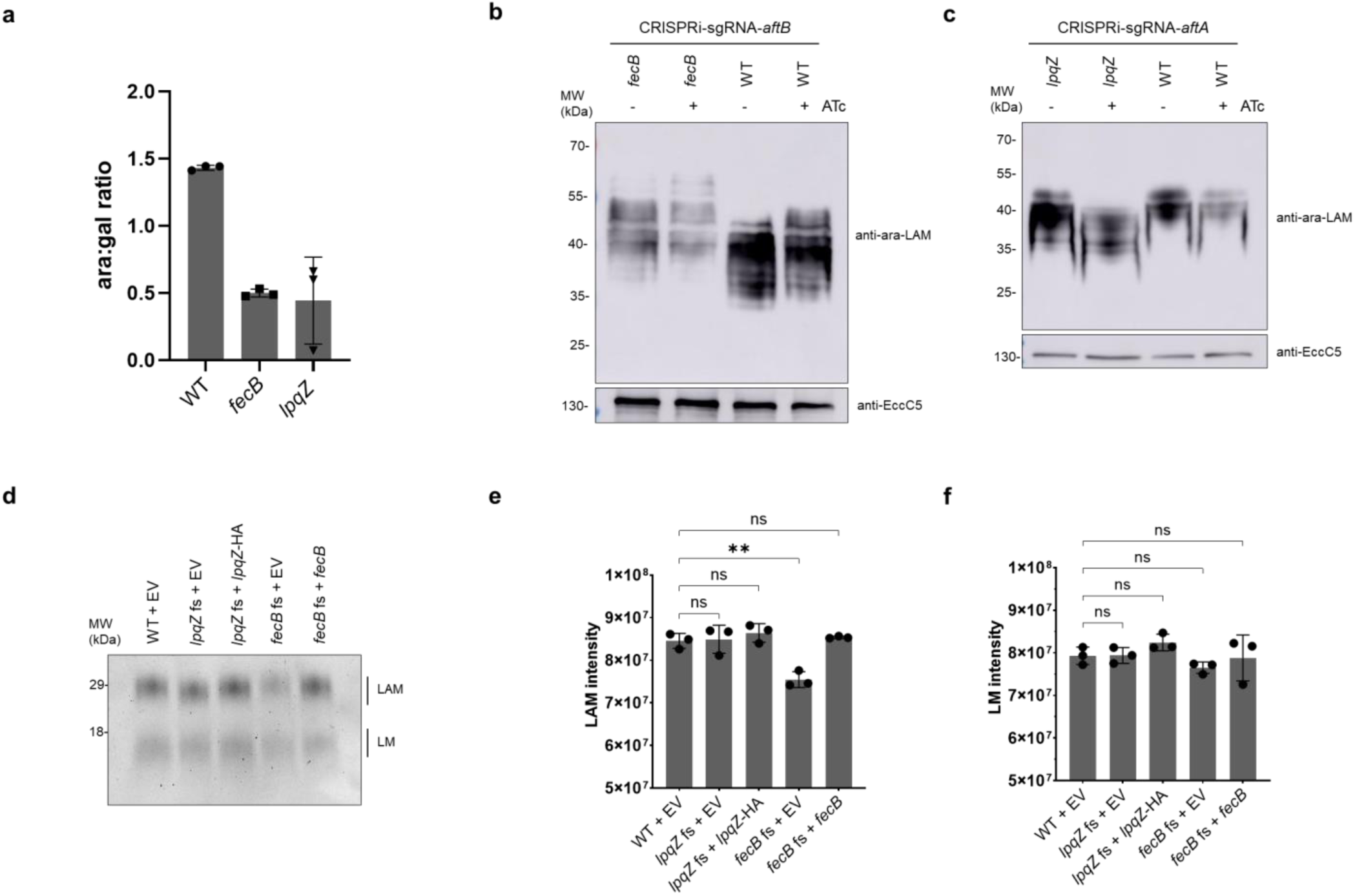
arabinogalactan and lipoarabinomannan profiles of *lpqZ* and *fecB* mutants. (**a**): arabinose to galactose ratios (ara:gal) were determined for *M. marinum* wild-type (WT), *fecB* and *lpqZ* fs mutants by extraction and digestion of arabinogalactan and subsequent HPAEC analysis. The data shows the mean of biological triplicates. Error bars show standard deviation (**b**): Whole cell lysates of *aftB* CRISPRi silencing in *M. marinum* WT and *fecB* fs mutant, with (+) and without (-) ATc induction were separated by SDS page. Western Blot was stained with Mab F30-5 antibody, recognizing terminal arabinan branches of LAM and anti-EccC5 as a loading control. Shown is a representative blot from two independent experiments (n=2) (**c**): *aftA* CRISPRi silencing in *M. marinum* WT and *lpqZ* fs mutant as in (b). Shown is a representative blot from two independent experiments (n=2) (**d**): representative LAM and LM extracts of *M. marinum* WT, *lpqZ* and *fecB* fs mutants, containing the empty pSMT3 vector (EV) and genetically complemented strains. Extracts were separated by SDS page and stained with Pro-Q Emerald. Means of three independent experiments (n=3) were calculated for LAM (**e**) and LM (**f**) intensities. Error bars show the standard deviation. Asterisks show the level of significance (** p<0.005, ns: not significant), determined by a one-way ANOVA with Dunnett’s multiple comparisons test.

The process of arabinogalactan synthesis and the enzymes involved, remains poorly understood. Among the key enzymes, AftA has been studied more extensively than AftB. AftA catalyzes the addition of the first arabinosyl residue to the galactan chain ^34^. In *M. tuberculosis*, the active site of AftA contains two conserved catalytic residues, D128 and R132, as experimentally demonstrated by Gong *et al*. ^35^. These residues are accessible for a putative decaprenyl-phosphoryl-arabinose (DPA) donor via a channel that leads to a binding cavity. Using molecular docking, Gong *et al*. proposed a possible acceptor binding site within the cavity by modeling the interaction of a synthetic eight-residue galactofuranosyl ligand. To test if this galactan chain can still bind the putative acceptor binding site in the presence of LpqZ, we performed docking experiments, following the approach of Gong *et al*. The HADDOCK software ^39,40^ predicted similar complex formations of AftA and LpqZ as AlphaFold2 (Supplementary Fig. 8). The results showed that the ligand remained capable of binding the pocket in close proximity to the conserved AftA (*M. marinum*) residues D108 and R112 in the presence of LpqZ. This suggests that, rather than preventing the binding of the galactan chain, LpqZ might play a positive regulatory role. These findings are consistent with our previous observations that LpqZ is required for proper arabinogalactan synthesis.

### FecB and LpqZ, together with AftB and AftA, are involved in linkage of arabinan chains of LAM

Additionally to its role in arabinogalactan synthesis ^37^, AftB is also responsible for the terminal β(1 → 2) linkage of an arabinosyl residue to LAM ^38^. To test if the mutation in *fecB* has an effect on the terminal linkage of LAM, we compared the effect of *aftB* gene silencing using CRISPRi in the *M. marinum* wild-type strain and the *fecB* mutant. Previously, Jankute *et al*. observed lower quantity of LAM and a shift towards higher molecular weight upon gene silencing of *aftB* in *M. smegmatis* using MAb F30-5 antibody^38^, which recognizes the terminal arabinan branches of LAM (ara-LAM) ^41^. Whole cell lysates of our wild-type *M. marinum* strain stained with F30-5 antibodies showed a slight shift in band pattern towards higher molecular weight upon *aftB* silencing (Fig. 4b), similar as in Jankute *et al*. Remarkably, the *fecB* mutant exhibited an even greater shift than the *aftB* silenced wild-type strain.

Additionally, both in the presence and absence of AftB, the shifted bands of the *fecB* mutant showed lower intensity than in the wild-type. This suggests that FecB, together with AftB, is indeed involved in the terminal linkage of arabinosyl to LAM. While the function of AftA has thus far only been linked to arabinogalactan synthesis ^34^, we investigated whether we could observe any effect on LAM. To explore this, we examined the effects of their depletion on ara-LAM, similarly to the experiments conducted for AftB and FecB. While silencing *aftA* alone in the wild-type strain did not produce a clear band shift, the additional deletion of *lpqZ* caused a distinct shift toward lower molecular weight. (Fig. 4c). These findings suggest that AftA and LpqZ may also play a role in modulating the structure of LAM. The expression of functional *lpqZ* and *fecB* under the control of the HSP60 promoter fully complemented the observed phenotypes (Supplementary Fig. 9).

To validate these findings, we extracted LAM from the cell envelope and assessed molecular weight changes and quantity in the mutants. The *lpqZ* mutant exhibited a slightly lower molecular weight LAM band, while the *fecB* mutant displayed a slightly higher molecular weight band on SDS page after Pro-Q Emerald staining (Fig. 4d). Furthermore, the *fecB* mutant produced significantly (p=0.0012) less LAM compared to the wild-type strain (Fig. 4e). These phenotypes were fully restored in the genetically complemented strains. In contrast, mutations in *fecB* and *lpqZ* did not significantly affect the size or quantity of the precursor LM (Fig. 4f).

### FecB and LpqZ are orphan SBPs that evolved novel functions in mycobacteria

Thus far, our findings indicate that the proteins produced by the orphan genes *fecB* and *lpqZ* have adapted new functions in mycobacteria. To further delve into this, we were interested in the evolutionary conservation of these proteins. To compare the structural conservation of FecB and LpqZ with other SBPs that still function in concert with ABC transporters, the ConSurf server was used for analyzing evolutionary conservation of amino acids ^42^. For each amino acid, the score of conservation was mapped to the predicted AlphaFold structures.

Interestingly, and in line with our findings, LpqZ showed high conservation in regions that are predicted to interact with AftA (Fig. 5a). The number of conserved residues was especially high towards the probable galactan binding cavity and the probable active site of AftA. In contrast, other SBPs like the *M.marinum* homologue ProX (25.17% aa identity) and *M.marinum* ModA showed most conserved amino acids around the middle part of the structure (Supplementary Fig. 10), which forms the substrate binding pocket ^43,44^. Similarly, FecB showed highly conserved amino acids in the interaction regions with AftB and MMAR_1667 (Fig. 5b). In contrast, the homologous SBPs FecB of *E.coli* and SirA (*S. aureus*) do not show amino acid conservation at these sites, but again rather around the center of the structure (Supplementary Fig. 11). Therefore, these evolutionary conservation profiles support our finding that FecB and LpqZ have functionally diverged from their original roles as SBPs.

**Figure 5:**
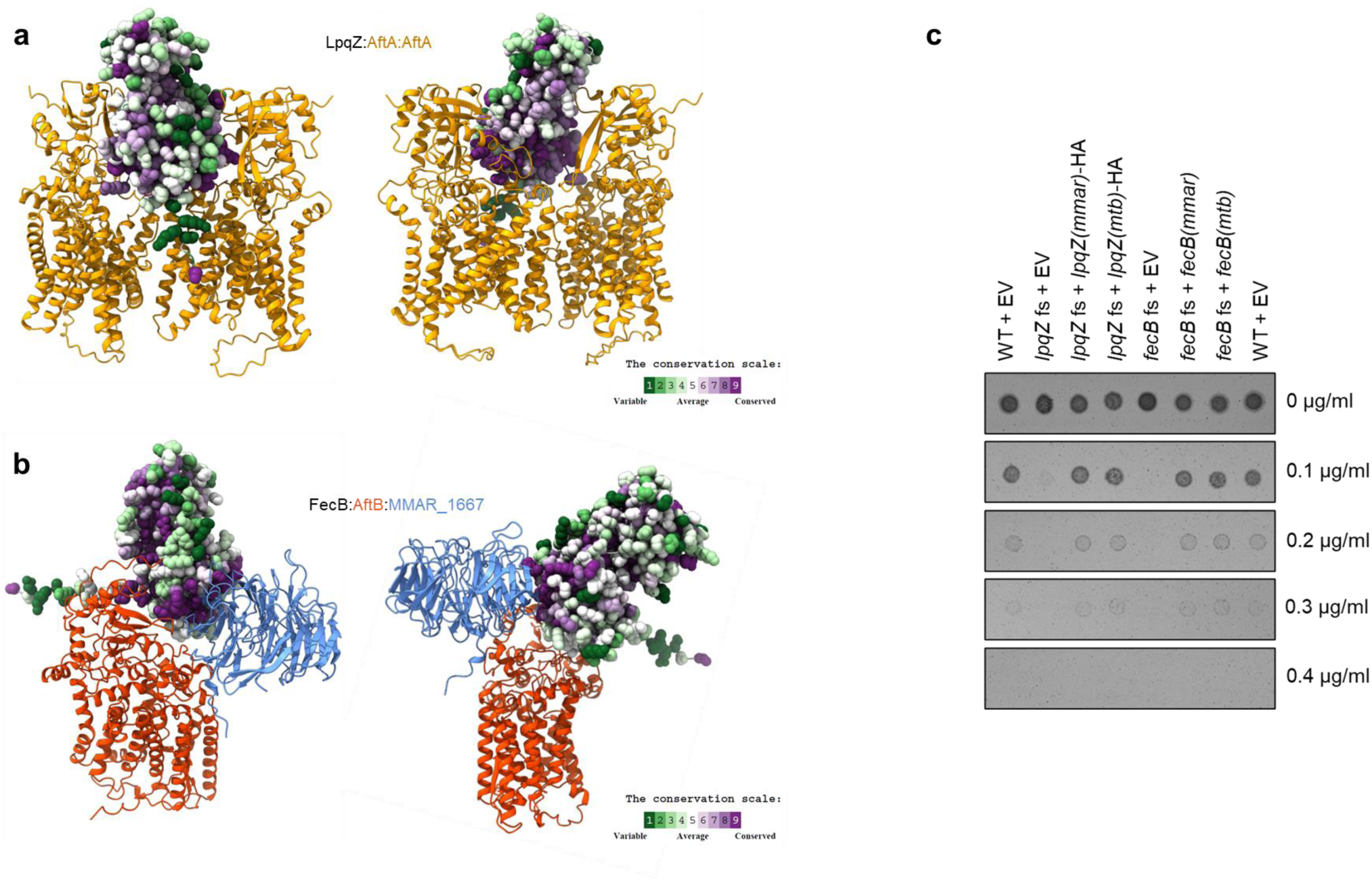
evolutionary conservation of LpqZ and FecB. (**a**): Conservation analysis of *M. marinum* LpqZ using the ConSurf server. LpqZ is depicted in atomic representation and amino acids are colored according to Consurf color code (green (variable), white (average), purple (conserved). Interaction with AftA dimer (yellow) is shown as predicted by AlphaFold2. (**b**): Conservation analysis of *M. marinum* FecB, performed as in (a). Interaction with AftB (light blue) and MMAR_1667 (red) is shown as predicted by AlphaFold2. (**c**) genetically complemented *M. marinum lpqZ* fs mutant *(lpqZ* fs + *lpqZ*-HA) and *fecB* fs mutant (*fecB* fs + *fecB*) with their respective *M. marinum* (*mmar*) and M. tuberculosis (*mtb*) genes were compared to wild-type (WT) and fs mutants containing the empty pSMT3 vector (EV) for susceptibility to different concentrations of rifampicin. Shown are representative pictures of two independent experiments (n=2).

To validate that the function of FecB and LpqZ is conserved in the major pathogen *M. tuberculosis*, we genetically complemented the *M. marinum* fs mutants with *M. tuberculosis fecB*(Mtb) and *lpqZ*(Mtb) respectively. The expressed Mtb genes fully complemented the antibiotic susceptibility phenotype of the mutants, suggesting that FecB and LpqZ play a similar role in *M. tuberculosis* (Fig. 5c).

## Discussion

Maybe the most important virulence factor of Mycobacteria is its protective cell envelope, which needs to be built, maintained and regulated by a variety of enzymes and accessory proteins ^8^. A key group of proteins involved in this process is the lipoproteins, some of which have demonstrated critical functional roles ^13,14,16,17^. We hypothesized that there are additional lipoproteins with key functions yet to be discovered.

In this study, we demonstrated through pooled susceptibility testing that a significant proportion of our lipoprotein library is essential for proper growth and intrinsic antibiotic resistance. Notably, *lpqZ* and *fecB* mutants exhibited heightened susceptibility to all tested antibiotics. Interestingly, both genes share homology with SBPs although they are not associated with an ABC transporter operon across all mycobacteria, suggesting that these orphan genes acquired novel functions. Co-IP and MS analyses identified AftA and AftB as interaction partners of LpqZ and FecB, respectively, implicating them in arabinogalactan and LAM synthesis. Indeed, both *lpqZ* and *fecB* mutants displayed alterations in LAM molecular size and arabinan-to-galactan ratio.

A hypersusceptible phenotype for a *fecB* mutant has previously been reported by Xu *et al*. through chemical genetic interaction profiling of a *M. tuberculosis* transposon mutant library, showing cross-sensitivity to all tested antibiotics^26^. These findings suggest that FecB primarily contributes to cell envelope integrity rather than iron metabolism, despite its homology to *E.coli* FecB and SirA. While Xu *et al*. did not observe growth differences for the *fecB* mutant under low-iron conditions, affinity binding experiments by Miranda *et al*. proposed that FecB possesses heme- and mycobactin-binding properties^32^. However, our observations of an overall disturbed cell envelope integrity align more closely with the findings of Xu *et al*., supporting a novel function of FecB. Recent studies emphasize the importance of LM and LAM in maintaining cell envelope integrity and intrinsic resistance to vancomycin and β-lactam antibiotics ^6,7^, further substantiating our hypothesis.

Interestingly, MMAR_1667 appears critical within the trimeric complex, as our frameshift mutant displayed a strong growth reduction similar to that of the *fecB* mutant and wild-type with repressed *aftB* expression. Notably, the *M. smegmatis* orthologue of MMAR_1667 (MSMEG_1053) was identified in the above-mentioned study by Miranda *et al*. through a co- IP of FecB-FLAG, ranking second in intensity after FecB itself, but has not further been investigated as a potential interaction partner. MMAR_1667 shares structural homology with the WD40 protein BamB (17.4% amino acid sequence identity), an auxiliary lipoprotein of the Bam complex, which facilitates outer membrane protein localization but whose precise functions remains unclear ^45^. Recent studies suggest that BamB clusters multiple Bam complexes into so-called precincts by cross-linking with neighboring BamA subunits ^46^.

Interestingly, MMAR_1667 harbors a predicted lipoprotein signal peptide (SignalP6.0), whereas its orthologue in *M. tuberculosis,* Rv3035, lacks a conventional lipobox, except for a glycine residue preceding the putative conserved cysteine. This raises the possibility that MMAR_1667 is a lipoprotein that was overlooked in our initial screen due to our selection criteria favoring conserved *M. tuberculosis* lipoproteins.

HA-tagged FecB was unable to rescue the *fecB* mutant phenotype. AlphaFold predictions suggest that the HA tag’s positioning at the center of the trimeric complex may cause steric hindrance and prevent proper complex formation or function. While we cannot exclude a secondary role for FecB in iron metabolism, our data strongly indicate that its primary function underlies the growth defect and antibiotic sensitivity phenotypes observed in *fecB* mutants.

LpqZ shares homology with ProX and other amino acid-binding proteins associated with ABC transporters. Notably, ProX is part of a functional ABC transport operon ProXW within the same genome, suggesting that *lpqZ* may arose from duplication events and subsequently diverged to adopt novel functions. Interestingly, LpqZ shares structural similarity with the lipoprotein MltF, a transglycosylase from *Pseudomonas aeruginosa*, which consists of both a catalytic and a non-catalytic regulatory module ^47^. Like LpqZ, the regulatory module of MltF resembles many periplasmic amino acid-binding proteins in ABC transporters ^47^. This homology suggests that not only LpqZ, but also other SBP-like proteins, may perform diverse cell envelope roles beyond canonical substrate transport.

Mycobacterial AraTs, a family of glycosyltransferases ^48^ mediate AG and LAM synthesis. These transmembrane proteins utilize DPA as an arabinosyl donor to elongate and branch arabinan domains of the galactan or mannan core ^49^. While AftA primes AG synthesis ^34^, the enzyme priming the mannan chain of LAM remains unidentified^50^. In AG synthesis, arabinan extension has been proposed to be done by EmbA and EmbB, while for LAM synthesis, this linkage might be done by EmbC ^49^. Both in AG and LAM synthesis, AftC is believed to be responsible for the internal α-(1→3) branching of arabinan ^51,52^, while AftD is believed to add the final few α-linked residues to the branches ^53,54^ before capping via terminal β(1 → 2) linkage is done by AftB ^37,38^.

Except for AftC, cryo-EM and crystal structures have been solved for all AraTs ^35,55–57^. AftD consists of a transmembrane domain with typical glycosyltransferase C (GT-C) fold and a periplasmatic soluble domain that consists of three carbohydrate binding modules (CBMs), which are thought to be responsible for the recognition and binding of complex arabinan chains ^56^. Interestingly, AftA has a GT-C fold core ^35^, but is lacking CBM domains, suggesting that LpqZ may compensate for this function (Supplementary Fig. 12). Similarly, FecB and MMAR_1667 may serve a CBM-like role for AftB, which shares structural features with AftA^55^.

Surprisingly, AftA depletion in the LpqZ-deficient strain resulted in lower molecular weight LAM molecules, suggesting a previously unrecognized role of these proteins in LAM synthesis. As we suspected, the presence of LpqZ is also important for the linkage of arabinan to arabinogalactan, as seen by a decreased arabinan to galactan ratio. Molecular docking of a synthetic galactan chain, which has also been used by Gong *et al*.^35^, showed that the galactan chain would still be able to bind to AftA in the presence of its binding partner LpqZ.

In contrast to AftA and LpqZ, AftB depletion in the FecB-deficient strain led to larger apparent molecular weight LAM molecules, potentially due to extended arabinan branches lacking terminal β(1 → 2)- linked arabinosyl units. This alteration may reduce LAM synthesis efficiency, which could explain our observations of lower LAM levels.

Even though LpqZ and FecB are not essential like their partner proteins AftA and AftB, their deletion caused growth defects and increased antibiotic susceptibility, in line with their proposed roles in AG and LAM synthesis. Our findings suggest that LpqZ and FecB may act as positive regulators of these pathways, potentially performing functions analogous to the CBMs of AftD. However, the exact molecular mechanisms underlying their involvement remain to be fully elucidated. Although LpqZ and FecB resemble classical SBPs, their lack of associated cytoplasmic transport systems and apparent adoption of novel functions highlight the need to revisit the assumption that structural similarity implies functional conservation.

The ability of *M. tuberculosis* LpqZ and FecB to rescue the phenotype of *M. marinum* mutants suggests that their functions are conserved in this major pathogen. These results present opportunities to explore LpqZ and FecB as potential secondary targets in future therapeutic strategies, particularly those aimed at sensitizing mycobacteria to existing antibiotics.

## Methods

### Bacterial strains and culture conditions

All bacterial strains that were used in this study are listed in supplementary table S3. *Mycobacterium marinum* M wild-type strain and derivative mutant strains were cultured at 30°C on Middlebrook 7H10 (Difco) plates supplemented with 10% OADC (Difco), or shaking at 90 RPM in 7H9 medium (Difco), supplemented with 10% ADC (Difco) and 0.02% tyloxapol. When required, appropriate antibiotics were added (50µg/ml hygromycin (Roche), 50µg/ml kanamycin (Sigma) or 30µg/ml streptomycin (Sigma)). Molecular cloning was performed using *E. coli* DH5α, which was grown on LB agar and LB medium at 37°C and 200RPM. Antibiotics were used at the same concentrations as stated above.

Electrocompetent *M. marinum* strains were prepared by growing cultures to OD_600_ of 0.8-1.5. Cultures were incubated for 16h with 1.5% glycine until harvesting and washing three times with cold 10% glycerol. Bacteria were resuspended at a concentration of approximately 20 OD/ml in 10% glycerol and preserved at -80°C.

### Plasmids

All generated plasmids were verified using Sanger Sequencing (Macrogen) and are provided in supplementary table S4. If not stated elsewise, amplification of genes of interest was done from template genomic DNA from *M. marinum* M or *M. tuberculosis* H37Rv strain using Phusion High Fidelity DNA polymerase (ThermoFisher) and specific primers (supplementary table S5), while digestion of plasmids was done using restriction enzymes provided by New England Biolabs (NEB). Genes of interest were cloned into the respective digested vectors using In-Fusion cloning (Takara Bio). For cloning into the pSMT3 plasmid (hyg^R^), HA-tagged genes were inserted downstream of the HSP60 promoter using the NheI and BamHI restriction sites, following the same strategy used for pSMT3-lpqZ-HA (Addgene 240159). In contrast, untagged genes were introduced using NheI and XbaI sites, as done for pSMT3- fecB (Addgene 240158). For expression from the atc-inducible p766’ promoter from pLJR962 (kana^R^, Addgene 115162), restriction sites ClaI and NotI were used. For cloning of pML1357 plasmids, the hygromycin cassette was removed from pML1357-mmar_0407 ^58^ with HpaI and NsiI restriction and exchanged for the amplified kanamycin cassette from pLJR962. For subsequent introduction of MMAR_1667-Strep, the plasmid was cut with NheI and MluI.

The cloning of pLJR965 (kana^R^, Addgene 115163) for the purpose of CRISPRi gene silencing was done as in ^59^. Similarly, the construction of pCRISPRx-Sth1-Cas9-L5 (kana^R^, Addgene 140993) containing sgRNAs for the respective target genes (Supplementary table S5) was done as described in ^23^. Targets were predicted using CHOPCHOP ^60^ with PAM sequences NNAGAA, NNGGAA and NNGGAG.

### Construction of CRISPR/Cas9 mutants

CRISPR/Cas9 frameshift mutants were constructed as described in ^23^. Briefly, electrocompetent *M. marinum* were electroporated at a single pulse of 2.5kV, 25µF, 720Ω with 1µg of pCRISPRx-Sth1-Cas9-L5 (Addgene 140993) plasmid harboring the respective gRNA(s). After 6h recovery in 7H9 medium, cells were plated on 7H10 plates containing kanamycin and 100ng/ml ATc (IBA Lifesciences). After 7-10 days incubation, single colonies were picked for PCR amplification of the targeted locus using Phusion High Fidelity DNA polymerase (ThermoFisher) with specific primers (supplementary table S5) and Sanger sequencing (Macrogen). For the replacement of pCRISPRx-Sth1-Cas9-L5, mutants were electroporated with pTdTomato-L5 (Addgene 140994) and selected on 7H10 plates containing streptomycin.

### Pooled antibiotic susceptibility testing

*M. marinum* CRISPR/Cas9 fs mutants and *M. marinum* wild-type pCRISPRx-Sth1-empty were grown to mid-log phase (approx. OD 1.2-1.6), washed in PBS and filtered (5µM Millex Millipore). Aliquots of OD10 were prepared and frozen in 10% glycerol in water at -80°C until equal amounts of the mutants and controls were pooled together. For growth on 7H10 plates containing the respective antibiotics, 1x10^6 CFU of the pooled bacteria stock were spread on plate and grown at 30°C until colonies were visible (7-14 days). To determine the appropriate antibiotic concentrations, the pool was grown on plates with varying antibiotic levels, and we selected those where bacterial growth was evident but notably reduced. For harvesting of DNA, all colonies were scraped from the plate using PBS and a spatula, taken up in 250µl resuspension solution (GeneJET Plasmid miniprep kit) and lysed with 0.1mm Zirconia beads in the bead beater for 3min. Afterwards, the lysed bacteria were heated at 98C for 10min and briefly centrifuged to collect large debris. 200µl were used for DNA extraction with a DNeasy blood and tissue kit (Qiagen). Isolation of DNA from the initial input pool was done in a similar way by lysing 150µl of the initial bacterial pool. For PCR amplification of the specific sgRNA locus on the individual CRISPR plasmids in the mutants, primers with a barcode (ACACAC; GTCTCT; ACATGT; ACTAGC; ATGATA; ACGACG; ATATCG; CTCGCA) for each pool (Fw:XXXXXXGCTCAGTCGAAAGACTGGGC, Rv: AGAGACAGGGTGTTGATTTCGGCATG) were used. PCR was performed in duplicate for each pool and the bands of interest were separated on an agarose gel and isolated together in one tube using a GeneJET gel extraction kit. The pooled DNA was sequenced by NOVOGENE, UK (PCR-product Illumina Sequencing PE150). 2GB raw data was demultiplexed in CLC workbench with the primer barcodes of each pool. Using 2FAST2Q, sgRNA reads were extracted and counted for each of the pools and gene IDs were assigned to each sgRNA. For data normalization and statistical testing, drugZ ^61^ was used. The half_window_size was adjusted accordingly for smaller gene populations to a value of 15. For analysis, technical replicates were averaged, whereas biological replicates (n=2) were treated as different groups. To assess the general growth of the mutants, the read counts of the 7H10 plate without antibiotics was compared to the input pool. For testing the effect of antibiotics, the read counts from the respective antibiotic plate was compared to the 7H10 plate without antibiotics. Log2-fold change data was plotted for each condition and p-values were indicated as they were derived from drugZ.

### Bacterial growth assay on solid medium

For assessing general bacterial growth on solid medium, bacterial cultures were grown to mi- log phase (approx. OD 1.2-1.6), washed twice in PBS and diluted to an OD of 0.1. Subsequently, 2µl of ten-fold serial dilutions were spotted on 7H10 agar plates and incubated at 30°C for 7 days. For assessing antibiotic susceptibility, OD 0.1 dilutions were directly spotted on 7H10 agar plates containing different concentrations of the respective antibiotic. For assessing growth upon gene silencing, strains containing pLJR965 with target sgRNAs were grown on selective 7H10 agar with and without 200ng/ml anhydrotetracycline (ATc).

### Ethidium bromide permeability assay

Mycobacterial cultures were grown to mid-log phase (approx. OD 1.2-1.6), harvested and washed twice in PBS. OD was adjusted to 0.8 and technical quadruplicates of 200µl bacterial suspension were prepared in PBS containing 5µg/ml ethidium bromide. Fluorescence (Exc. 360nm/ Em. 590nm) was measured at 30°C using a plate reader (Biotek Synergy H1) every 5min over a time course of 4h.

### Protein purification of HA-tagged and Strep-tagged protein from the cell envelope

Mycobacterial cultures were grown to mid-log phase (approx. OD 1.2-1.6), harvested at 5000 RPM for 5min and washed in PBS. When needed, protein expression was induced with 200ng/ml ATc for 48h. Pellets were resuspended in 10% glycerol in PBS at a concentration of 50 OD/ml and passed three times through a One-shot cell disrupter (Constant Systems) at 0.83kbar. Unbroken cells were removed by centrifuging twice at 5000RPM for 5min and 4°C. The supernatant was centrifuged at 150000xg for 1.5h at 4°C. The pellet was resuspended in a tenth of the initial volume in 10% glycerol in PBS. Protein concentrations were determined by BCA and adjusted to 1mg/ml in 1ml 10% glycerol in PBS. For solubilization of membrane proteins, 0.25% n-dodecyl-ß-D-maltoside (DDM) was added and incubated for 1h head over head rotation at 4°C. After centrifugation at 100000xg for 30min at 4°C, the resulting soluble fractions were used for purification of HA-tagged proteins using a µMACS™ HA isolation kit (Miltenyi Biotec) and a magnet separator according to the manufacturer. For washing, 10% glycerol in PBS with 0.03% DDM was used and the first two elution steps were done using 2mg/ml HA peptide in washing buffer, whereas the last elution was done as described by the manufacturer. For strep-tag purification, membrane fractions were adjusted to 4.5mg/ml in 10% glycerol PBS and solubilized. Naturally biotinylated proteins were blocked with 0.5mg/ml avidin for 10min before centrifugation. The resulting soluble fractions were incubated with StrepTactin resin (IBA Lifesciences) under rotation at 4°C for 1h. Beads were washed five times with 10% glycerol in PBS and eluted three times with elution buffer (50mM TrisHCl pH8, 300mM NaCl, 5% glycerol, 10mM desthiobiotin). To check for purification efficiency, samples were taken after each step, boiled in SDS loading buffer for 10min at 98°C and examined by SDS page. For each purification sample, all three elutions were pooled together for SDS-page and mass spectrometry as described below.

### Mass spectrometry

For identification of protein interaction partners, elution samples from Strep-tagged or HA-tagged protein purification were analyzed by LC-MS/MS as described in ^62^. Briefly, elution samples were boiled in the presence of SDS buffer at 98°C for 10min and loaded on a SDS-page gels. Gels were stained with Commasie Brilliant Blue G-250 and the total lane of each sample was excised from the gel for subsequent in-gel digestion. Peptides were separated by the Ultimate 3000 nanoLC-MS/MS system (Dionex LC-Packings, Amsterdam, The Netherlands) and proteins were identified by searching against the uniprot database entry of *M. marinum* BAA-535_230714. For calculation of relative LFQ intensity (fold change), the average LFQ of tagged samples was divided by the average LFQ of control samples, whereby the minimum LFQ value of all replicates was added as a constant to both averages to circumvent division by 0. In the case of the MMAR_1667-Strep co-IP data set, we observed one outlier (MMAR_0418), which we considered as biologically irrelevant due to its appearance with high values in a parallel experiment. Therefore, it was excluded from the figure.

### AlphaFold complex predictions and evolutionary conservation analysis

Complex predictions were done using AlphaFold2 from the Google ColabFold v1.5.5 using MMseqs2 for multimer predictions. If not indicated elsewhere, amino acid sequences of *M. marinum* were used for predictions. Amino acid sequences were retrieved from Mycobrowser and predicted signal peptides were removed for all structural predictions. Default settings were used for the alphafold2_multimer_v2 model type. ChimeraX was used to color by chain and by b-factor attribute.

Analysis of evolutionary conservation was done on the basis of predicted AlphaFold structures using the ConSurf server ^42^ with a set minimum percentage of identity of 40% and a maximum of 75%.

### Molecular docking

Protein–ligand docking was performed using HADDOCK version 2.4 ^39,40^. A synthetic galactan ligand comprising eight galactofuranosyl residues was used. This ligand represents the galactan portion of the octyl-galactoside LG8, as described by Angala et al. ^63^, and is the form used for docking in Gong *et al*. ^35^. The ligand was constructed using BIOVIADraw2020. The three-dimensional structure was generated using Open Babel version 3.1.1 ^64^ with MMFF94-based energy minimization, and a PDB file was prepared via Open Babel GUI. To account for ligand flexibility, an ensemble of 2,000 conformers was generated using RDKit (https://www.rdkit.org) version 2020.09.3. This ensemble was used as input for HADDOCK, which uses restraints to guide the docking. In our case we used random restraints from surface accessible molecules. During the initial rigid-body energy minimization step, the number of generated structures was increased from the default 1,000 to 40,000, resulting in 20 docking models per ligand conformer. The top 200 structures, ranked by HADDOCK score, were subjected to a semi-flexible simulated annealing protocol in torsion angle space, allowing for gradual introduction of flexibility at the protein–ligand interface. The final flexible refinement step in explicit solvent was not performed. A total of 200 docking models were analyzed, and the final complex was selected based on two criteria: (1) the ability to form a hydrogen bond between the C5-linked hydroxyl group of the galactan chain and the catalytic residue AftA-D128, and (2) a favorable HADDOCK score. This selection approach followed a similar strategy to that described by ^35^.

### Lipoarabinomannan detection

Mycobacterial cultures were grown grown to an OD₆₀₀ of approx. 0.4 in 7H9 containing 10% ADC in the absence of detergents. Cultures were split in half and one of the cultures was induced with 200ng/ml atc for gene knock down purposes. After 3 days, cultures were harvested at 5000 RPM for 5min and washed in PBS. Pellets were resuspended in 10% glycerol in PBS at a concentration of 50 OD/ml and passed three times through a One-shot cell disrupter (Constant Systems) at 0.83kbar. Unbroken cells were removed by centrifuging twice at 5000RPM for 5min and 4°C. Protein concentrations were determined by BCA and normalized. Samples were boiled in SDS sample buffer containing DTT for 10 min and sonicated in a sonifier (BRANSON sonifier 250) for 1min before loading on a 10% SDS page.

### Protein detection

For protein detection by Western blot, proteins were transferred to a nitrocellulose membrane and visualized by standard immunodetection techniques. For detection of HA-tagged, FLAG- tagged and Strep-tagged proteins, anti-HA (HA.11, Covance), anti-FLAG (M2 monoclonal antibody produced in mouse, Sigma) and anti-Strep (Novusbio, NBP2-41073) antibodies were used. For detection of terminal arabinan chains of lipoarabinomannan, anti-arabinan LAM F30-5 antibody ^41^ was used. F30-5 was provided by Arend Kolk, Royal Tropical Institute, Amsterdam, the Netherlands. Additionally, for detection of EccC5 we used anti- EccC5 antibodies ^65^.

### Purification and analysis of mycobacterial glycolipids

Mycobacterial glycolipids, including lipomannan (LM) and lipoarabinomannan (LAM), were isolated and analyzed as previously described ^3^. Briefly, *M. marinum* cultures (1 L) were grown to an optical density at 600 nm (OD_₆₀₀_) of 0.8. Cells were harvested by centrifugation at 6,000 × g for 10 min and resuspended in 20 mL phosphate-buffered saline (PBS) containing 0.1% (v/v) Tween-80. Cell lysis was achieved by bead-beating, and the lysate was transferred to Teflon-capped glass tubes. For glycolipid extraction, an equal volume of water-saturated phenol was added to the lysate, and the mixture was incubated at 85 °C for 2 h with intermittent mixing. The aqueous phase was separated by centrifugation at 4,000 × g for 10 min and collected. This phenol extraction was repeated twice to maximize recovery.

Combined aqueous phases were dialyzed overnight against tap water, followed by a 2 h dialysis against double-distilled water (ddH₂O), and subsequently lyophilized. Lyophilized glycolipids were resuspended in ddH₂O and normalized to 1 mg wet cell mass per mL. For visualization, 20 µL of each sample was mixed with 5 µL SDS-PAGE loading buffer and separated on a 4–20% precast polyacrylamide gel. Following electrophoresis, gels were fixed in 50% methanol, 5% acetic acid for 30 min (twice), then washed three times in 3% acetic acid. Glycolipids were oxidized in periodic acid for 30 min and washed again with 3% acetic acid. Staining was performed using the Pro-Q Emerald 300 Glycoprotein Gel Stain Kit (Thermo Fisher Scientific). Gels were imaged using a Bio-Rad Gel Doc system under 300 nm UV light, and densitometry analysis was carried out using ImageLab 6.1 (Bio-Rad Laboratories).

### Arabinogalactan Extraction

*M. marinum* strains were cultured in 100 mL Middlebrook 7H9 medium (BD Difco), supplemented with OADC enrichment (BD Biosciences) and 0.05% (v/v) Tween 80, and incubated at 30 °C with shaking until reaching an OD₆₀₀ of 0.8. Cells were harvested by centrifugation at 4,000 × g for 30 min at 4 °C, washed once with phosphate-buffered saline (PBS) containing 0.05% (v/v) Tween 80, and centrifuged under the same conditions. Cell pellets derived from 100 mL cultures were resuspended in 1 mL PBS-Tween, followed by dilution in 15 mL PBS. Cells were lysed using a FastPrep-24 Bead Beating System (MP Biomedicals) with two 45 s pulses, separated by a 3 min incubation on ice. Lysates were transferred to 1.5 mL tubes and treated with 4% (w/v) sodium dodecyl sulfate (SDS) at 110 °C for 15 min. Samples were centrifuged at 15,000 × g for 1 h at 25 °C, and supernatants discarded. Pellets were washed eight times with warm water (25 °C) using identical centrifugation parameters to remove residual SDS. To remove mycolic acids, pellets were incubated in 6 mL of 0.5% (w/v) potassium hydroxide in methanol at 37 °C with constant stirring for 4 days. Samples were chilled, centrifuged at 15,000 × g for 1 h at 4 °C, and supernatants discarded. Pellets were washed once with cold methanol, centrifuged as above, and residual methanol allowed to evaporate. Pellets were then resuspended in cold diethyl ether, transferred to glass tubes, and centrifuged at 3,000 × g for 20 min at 4 °C. Subsequently, pellets were resuspended in 60 mL PBS containing 15 µg/mL proteinase K (Sigma-Aldrich) and incubated overnight at 55 °C with stirring. After digestion, samples were centrifuged at 15,000 × g for 30 min at 25 °C, and supernatants were discarded. Pellets were washed once with water under the same conditions, then resuspended in 6 mL of 0.2 M sulfuric acid, transferred to round-bottom flasks, and heated at 85 °C with stirring.

Neutralization was performed by the gradual addition of calcium carbonate under pH monitoring. After neutralization, samples were centrifuged at 15,000 × g for 30 min at 25 °C. Supernatants containing arabinogalactan were retained, and pellets were washed twice with water and centrifuged as above. All supernatants were pooled. The combined extract was dialyzed overnight at 4 °C in continuously flowing water using 3.5 kDa molecular weight cutoff (MWCO) dialysis tubing (Spectra/Por). Dialyzed material was frozen and lyophilized. Pre-weighed containers were used to determine arabinogalactan yield. Lyophilized material was hydrolyzed in 1 mL of 300 mM hydrochloric acid at 37 °C for 1 h, neutralized with 1 M sodium hydroxide, and evaporated to dryness in pre-weighed Eppendorf tubes using a SpeedVac centrifugal evaporator.

### HPAEC analysis of acid-hydrolyzed arabinogalactan

Hydrolyzed samples were analyzed using a Dionex ICS-6000 high-performance anion-exchange chromatography (HPAEC) system equipped with a CarboPac PA1 analytical column and guard column (Thermo Fisher Scientific). Monosaccharide separation was performed at 1 mL/min with an isocratic elution of 100 mM NaOH over 20 min, followed by a 10 min wash with 500 mM NaOH. Detection was carried out using pulsed amperometric detection (PAD) with a gold working electrode and a palladium-hydrogen reference electrode, operated under the standard Carbohydrate Quad waveform at a frequency of 2 Hz. Quantification and retention time comparisons were performed using 50 µM aqueous solutions of D-arabinose and D-galactose standards (Sigma-Aldrich). Chromatographic data were processed using Chromeleon™ Chromatography Management Software (v6.8, Thermo Fisher Scientific).

## Statistical analysis

Graphpad Prism 10 was used for preparation of all plots and statistical analysis. DrugZ test for analysis of pooled barcoded experiments was carried out as mentioned above.

## Acknowledgements

This work was supported by the ERC Advanced grant 832721, NWO XL grant XL21.006, the NWO grant OCENW.KLEIN.439. and ZonMW grant 09120012010040. We thank Esther Keizer (Vrije Universiteit Amsterdam) for kindly providing the pLJR965-*aftA* and pLJR965- *aftB* plasmids used in this study. We are also grateful to Cor Lieftink (Netherlands Cancer Institute) and Traver Hart (University of Texas) for their helpful advice on selecting appropriate statistical tests and parameters for DrugZ analysis in our dataset.

## Author contributions

Robin Lissner: investigation, writing – original draft, writing – review and editing, conceptualization, methodology, validation, formal analysis, visualization

Aaron Franklin: investigation, validation

Samuel T. Benedict: investigation, validation

Patrick J. Moynihan: methodology, resources

Vicky Charitou: software, formal analysis

Alexander Speer: methodology

Jaco Knol: investigation, resources

Connie Jimenez: resources

Sergey Nejentsev: conceptualization, writing – review and editing, funding acquisition

Coen Kuijl: conceptualization, supervision, writing - review and editing

Wilbert Bitter: conceptualization, supervision, writing – review and editing, funding acquisition

## Competing interests statement

The authors declare no competing interests.

## Notes

### Competing Interest Statement

The authors have declared no competing interest.

