## supplementary figures and tables for "A new role for lipoproteins LpqZ and FecB in orchestrating mycobacterial cell envelope biogenesis"

**Supplementary data**

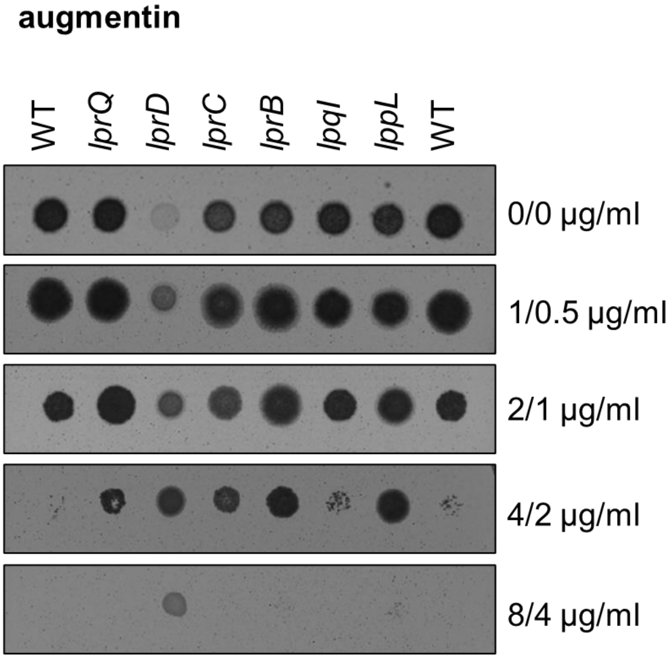


**Supplementary Figure 1: augmentin susceptibility for mutants with increased resistance**. lipoprotein CRISPR/Cas9 fs mutants and *M. marinum* wild-type strain (WT pCRISPRx-Sth1-Cas9-L5 empty) were spotted on 7H10 agar plates containing different concentrations of augmentin as in Figure 2c.

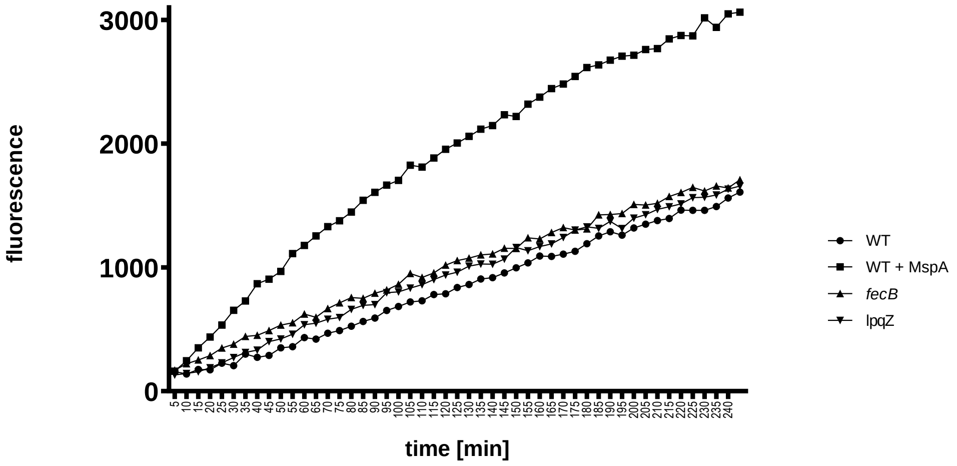


**Supplementary Figure 2: cell envelope permeability of lpqZ and fecB mutants.** *M. marinum* wild-type (WT pCRISPRx-Sth1-Cas9-L5 empty), a positive control (WT overexpressing MspA porin from *M. smegmatis*) and *fecB* and *lpqZ* CRISPR/Cas9 fs mutants were incubated with 5µg/ml ethidium bromide and uptake was monitored over a time course of 4h. Fluorescence was measured every 5min at Exc. 360nm and Em. 590nm. Data shows the mean of technical quadruplicates and is representative of two independent experiments.

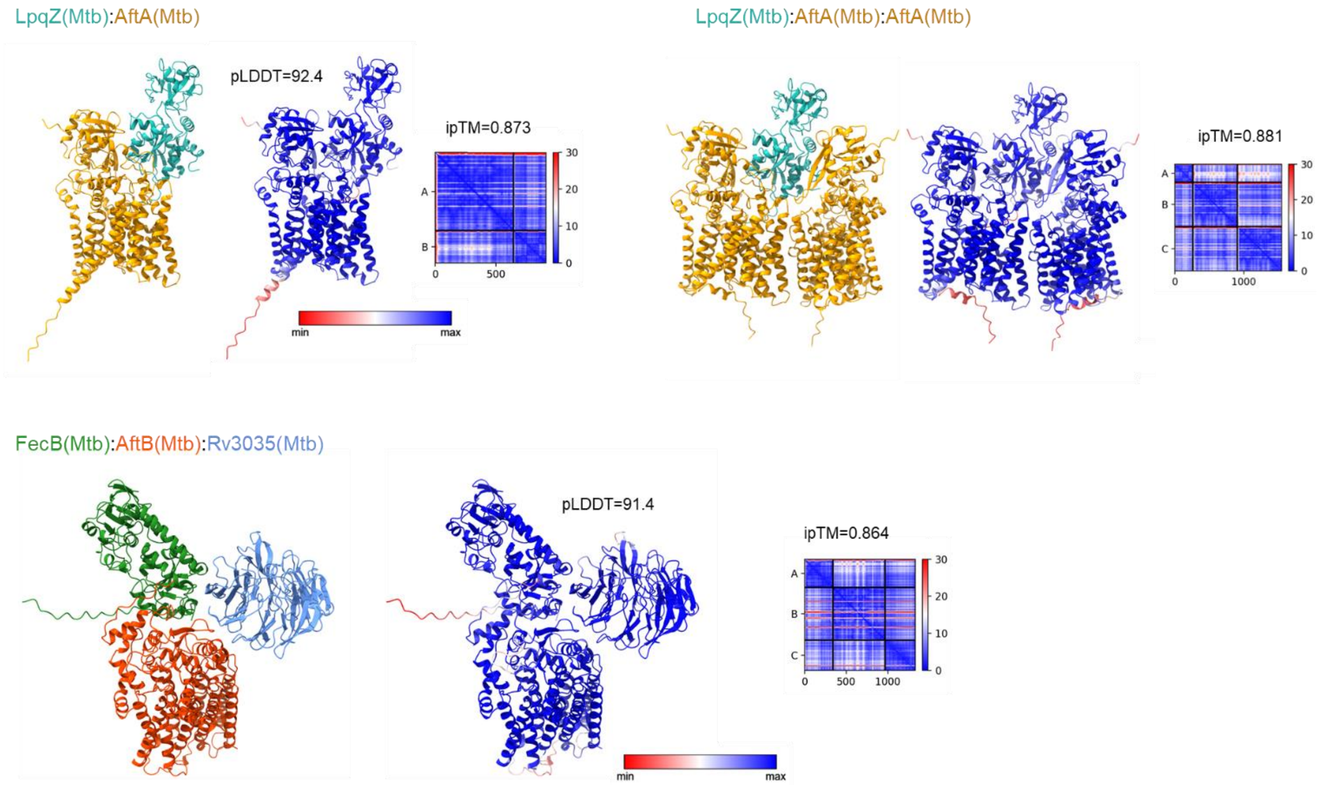


**Supplementary Figure 3: AlphaFold complex prediction of the putative interaction partners of LpqZ and FecB in *M. tuberculosis***. Shown are the complexes colored by chain (left), colored by pLDDT (middle) and PAE plot with ipTM values (right) of LpqZ and FecB with their respective interaction partners as in Figure 3c.

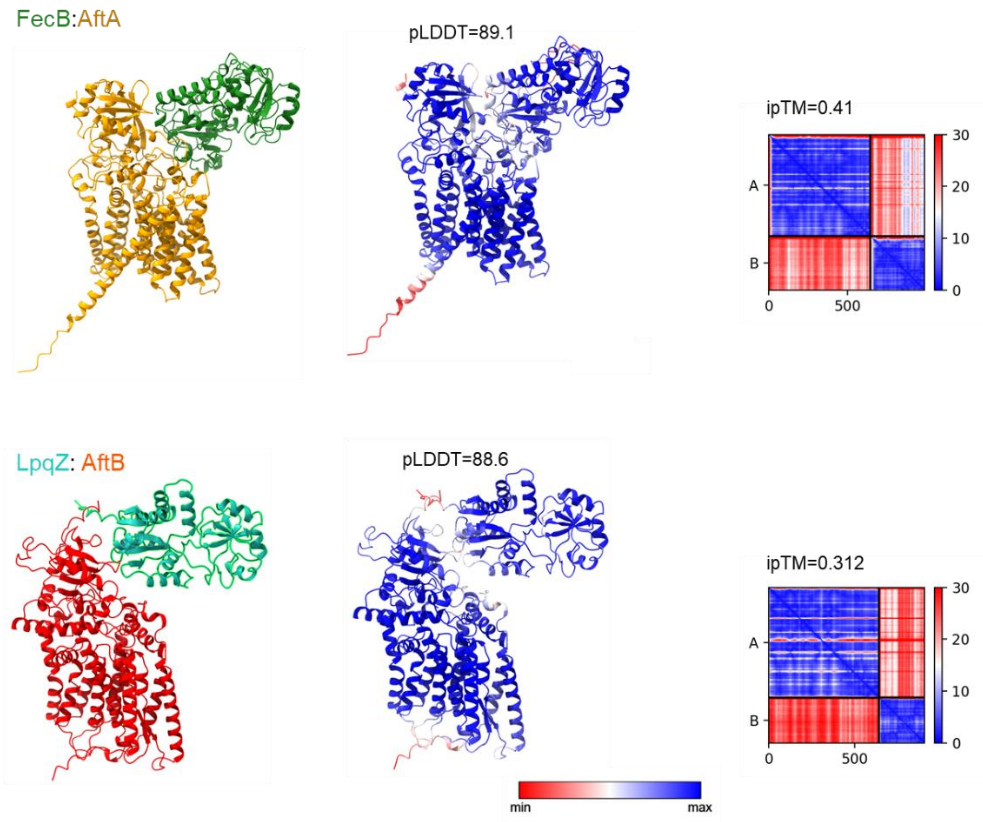


**Supplementary Figure 4: AlphaFold complex predictions of swapped interaction partners.** *M. marinum* LpqZ and FecB were predicted with AftB and AftA, respectively as a negative control of ipTM scores. Complexes are colored by chain (left), colored by pLDDT (middle) and PAE plot with ipTM values (right).

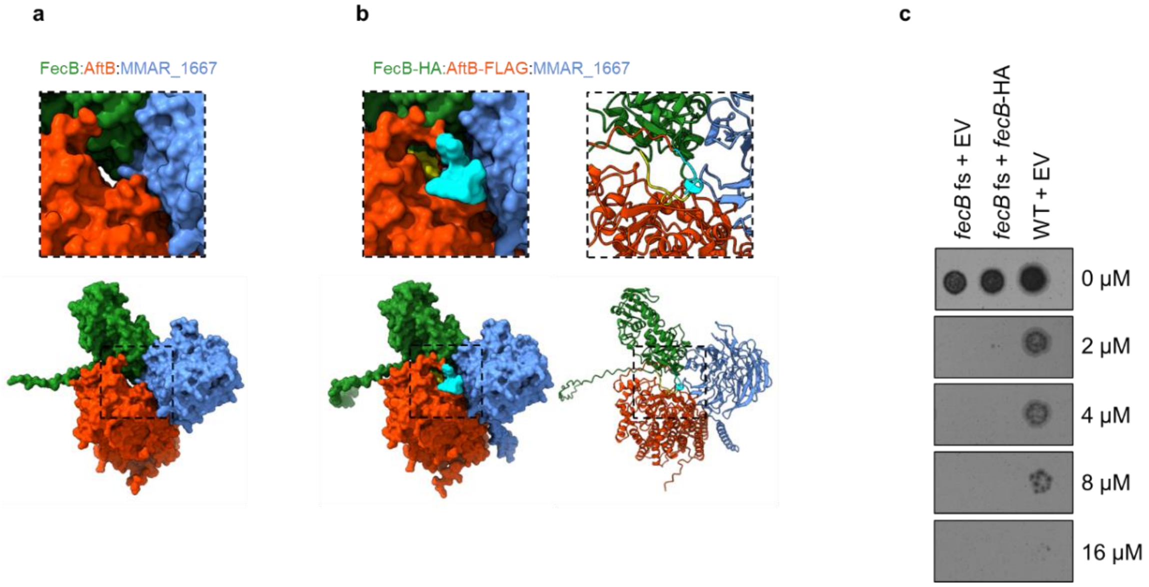


**Supplementary Figure 5: tagged FecB and AftB proteins might interfere with the complex formation** (**a**): AlphaFold prediction of *M. marinum* FecB:AftB:MMAR_1667 shows the center of the trimeric complex. (**b**): AlphaFold prediction of FecB-HA:AftB-FLAG:MMAR_1667. c-terminal HA-tag of FecB is labeled in yellow, c-terminal FLAG tag of AftB is labeled in cyan, localized at the center of the trimeric complex (**c**): expression of fecB-HA does not complement the *fecB* mutant phenotype. *M. marinum* WT strain, *fecB* fs mutant containing the empty pSMT3 vector (EV) and *fecB*-HA complementation strains were spotted on 7H10 plates containing different concentrations of vancomycin.

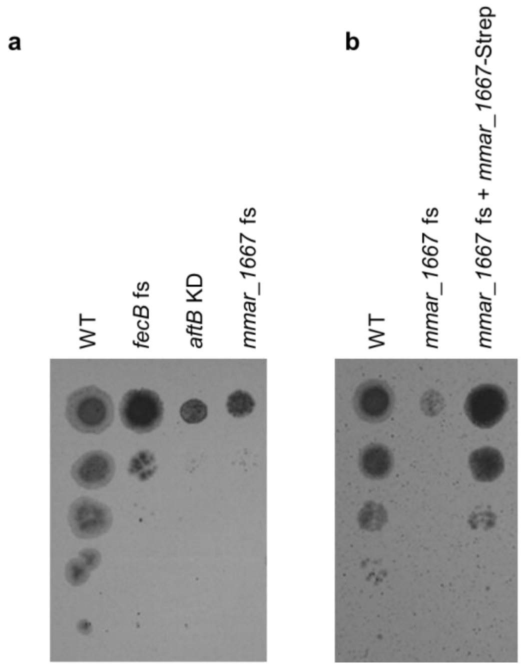


**Supplementary Figure 6: growth depletion of the *mmar_1667* mutant.** (**a**): for comparison of growth reduction, wild-type (*M. marinum* WT pCRISPRx-Sth1-Cas9 L5 empty), *M. marinum* CRISPR/Cas9 frameshift mutants *fecB*, *mmar_1667* and CRISPRi knockdown of *aftB* were spotted in 10-fold serial dilutions on 7H10 kanamycin plates containing 200ng/ml anhydrotetracycline. (**b**) Similar as in (a), the WT strain and the *mmar_1667* mutant were spotted along with the complemented mutant strain, expressing *mmar_1667*-Strep

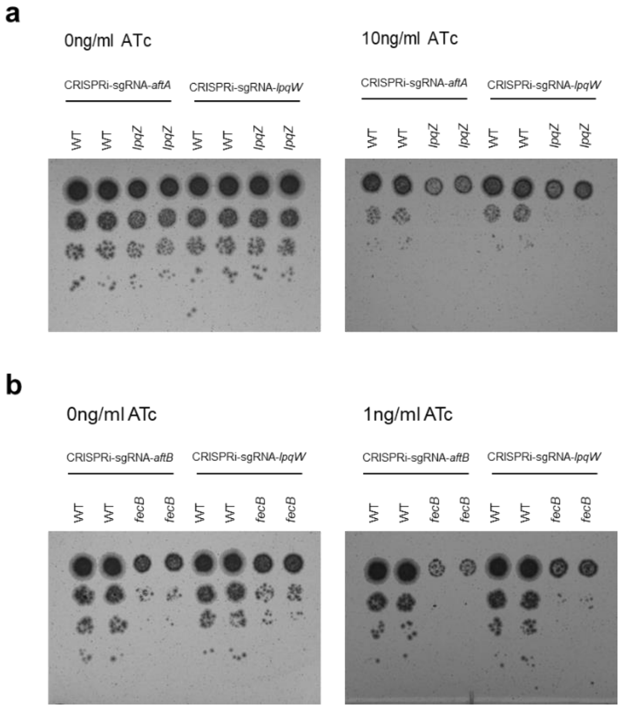


**Supplementary Figure 7: *aftA* and *aftB* gene silencing in *lpqZ* and *fecB* mutants. (a)** *M. marinum* wild-type and *lpqZ* fs mutant containing pLJR965-sgRNA-*aftA* and pLJR965-sgRNA-*lpqW* were spotted in 10-fold serial dilution on 7H10 kanamycin plates containing 0 and 10ng/ml anhydrotetracycline (ATc). (**b**) *M. marinum* WT and *fecB* fs mutant containing pLJR965-sgRNA-*aftB* and pLJR965-sgRNA-*lpqW* were spotted in 10-fold serial dilutions on 7H10 kanamycin plates containing 0 and 1 ng/ml ATc.

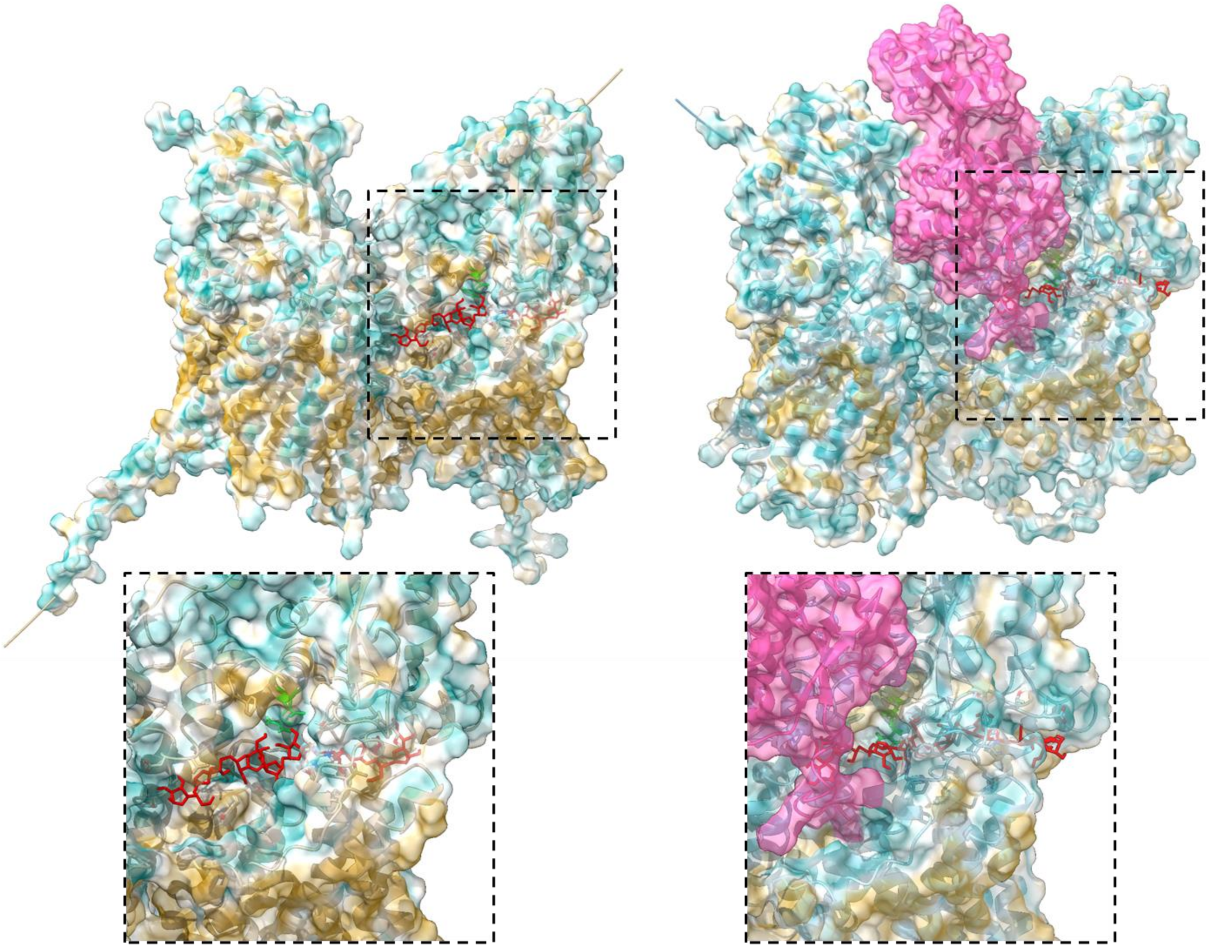

**Supplementary Figure 8: molecular docking of a synthetic galactan ligand to AftA**. A linear eight-residue galactofuranosyl chain (red) in the *M. marinum* AftA dimer (left) and AftA dimer with LpqZ (magenta, right) using HADDOCK version 2.4. The catalytic residues D108 and R112 are labeled in green.

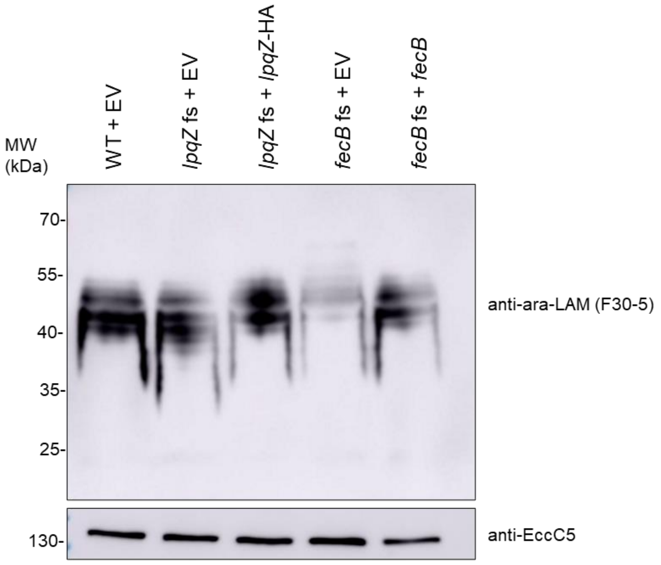


**Supplementary Figure 9: Lipoarabinomannan profiles of *lpqZ* and *fecB* complemented strains.** Whole cell lysates of *M. marinum* wild-type (WT), *lpqZ* and *fecB* fs mutants, containing the empty pSMT3 vector (EV) and genetically complemented strains were separated by SDS page and western blot was stained with Mab F30-5 antibody, recognizing terminal arabinan branches of LAM.

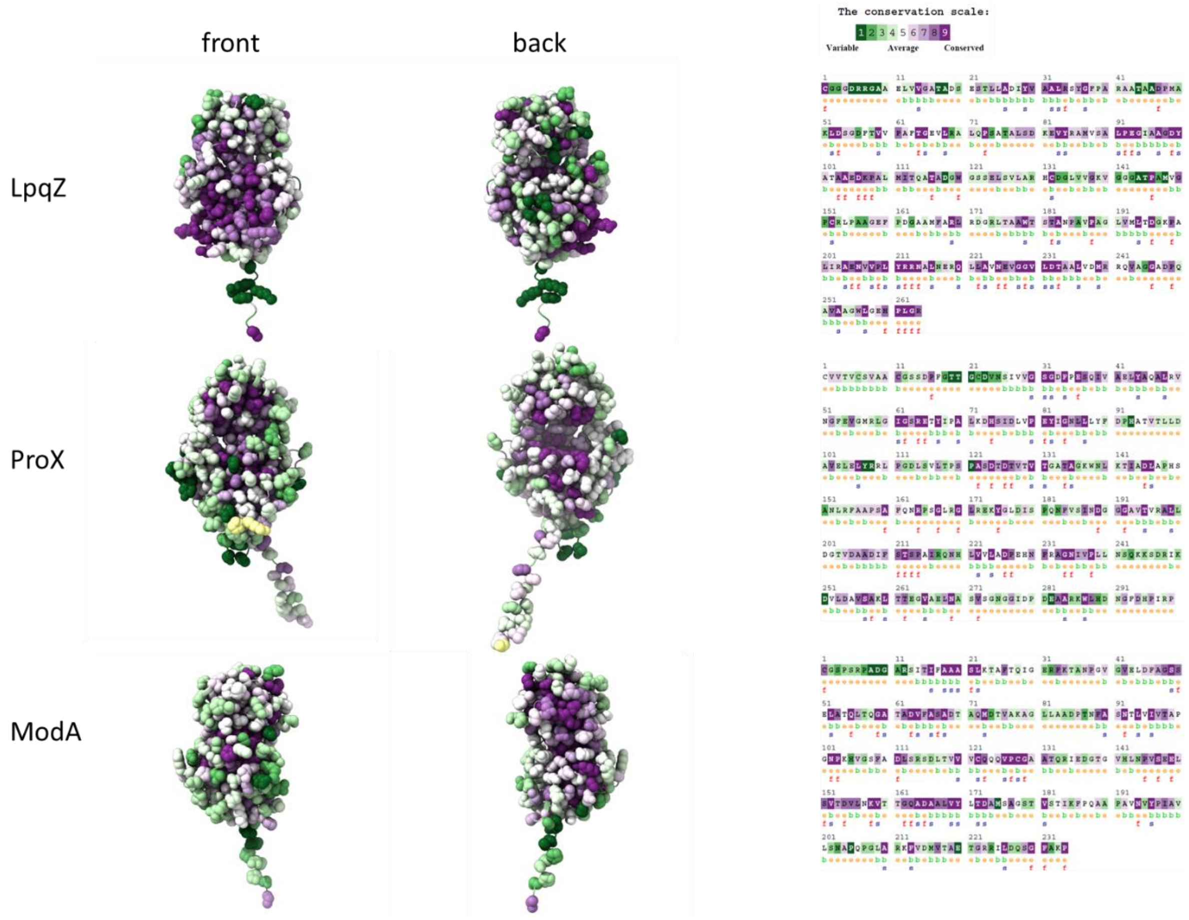


**Supplementary Figure 10**: **evolutionary conservation of LpqZ, ProX and ModA amino acids.** Conservation analysis of *M. marinum* LpqZ, ProX and ModA using ConSurf server. Proteins are depicted in atomic representation and amino acids are colored according to ConSurf color code (green (variable), white (average), purple (conserved), yellow (insufficient data).

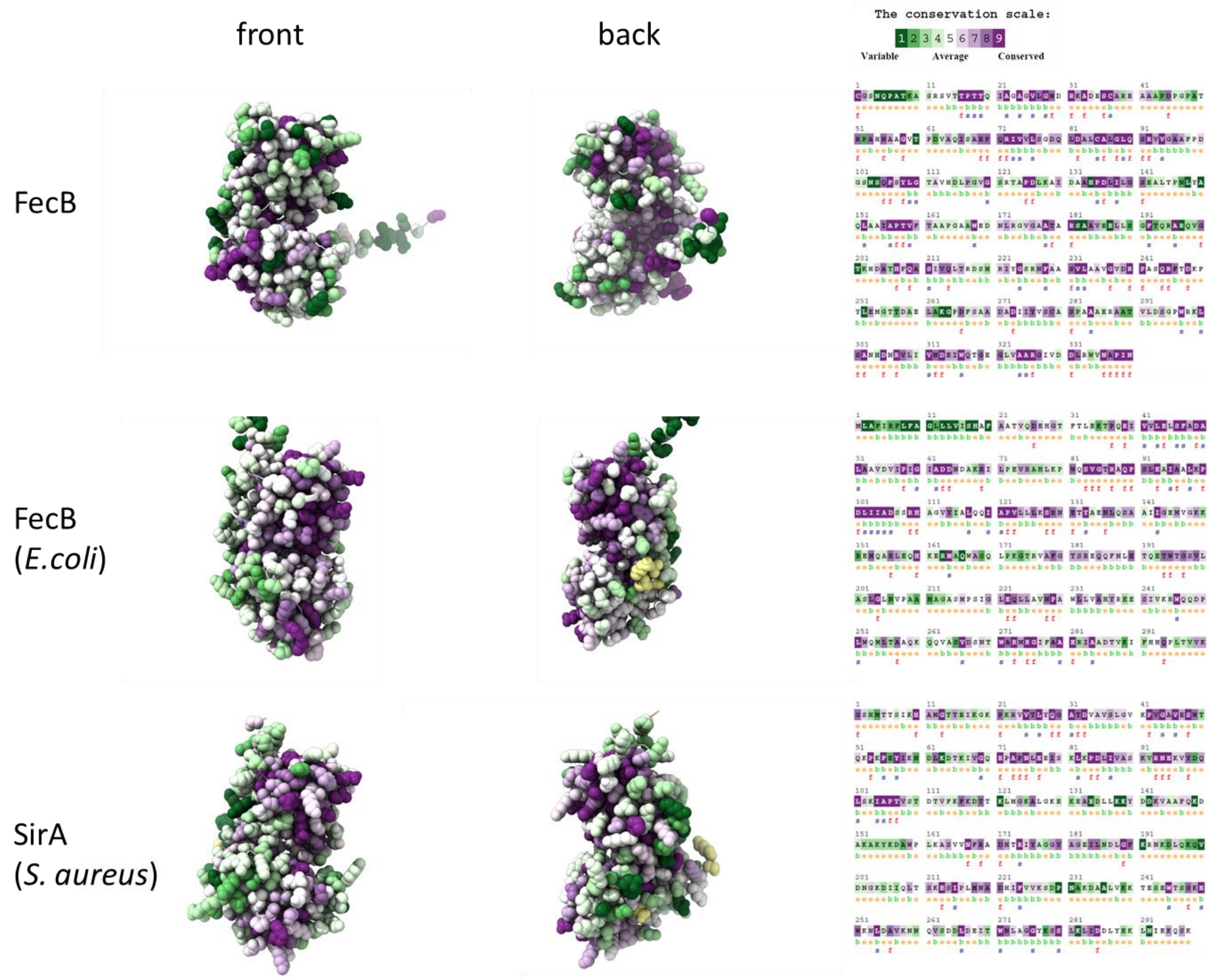


**Supplementary Figure 11: evolutionary conservation of FecB, FecB (*E.coli*) and SirA (*S. aureus*) amino acids.** Conservation analysis of *M. marinum* FecB, *E.coli* FecB and *S. aureus* SirA using ConSurf server. Proteins are depicted in atomic representation and amino acids are colored according to ConSurf color code (green (variable), white (average), purple (conserved), yellow (insufficient data).

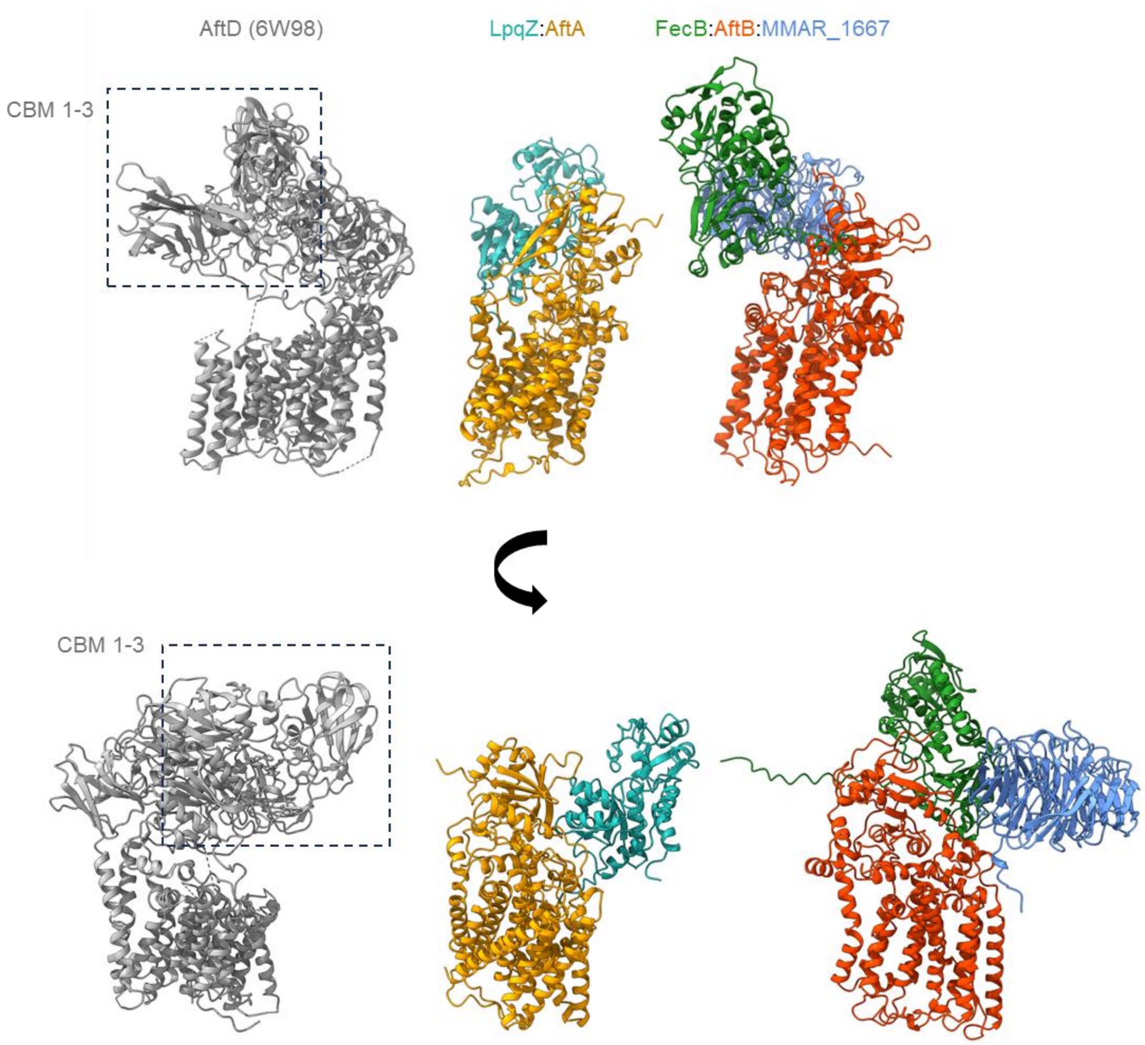


**Supplementary Figure 12: Comparison of AftD carbohydrate binding modules to LpqZ, FecB and MMAR_1667.** AftD (PDB:6W98) and its carbohydrate binding modules (CBMs) 1-3 were represented in grey by ChimeraX, as described by Tan *et al*.[1]. AlphaFold predictions of *M. marinum* LpqZ:AftA and FecB:AftB:MMAR_1667 from Figure 3c were compared in similar perspectives in ChimeraX.

**Supplementary Table 1: overview about putative lipoproteins.** Predicted lipoproteins according to Sutcliffe and Harrington[2] and annotated lipoproteins in Mycobrowser (https://mycobrowser.epfl.ch/).

| gene M. marinum | mmar M. marinum | gene Mtb | Rv Mtb | note |
|---|---|---|---|---|
| blaC | mmar_3050 | blaC | Rv2068c | in this study |
| dppA | mmar_5154 | dppA | Rv3666c | in this study |
| dsbF | mmar_5057 | dsbF | Rv1677 | in this study |
| fecB | mmar_1650 | fecB | Rv3044 | in this study |
| fgd2 | mmar_4200 | fgd2 | Rv0132c | in this study |
| ggtB | mmar_3713 | ggtB | Rv2394 | in this study |
| glnH | mmar_0714 | glnH | Rv0411c | in this study |
| lppD | mmar_2794 | lppD | Rv1899c | in this study |
| lppE | mmar_2769 | lppE | Rv1881c | in this study |
| lppF | mmar_0949 | lppF | Rv1921c | in this study |
| lppH | mmar_5074 | lppH | Rv3576 | in this study |
| lppI | mmar_3021 | lppI | Rv2046 | in this study |
| lppJ | mmar_5467 | lppJ | Rv2080 | in this study |
| lppK | mmar_3092 | lppK | Rv2116 | in this study |
| lppL | mmar_3118 | lppL | Rv2138 | in this study |
| lppM | mmar_3206 | lppM | Rv2171 | in this study |
| lppN | mmar_3366 | lppN | Rv2270 | in this study |
| lppP | mmar_3638 | lppP | Rv2330c | in this study |
| lppR | mmar_3722 | lppR | Rv2403c | in this study |
| lppS | mmar_3872 | ldtB | Rv2518c | in this study |
| lppU | mmar_1923 | lppU | Rv2784c | in this study |
| lppV_1 | mmar_1914 | lppV | Rv2796c | in this study |
| lppZ | mmar_1706 | lppZ | Rv3006 | in this study |
| lpqA | mmar_1691 | lpqA | Rv3016 | in this study |
| lpqC | mmar_1236 | lpqC | Rv3298c | in this study |
| lpqD | mmar_1154 | lpqD | Rv3390 | in this study |
| lpqE | mmar_5084 | lpqE | Rv3584 | in this study |
| lpqF | mmar_5092 | lpqF | Rv3593 | in this study |
| lpqG | mmar_5123 | lpqG | Rv3623 | in this study |
| lpqH | mmar_5315 | lpqH | Rv3763 | in this study |
| lpqI | mmar_0499 | lpqI | Rv0237 | in this study |
| lpqJ | mmar_0620 | lpqJ | Rv0344c | in this study |
| lpqK | mmar_0696 | lpqK | Rv0399c | in this study |
| lpqL | mmar_0726 | lpqL | Rv0418 | in this study |
| lpqM | mmar_0728 | lpqM | Rv0419 | in this study |

| | | | | |
|---|---|---|---|---|
| lpqN | mmar_0950 | lpqN | Rv0583c | in this study |
| lpqP | mmar_1000 | lpqP | Rv0671 | in this study |
| lpqQ | mmar_1362 | lpqQ | Rv0835 | in this study |
| lpqR | mmar_4800 | lpqR | Rv0838 | in this study |
| lpqS | mmar_4769 | lpqS | Rv0847 | in this study |
| lpqT | mmar_4470 | lpqT | Rv1016c | in this study |
| lpqU | mmar_4463 | lpqU | Rv1022 | in this study |
| lpqV | mmar_4402 | lpqV | Rv1064c | in this study |
| lpqY | mmar_4206 | lpqY | Rv1235 | in this study |
| lpqZ | mmar_4196 | lpqZ | Rv1244 | in this study |
| lprA | mmar_4152 | lprA | Rv1270c | in this study |
| lprB | mmar_4146 | lprB | Rv1274 | in this study |
| lprC | mmar_4145 | lprC | Rv1275 | in this study |
| lprD | mmar_4038 | lprD | Rv1343c | in this study |
| lprE | mmar_4190 | lprE | Rv1252c | in this study |
| lprF | mmar_2188 | lprF | Rv1368 | in this study |
| lprG | mmar_2220 | lprG | Rv1411c | in this study |
| lprH | mmar_2225 | lprH | Rv1418 | in this study |
| lprJ | mmar_2495 | lprJ | Rv1690 | in this study |
| lprK | mmar_0416 | lprK | Rv0173 | in this study |
| lprL | mmar_2364 | lprL | Rv0593 | in this study |
| lprM | mmar_2886 | lprM | Rv1970 | in this study |
| lprN | mmar_4983 | lprN | Rv3495c | in this study |
| lprO | mmar_0422 | lprO | Rv0179c | in this study |
| lprQ | mmar_0809 | lprQ | Rv0483 | in this study |
| mmar_0525 | mmar_0525 | Rv0265c | Rv0265c | in this study |
| mmar_1840 | mmar_1840 | Rv2864c | Rv2864c | in this study |
| mmar_2122 | mmar_2122 | Rv2585c | Rv2585c | in this study |
| mmar_2827 | mmar_2827 | Rv1922 | Rv1922 | in this study |
| mmar_3014 | mmar_3014 | Rv2041c | Rv2041c | in this study |
| modA | mmar_2731 | modA | Rv1857 | in this study |
| mpt70 | mmar_1834 | mpt83 | Rv2873 | in this study |
| oppA | mmar_4139 | oppA | Rv1280c | in this study |
| proX | mmar_5302 | proX | Rv3759c | in this study |
| pstS2 | mmar_4576 | pstS2 | Rv0932c | in this study |
| pstS3 | mmar_4580 | pstS3 | Rv0928 | in this study |
| rpfB | mmar_4479 | rpfB | Rv1009 | in this study |
| ugpB | mmar_1900 | ugpB | Rv2833c | in this study |
| uspC | mmar_3619 | uspC | Rv2318 | in this study |
| lppW | mmar_1803 | lppW | Rv2905 | no fs mutant could be made |
| lppX | mmar_1763 | lppX | Rv3244c | no fs mutant could be made |
| lpqB | mmar_1301 | lpqB | Rv2945c | no fs mutant could be made |
| lpqW | mmar_4288 | lpqW | Rv1166 | no fs mutant could be made |
| subI | mmar_3718 | subI | Rv2400c | no fs mutant could be made |
| - | - | lppA | Rv2543 | not in M. marinum |
| - | - | lppB | Rv2544 | not in M. marinum |
| - | - | lppC | Rv1911c | not in M. marinum |
| - | - | lppG | Rv1946c | not in M. marinum |
| - | - | lppO | Rv2290 | not in M. marinum |
| - | - | lppQ | Rv2341 | not in M. marinum |
| - | - | lppT | Rv1799 | not in M. marinum |
| - | - | lppY | Rv2999 | not in M. marinum |
| - | - | lpqO | Rv0604 | not in M. marinum |
| - | - | lpqX | Rv1228 | not in M. marinum |
| - | - | lprI | Rv1541c | not in M. marinum |
| - | - | lprP | Rv0962c | not in M. marinum |
| - | - | pstS1 | Rv0934 | not in M. marinum |
| - | - | Rv0381c | Rv0381c | not in M. marinum |
| - | - | caeA | Rv2224c | not in M. marinum |
| - | - | Rv2251 | Rv2251 | not in M. marinum |
| - | - | Rv2672 | Rv2672 | not in M. marinum |
| - | - | fecB2 | Rv0265c | not in M. marinum |
| - | - | Rv0460 | Rv0460 | not in M. marinum |
| - | - | Rv0526 | Rv0526 | not in M. marinum |
| - | - | Rv2293c | Rv2293c | not in M. marinum |
| - | - | Rv0846c | Rv0846c | not in M. marinum |
| - | - | Rv2843 | Rv2843 | not in M. marinum |
| - | - | Rv0679c | Rv0679c | not in M. marinum |
| mce6E | mmar_0181 | - | - | not in Mtb |
| mmar_4690 | mmar_4690 | - | - | not in Mtb |
| mmar_4706 | mmar_4706 | - | - | not in Mtb |
| mmar_5244 | mmar_5244 | - | - | not in Mtb |
| mmar_4884 | mmar_4884 | - | - | not in Mtb |
| mmar_5075 | mmar_5075 | - | - | not in Mtb |
| lppR_1 | mmar_3723 | - | - | not in Mtb |
| lpqL_1 | mmar_0727 | - | - | not in Mtb |
| LpqP like | mmar_0217 | - | - | not in Mtb |
| lpqP_1 | mmar_0227 | - | - | not in Mtb |
| lprM_1 | mmar_4055 | - | - | not in Mtb |
| phoS2_1 | mmar_1234 | - | - | not in Mtb |
| sodC | mmar_0747 | sodC | Rv0432 | omitted due to oversight |

**Supplementary Table 2: Foldseek searches of FecB and LpqZ.** Top 10 hits of structural similarity searches for FecB (7UQ0) and LpqZ (AF-O50459-F1) using Foldseek.

| input | target | description | organism | sequence identity | E-Value |
|---|---|---|---|---|---|
| **fecB (7UQ0)** | AF-O53291-F1-model_v4 | Probable FEIII-dicitrate-binding periplasmic lipoprotein FecB | Mycobacterium tuberculosis H37Rv | 96.5 | 1.11E-61 |
| | AF-X8FEP3-F1-model_v4 | Periplasmic binding family protein | Mycobacterium ulcerans str. Harvey | 77.8 | 7.96E-55 |
| | AF-Q9CBQ4-F1-model_v4 | Putative FeIII-dicitrate transporter lipoprotein | Mycobacterium leprae TN | 76.8 | 1.92E-53 |
| | AF-K0F3R8-F1-model_v4 | Transporter | Nocardia brasiliensis ATCC 700358 | 32.7 | 1.79E-28 |
| | AF-Q2G1N4-F1-model_v4 | **Periplasmic binding protein, putative (SirA)** | Staphylococcus aureus subsp. aureus NCTC 8325 | 20.8 | 5.96E-21 |
| | AF-P15028-F1-model_v4 | **Fe(3+) dicitrate-binding periplasmic protein (FecB)** | Escherichia coli K-12 | 20 | 1.13E-18 |
| | AF-Q2FZM4-F1-model_v4 | Fe/B12 periplasmic-binding domain-containing protein | Staphylococcus aureus subsp. aureus NCTC 8325 | 18.2 | 5.22E-18 |
| | AF-Q2FW75-F1-model_v4 | ABC transporter periplasmic binding protein, putative | Staphylococcus aureus subsp. aureus NCTC 8325 | 17.9 | 1.12E-17 |
| | AF-A0A133CZA2-F1-model_v4 | Iron ABC transporter substrate-binding protein | Enterococcus faecium | 19.8 | 5.54E-18 |
| | AF-K0F4D2-F1-model_v4 | Iron-siderophore binding protein | Nocardia brasiliensis ATCC 700358 | 23.2 | 6.61E-18 |
| **lpqZ (AF-O50459-F1)** | AF-O50459-F1-model_v4 | Probable lipoprotein LpqZ | Mycobacterium tuberculosis H37Rv | 100 | 5.45E-58 |
| | AF-Q9CC99-F1-model_v4 | Lipoprotein | Mycobacterium leprae TN | 73 | 4.75E-47 |
| | AF-K0F5R4-F1-model_v4 | Putative transporter substrate-binding protein | Nocardia brasiliensis ATCC 700358 | 29.1 | 5.37E-24 |
| | AF-X8F366-F1-model_v4 | Substrate binding domain of ABC-type glycine betaine transport system family protein | Mycobacterium ulcerans str. Harvey | 23 | 2.85E-20 |
| | AF-O69725-F1-model_v4 | **Possible osmoprotectant (Glycine betaine/carnitine/choline/L-proline) binding lipoprotein ProX** | Mycobacterium tuberculosis H37Rv | 23.7 | 5.72E-19 |
| | AF-K0FC11-F1-model_v4 | ABC amino acid transporter, substrate binding component | Nocardia brasiliensis ATCC 700358 | 29.1 | 1.28E-19 |
| | AF-Q8ZPK2-F1-model_v4 | Osmoprotectant-binding protein OsmX | Salmonella enterica subsp. enterica serovar Typhimurium str. LT2 | 17.4 | 1.06E-16 |
| | AF-A0A132P2M9-F1-model_v4 | ABC transporter permease subunit | Enterococcus faecium | 17.3 | 1.57E-15 |
| | AF-Q8DNJ9-F1-model_v4 | ABC transporter membrane-spanning permease-choline transporter | Streptococcus pneumoniae R6 | 19 | 6.46E-14 |
| | AF-Q9HXC4-F1-model_v4 | Probable binding protein component of ABC transporter | Pseudomonas aeruginosa PAO1 | 20.5 | 3.86E-15 |

**Supplementary Table 3: bacterial strains used in this study.** Used strains with their characteristics, containing plasmids and origin.

| strain | characteristics | plasmids | origin | reference |
| --- | --- | --- | --- | --- |
| Mycobacterium marinum M WT | WT strain | - | | [3] |
| C20: Mycobacterium marinum M WT pCRISPRx-Sth1 Cas9-L5 | Mycobacterium marinum M WT containing the empty pCRISPRx-Sth1 Cas9-L5 | pCRISPRx-Sth1 Cas9-L5 | Mycobacterium marinum M (USA) WT | This study |
| B2: Mycobacterium marinum M *dppA* | deletion in mmar_5154, CP000854.1:g.6238426_6238439del p.(T399_N402delS403Hfs*60) | pCRISPRx-Sth1 Cas9-L5-sgRNA-x | Mycobacterium marinum M (USA) WT | This study |
| B3: Mycobacterium marinum M *fecB* | deletion in mmar_1650, CP000854.1:g.1996165_1996180del p.(G125_Q129delS131Pfs*21) | pCRISPRx-Sth1 Cas9-L5-sgRNA-x | Mycobacterium marinum M (USA) WT | This study |
| B4: Mycobacterium marinum M *lppE* | deletion in mmar_2769, CP000854.1:g.3376280_3376343del p.(P29_C49delS50Vfs*23) | pCRISPRx-Sth1 Cas9-L5-sgRNA-x | Mycobacterium marinum M (USA) WT | This study |
| B5: Mycobacterium marinum M *lppF* | insertion in mmar_0949, CP000854.1:g.1158304_1158305insT p.(V89Rfs*87) | pCRISPRx-Sth1 Cas9-L5-sgRNA-x | Mycobacterium marinum M (USA) WT | This study |
| B6: Mycobacterium marinum M *dsbF* | deletion in mmar_5057, CP000854.1:g.6139071_6139159del p.(W81_A109delE111Gfs*182Mext*296) | pCRISPRx-Sth1 Cas9-L5-sgRNA-x | Mycobacterium marinum M (USA) WT | This study |
| B8: Mycobacterium marinum M *lppI* | deletion in mmar_3021, CP000854.1:g.3644530_3644540del p.(T43_K45delL46Sfs*27) | pCRISPRx-Sth1 Cas9-L5-sgRNA-x | Mycobacterium marinum M (USA) WT | This study |
| B9: Mycobacterium marinum M *lppJ* | deletion in mmar_5467, CP000854.1:g.6613219_6613237del p.(C49_D54delP55Rfs*9) | pCRISPRx-Sth1 Cas9-L5-sgRNA-x | Mycobacterium marinum M (USA) WT | This study |
| B10: Mycobacterium marinum M *lppM* | deletion in mmar_3206, CP000854.1:g.3910199_3910203del p.(A62delK63Tfs*243Lext*49) | pCRISPRx-Sth1 Cas9-L5-sgRNA-x | Mycobacterium marinum M (USA) WT | This study |
| B12: Mycobacterium marinum M *lppN* | deletion in mmar_3366, CP000854.1:g.4150222_4150223del p.(A28Gfs*36) | pCRISPRx-Sth1 Cas9-L5-sgRNA-x | Mycobacterium marinum M (USA) WT | This study |
| B14: Mycobacterium marinum M *lppU* | deletion in mmar_1923, CP000854.1:g.2338302_2338303del p.(V8Afs*28) | pCRISPRx-Sth1 Cas9-L5-sgRNA-x | Mycobacterium marinum M (USA) WT | This study |
| B16: Mycobacterium marinum M *lppV_1* | deletion in mmar_1914, CP000854.1:g.2330300_2330303del p.(L27delR29Vfs*9) | pCRISPRx-Sth1 Cas9-L5-sgRNA-x | Mycobacterium marinum M (USA) WT | This study |
| B17: Mycobacterium marinum M *lppK* | indel in mmar_3092, CP000854.1:g.3744552_3744553insC3744553_3744584del p.(L63_P72delG73Qfs*34) | pCRISPRx-Sth1 Cas9-L5-sgRNA-x | Mycobacterium marinum M (USA) WT | This study |
| B18: Mycobacterium marinum M *lppL* | deletion in mmar_3118, CP000854.1:g.3802106_3802125del p.(P56_R61delP62Afs*57) | pCRISPRx-Sth1 Cas9-L5-sgRNA-x | Mycobacterium marinum M (USA) WT | This study |
| B20: Mycobacterium marinum M *lppP* | deletion in mmar_3638, CP000854.1:g.4477931del p.(C38Afs*46) | pCRISPRx-Sth1 Cas9-L5-sgRNA-x | Mycobacterium marinum M (USA) WT | This study |
| B22: Mycobacterium marinum M *lpqC* | deletion in mmar_1236, CP000854.1:g.1506446_1506449del p.(A120Efs*96) | pCRISPRx-Sth1 Cas9-L5-sgRNA-x | Mycobacterium marinum M (USA) WT | This study |
| B24: Mycobacterium marinum M *lpqD* | deletion in mmar_1154, CP000854.1:g.1386441_1386447del p.(T51delE52Cfs*6) | pCRISPRx-Sth1 Cas9-L5-sgRNA-x | Mycobacterium marinum M (USA) WT | This study |
| B26: Mycobacterium marinum M *lppZ* | insertion in mmar_1706, CP000854.1:g.2056325_2056326insG p.(S105Qfs*17) | pCRISPRx-Sth1 Cas9-L5-sgRNA-x | Mycobacterium marinum M (USA) WT | This study |
| B27: Mycobacterium marinum M *lpqA* | insertion in mmar_1691, CP000854.1:g.2038389_2038390insC p.(P84Afs*32) | pCRISPRx-Sth1 Cas9-L5-sgRNA-x | Mycobacterium marinum M (USA) WT | This study |
| B29: Mycobacterium marinum M *lpqE* | deletion in mmar_5084, CP000854.1:g.6163669_6163717del p.(P68_P83delD84Ifs*18) | pCRISPRx-Sth1 Cas9-L5-sgRNA-x | Mycobacterium marinum M (USA) WT | This study |
| B31: Mycobacterium marinum M *lpqG* | deletion in mmar_5123, CP000854.1:g.6202391_6202407del p.(V73_M77delD72Efs*22) | pCRISPRx-Sth1 Cas9-L5-sgRNA-x | Mycobacterium marinum M (USA) WT | This study |
| B33: Mycobacterium marinum M *lpqH* | deletion in mmar_5315, CP000854.1:g.6422316_6422329del p.(N15_I18delV14Gfs*100Lext*5) | pCRISPRx-Sth1 Cas9-L5-sgRNA-x | Mycobacterium marinum M (USA) WT | This study |
| B35: Mycobacterium marinum M *lpqI* | insertion in mmar_0499, CP000854.1:g.590503_590504insT p.(T6Ifs*34) | pCRISPRx-Sth1 Cas9-L5-sgRNA-x | Mycobacterium marinum M (USA) WT | This study |
| B36: Mycobacterium marinum M *lpqJ* | insertion in mmar_0620, CP000854.1:g.732794_732795insT p.(R48Qfs*108) | pCRISPRx-Sth1 Cas9-L5-sgRNA-x | Mycobacterium marinum M (USA) WT | This study |
| B38: Mycobacterium marinum M *lpqY* | insertion in mmar_4206, CP000854.1:g.5183644_5183645insCTGGACATCGGCATGC p.(T148Cfs*26) | pCRISPRx-Sth1 Cas9-L5-sgRNA-x | Mycobacterium marinum M (USA) WT | This study |
| B39: Mycobacterium marinum M *lprA* | deletion in mmar_4152, CP000854.1:g.5113541_5113581del p.(A58_T70delE57Gfs*64) | pCRISPRx-Sth1 Cas9-L5-sgRNA-x | Mycobacterium marinum M (USA) WT | This study |
| B41: Mycobacterium marinum M *glnH* | deletion in mmar_0714, CP000854.1:g.861494_861503del p.(D60_S62delS63Afs*32) | pCRISPRx-Sth1 Cas9-L5-sgRNA-x | Mycobacterium marinum M (USA) WT | This study |
| B42: Mycobacterium marinum M *lppD* | indel in mmar_2794, CP000854.1:g.3396955_3396956insGCCGCTCGCTGGCCTTGGTGGCATTTGGCACCGGCGTCGGCGGCTTCCCGCT3396956_3397009del p.(C106*) | pCRISPRx-Sth1 Cas9-L5-sgRNA-x | Mycobacterium marinum M (USA) WT | This study |
| B44: Mycobacterium marinum M *lppH* | insertion in mmar_5074, CP000854.1:g.6154735_6154736insT p.(D86Gfs*35) | pCRISPRx-Sth1 Cas9-L5-sgRNA-x | Mycobacterium marinum M (USA) WT | This study |
| B46: Mycobacterium marinum M *lppR* | deletion in mmar_3722, CP000854.1:g.4585310_4585319del p.(Q59_F61delT62Rfs*66) | pCRISPRx-Sth1 Cas9-L5-sgRNA-x | Mycobacterium marinum M (USA) WT | This study |
| B47: Mycobacterium marinum M *lpqK* | deletion in mmar_0696, CP000854.1:g.832138_832142del p.(V7delA8Wfs*145) | pCRISPRx-Sth1 Cas9-L5-sgRNA-x | Mycobacterium marinum M (USA) WT | This study |
| B49: Mycobacterium marinum M *lpqM* | deletion in mmar_0728, CP000854.1:g.872835_872845del p.(S148_C150delN151Hfs*19) | pCRISPRx-Sth1 Cas9-L5-sgRNA-x | Mycobacterium marinum M (USA) WT | This study |
| B51: Mycobacterium marinum M *lpqP* | deletion in mmar_1000, CP000854.1:g.1225418_1225428del p.(I38_V40delG41Rfs*42) | pCRISPRx-Sth1 Cas9-L5-sgRNA-x | Mycobacterium marinum M (USA) WT | This study |
| B52: Mycobacterium marinum M *lpqQ* | deletion in mmar_1362, CP000854.1:g.1656737_1656738del p.(V75Rfs*8) | pCRISPRx-Sth1 Cas9-L5-sgRNA-x | Mycobacterium marinum M (USA) WT | This study |

| | | | | |
|---|---|---|---|---|
| B53: Mycobacterium marinum M *lpqU* | deletion in mmar_4463, CP000854.1:g.5481908_5481909del p.(E88Gfs*58) | pCRISPRx-Sth1 Cas9-L5-sgRNA-x | Mycobacterium marinum M (USA) WT | This study |
| B54: Mycobacterium marinum M *lpqV* | deletion in mmar_4402, CP000854.1:g.5415081_5415091del p.(E81_T83delA80Gfs*156Iext*34) | pCRISPRx-Sth1 Cas9-L5-sgRNA-x | Mycobacterium marinum M (USA) WT | This study |
| B55: Mycobacterium marinum M *lprC* | deletion in mmar_4145, CP000854.1:g.5105102_5105115del p.(Y59_L62delQ58Pfs*5) | pCRISPRx-Sth1 Cas9-L5-sgRNA-x | Mycobacterium marinum M (USA) WT | This study |
| B57: Mycobacterium marinum M *lpqN* | deletion in mmar_0950, CP000854.1:g.1159251_1159255del p.(D69delI70Pfs*44) | pCRISPRx-Sth1 Cas9-L5-sgRNA-x | Mycobacterium marinum M (USA) WT | This study |
| B58: Mycobacterium marinum M *lpqT* | deletion in mmar_4470, CP000854.1:g.5491540_5491544del p.(G60delV61Qfs*219Vext*263) | pCRISPRx-Sth1 Cas9-L5-sgRNA-x | Mycobacterium marinum M (USA) WT | This study |
| B59: Mycobacterium marinum M *lprD* | deletion in mmar_4038, CP000854.1:g.4980853_4980916del p.(L47_F67delV68Cfs*16) | pCRISPRx-Sth1 Cas9-L5-sgRNA-x | Mycobacterium marinum M (USA) WT | This study |
| B60: Mycobacterium marinum M *lprE* | deletion in mmar_4190, CP000854.1:g.5159695_5159711del p.(S33_K37delT38Pfs*60) | pCRISPRx-Sth1 Cas9-L5-sgRNA-x | Mycobacterium marinum M (USA) WT | This study |
| B62: Mycobacterium marinum M *uspC* | insertion in mmar_3619, CP000854.1:g.4444537_4444538insT p.(S100Ffs*11) | pCRISPRx-Sth1 Cas9-L5-sgRNA-x | Mycobacterium marinum M (USA) WT | This study |
| B63: Mycobacterium marinum M *lppS* | deletion in mmar_3872, CP000854.1:g.4811945_4811955del p.(A34_K36delV37Dfs*69) | pCRISPRx-Sth1 Cas9-L5-sgRNA-x | Mycobacterium marinum M (USA) WT | This study |
| B65: Mycobacterium marinum M *mmar_1900* | deletion in mmar_1900, CP000854.1:g.2314663_2314675del p.(Q49_S51delG48Afs*6) | pCRISPRx-Sth1 Cas9-L5-sgRNA-x | Mycobacterium marinum M (USA) WT | This study |
| B66: Mycobacterium marinum M *lprF* | insertion in mmar_2188, CP000854.1:g.2633975_2633976insA p.(P25Tfs*202) | pCRISPRx-Sth1 Cas9-L5-sgRNA-x | Mycobacterium marinum M (USA) WT | This study |
| B67: Mycobacterium marinum M *lprG* | insertion in mmar_2220, CP000854.1:g.2670518_2670519insA p.(T49Hfs*71) | pCRISPRx-Sth1 Cas9-L5-sgRNA-x | Mycobacterium marinum M (USA) WT | This study |
| B68: Mycobacterium marinum M *lprJ* | deletion in mmar_2495, CP000854.1:g.3039763_3039764del p.(L26Dfs*130Text*8) | pCRISPRx-Sth1 Cas9-L5-sgRNA-x | Mycobacterium marinum M (USA) WT | This study |
| B69: Mycobacterium marinum M *lprK* | insertion in mmar_0416, CP000854.1:g.482779_482780insA p.(L103Afs*38) | pCRISPRx-Sth1 Cas9-L5-sgRNA-x | Mycobacterium marinum M (USA) WT | This study |
| B71: Mycobacterium marinum M *lprL* | deletion in mmar_2364, CP000854.1:g.2853425_2853426del p.(Q105Efs*3) | pCRISPRx-Sth1 Cas9-L5-sgRNA-x | Mycobacterium marinum M (USA) WT | This study |
| B73: Mycobacterium marinum M lprM | deletion in mmar_2886, CP000854.1:g.3492920_3492921del p.(V78Gfs*49) | pCRISPRx-Sth1 Cas9-L5-sgRNA-x | Mycobacterium marinum M (USA) WT | This study |
| B74: Mycobacterium marinum M *lprO* | insertion in mmar_0422, CP000854.1:g.489249_489250insA p.(V106Rfs*15) | pCRISPRx-Sth1 Cas9-L5-sgRNA-x | Mycobacterium marinum M (USA) WT | This study |
| B75: Mycobacterium marinum M *proX* | deletion in mmar_5302, CP000854.1:g.6402158_6402174del p.(F69_M73delR74Vfs*183) | pCRISPRx-Sth1 Cas9-L5-sgRNA-x | Mycobacterium marinum M (USA) WT | This study |
| B76: Mycobacterium marinum M *pstS2* | deletion in mmar_4576, CP000854.1:g.5615285_5615300del p.(V43_G46delG47Lfs*44) | pCRISPRx-Sth1 Cas9-L5-sgRNA-x | Mycobacterium marinum M (USA) WT | This study |
| B77: Mycobacterium marinum M *psts3* | insertion in mmar_4580, CP000854.1:g.5621568_5621569insA p.(K47Efs*44) | pCRISPRx-Sth1 Cas9-L5-sgRNA-x | Mycobacterium marinum M (USA) WT | This study |
| B78: Mycobacterium marinum M *mmar_2827* | deletion in mmar_2827, CP000854.1:g.3428172_3428227del p.(A79_G96delI78Nfs*23) | pCRISPRx-Sth1 Cas9-L5-sgRNA-x | Mycobacterium marinum M (USA) WT | This study |
| B79: Mycobacterium marinum M *mmar_3014* | insertion in mmar_3014, CP000854.1:g.3637855_3637856insA p.(D35*) | pCRISPRx-Sth1 Cas9-L5-sgRNA-x | Mycobacterium marinum M (USA) WT | This study |
| B80: Mycobacterium marinum M *mmar_2122* | deletion in mmar_2122, CP000854.1:g.2559148_2559151del p.(P171delQ173Rfs*26) | pCRISPRx-Sth1 Cas9-L5-sgRNA-x | Mycobacterium marinum M (USA) WT | This study |
| B81: Mycobacterium marinum M *mmar_1840* | deletion in mmar_1840, CP000854.1:g.2240322_2240338del p.(R96_H100delL101Sfs*9) | pCRISPRx-Sth1 Cas9-L5-sgRNA-x | Mycobacterium marinum M (USA) WT | This study |
| C2: Mycobacterium marinum M lprH | deletion in mmar_2225, CP000854.1:g.2673858del p.(G29Vfs*93) | pCRISPRx-Sth1 Cas9-L5-sgRNA-x | Mycobacterium marinum M (USA) WT | This study |
| C3: Mycobacterium marinum M MPT70 | deletion in mmar_1834, CP000854.1:g.2233450_2233451del p.(P6Sfs*160) | pCRISPRx-Sth1 Cas9-L5-sgRNA-x | Mycobacterium marinum M (USA) WT | This study |
| C4: Mycobacterium marinum M oppA | insertion in mmar_4139, CP000854.1:g.5095841_5095842insC p.(D159Rfs*9) | pCRISPRx-Sth1 Cas9-L5-sgRNA-x | Mycobacterium marinum M (USA) WT | This study |
| C5: Mycobacterium marinum M lpqL | deletion in mmar_0726, CP000854.1:g.869646_869688del p.(E116_A129delH130Tfs*14) | pCRISPRx-Sth1 Cas9-L5-sgRNA-x | Mycobacterium marinum M (USA) WT | This study |
| C7: Mycobacterium marinum M lpqS | insertion in mmar_4769, CP000854.1:g.5821295_5821296insC p.(H46Qfs*9) | pCRISPRx-Sth1 Cas9-L5-sgRNA-x | Mycobacterium marinum M (USA) WT | This study |
| C9: Mycobacterium marinum M lpqZ | deletion in mmar_4196, CP000854.1:g.5172033_5172042del p.(E101_Y103delK100Nfs*27) | pCRISPRx-Sth1 Cas9-L5-sgRNA-x | Mycobacterium marinum M (USA) WT | This study |
| C10: Mycobacterium marinum M lprN | insertion in mmar_4983, CP000854.1:g.6048309_6048310insT p.(A86Sfs*87) | pCRISPRx-Sth1 Cas9-L5-sgRNA-x | Mycobacterium marinum M (USA) WT | This study |
| C11: Mycobacterium marinum M lprQ | insertion in mmar_0809, CP000854.1:g.971312_971313insT p.(E67Rfs*20) | pCRISPRx-Sth1 Cas9-L5-sgRNA-x | Mycobacterium marinum M (USA) WT | This study |
| C13: Mycobacterium marinum M modA | deletion in mmar_2731, CP000854.1:g.3331862_3331868del p.(L72_A73delT74Pfs*64) | pCRISPRx-Sth1 Cas9-L5-sgRNA-x | Mycobacterium marinum M (USA) WT | This study |
| C15: Mycobacterium marinum M *mmar_0525* | deletion in mmar_0525, CP000854.1:g.618262del p.(Y139Ifs*40) | pCRISPRx-Sth1 Cas9-L5-sgRNA-x | Mycobacterium marinum M (USA) WT | This study |
| C17: Mycobacterium marinum M lpqF | deletion in mmar_5092, CP000854.1:g.6170210_6170211insA p.0? | pCRISPRx-Sth1 Cas9-L5-sgRNA-x | Mycobacterium marinum M (USA) WT | This study |
| C19: Mycobacterium marinum M lpqR | deletion in mmar_4800, CP000854.1:g.5854542_5854761del p.(I82_W153delA81Gfs*19) | pCRISPRx-Sth1 Cas9-L5-sgRNA-x | Mycobacterium marinum M (USA) WT | This study |

| | | | | |
|---|---|---|---|---|
| C22: Mycobacterium marinum M rpfB | deletion in mmar_4479, CP000854.1:g.5502159_5502165del p.(L29_T30delV31Sfs*8) | pCRISPRx-Sth1 Cas9-L5-sgRNA-x | Mycobacterium marinum M (USA) WT | This study |
| C24: Mycobacterium marinum M ggtB | deletion in mmar_3713, CP000854.1:g.4573170_4573176del p.(Y133_Y134delD135Tfs*124) | pCRISPRx-Sth1 Cas9-L5-sgRNA-x | Mycobacterium marinum M (USA) WT | This study |
| C26: Mycobacterium marinum M fgd2 | deletion in mmar_4200, CP000854.1:g.5176887_5176903del p.(A83_G87delL88Gfs*116) | pCRISPRx-Sth1 Cas9-L5-sgRNA-x | Mycobacterium marinum M (USA) WT | This study |
| C27: Mycobacterium marinum M blaC | deletion in mmar_3050, CP000854.1:g.3694724_3694850del p.(L25_V65delY66Cfs*44) | pCRISPRx-Sth1 Cas9-L5-sgRNA-x | Mycobacterium marinum M (USA) WT | This study |
| C29: Mycobacterium marinum M lprB | insertion in mmar_4146, CP00854.1:g.5105682_5105683insT p.(Q54Tfs*55) | pCRISPRx-Sth1 Cas9-L5-sgRNA-x | Mycobacterium marinum M (USA) WT | This study |
| G64: Mycobacteriummarinum M mmar_1667 | deletion in mmar_1667, CP000854.1:g.2014237_2014250del p.(C100_M103delE104Vfs*103) | pCRISPRx-Sth1 Cas9-L5-sgRNA-x | Mycobacterium marinum M (USA) WT | This study |
| D24: Mycobacterium marinum M WT pTdTomato-L5 | contains pTdTomato-L5 (Addgene 140994) to exchange pCRISPRx-Sth1 Cas9-L5 | pTdTomato-L5 | C20: Mycobacterium marinum M WT pCRISPRx-Sth1 Cas9-L5 | This study |
| D11: Mycobacterium marinum M lpqZ pTdTomato-L5 | contains pTdTomato-L5 (Addgene 140994) to exchange pCRISPRx-Sth1 Cas9-L5 | pTdTomato-L5 | C9: Mycobacterium marinum M lpqZ | This study |
| C44: Mycobacterium marinum M fecB pTdTomato-L5 | contains pTdTomato-L5 (Addgene 140994) to exchange pCRISPRx-Sth1 Cas9-L5 | pTdTomato-L5 | B3: Mycobacterium marinum M *fecB* | This study |
| E12: Mycobacterium marinum M WT pSMT3-empty | strain D24 containing empty pSMT3 | pTdTomato-L5, pSMT3-empty | D24: Mycobacterium marinum M WT pTdTomato-L5 | This study |
| E8: Mycobacterium marinum M lpqZ pSMT3-empty | strain D11 containing empty pSMT3 | pTdTomato-L5, pSMT3-empty | D11: Mycobacterium marinum M lpqZ pTdTomato-L5 | This study |
| D78: Mycobacterium marinum M lpqZ pSMT3-lpqZ-HA | strain D11 containing pSMT3-lpqZ-HA | pTdTomato-L5, pSMT3-lpqZ-HA | D11: Mycobacterium marinum M lpqZ pTdTomato-L5 | This study |
| F50: Mycobacterium marinum M fecB pSMT3-empty | strain C44 containing empty pSMT3 | pTdTomato-L5, pSMT3-empty | C44: Mycobacterium marinum M fecB pTdTomato-L5 | This study |
| F24: Mycobacterium marinum M fecB pSMT3-fecB-HA | strain C44 containing pSMT3-fecB-HA | pTdTomato-L5, pSMT3-fecB-HA | C44: Mycobacterium marinum M fecB pTdTomato-L5 | This study |
| G37: Mycobacterium marinum M fecB pSMT3-fecB | strain C44 containing pSMT3-fecB | pTdTomato-L5, pSMT3-fecB | C44: Mycobacterium marinum M fecB pTdTomato-L5 | This study |
| F56: Mycobacterium marinum M WT pLJR965-sgRNA-AftA | strain D24 containing pLJR965-AftA | pTdTomato-L5, pLJR965-sgRNA-x | D24: Mycobacterium marinum M WT pTdTomato-L5 | This study |
| F58: Mycobacterium marinum M lpqZ pLJR965-sgRNA-AftA | strain D11 containing pLJR965-AftA | pTdTomato-L5, pLJR965-sgRNA-x | D11: Mycobacterium marinum M lpqZ pTdTomato-L5 | This study |
| F63: Mycobacterium marinum WT pLJR965-sgRNA-lpqW | strain D24 containing pLJR965-lpqW | pTdTomato-L5, pLJR965-sgRNA-x | D24: Mycobacterium marinum M WT pTdTomato-L5 | This study |
| F66: Mycobacterium marinum lpqZ pLJR965-sgRNA-lpqW | strain D11 containing pLJR965-lpqW | pTdTomato-L5, pLJR965-sgRNA-x | D11: Mycobacterium marinum M lpqZ pTdTomato-L5 | This study |
| G1: Mycobacterium marinum M WT pLJR965-sgRNA-aftB | strain B1 containing pLJR965-sgRNA-aftB | pLJR965-sgRNA-x | Mycobacterium marinum M (USA) WT | This study |
| G5: Mycobacterium marinum M fecB pLJR965-sgRNA-aftB | strain C44 containing pLJR965-sgRNA-aftB | pTdTomato-L5, pLJR965-sgRNA-x | C44: Mycobacterium marinum M fecB pTdTomato-L5 | This study |
| G7: Mycobacterium marinum M fecB pLJR965-sgRNA-lpqW | strain C44 containing pLJR965-sgRNA-lpqW | pTdTomato-L5, pLJR965-sgRNA-x | C44: Mycobacterium marinum M fecB pTdTomato-L5 | This study |
| G17: Mycobacterium marinum M WT pSMT3-empty pLJR962-AftA-FLAG | strain E12 containing pLJR962-AftA-FLAG | pTdTomato-L5, pSMT3-empty, pLJR962-AftA-FLAG | E12: Mycobacterium marinum M WT pSMT3-empty | This study |
| G20: Mycobacterium marinum M lpqZ pSMT3-lpqZ-HA pLJR962-AftA-FLAG | strain D78 containing pLJR962-AftA-FLAG | pTdTomato-L5, pSMT3-LpqZ-HA, pLJR962-AftA-FLAG | D78: Mycobacterium marinum M lpqZ pSMT3-lpqZ-HA | This study |
| D53: Mycobacterium marinum M WT pSMT3-MspA | WT strain containing pSMT3-MspA | pSMT3-MspA | Mycobacterium marinum M (USA) WT | [4] |
| G72: Mycobacterium marinum M mmar_1667 pTdTomato-L5.2 | strain G64 containing pTdTomato-L5 (Addgene 140994) to exchange pCRISPRx-Sth1 Cas9-L5 | pTdTomato-L5 | G64: Mycobacteriummarinum M mmar_1667 | This study |
| H15: Mycobacterium marinum M mmar_1667 pML1357kana-mmar1667-Strep | strain G72 containing pML1357kana-mmar1667-Strep | pTdTomato-L5, pML1357kana-mmar_1667-Strep | G72: Mycobacterium marinum M mmar_1667 pTdTomato-L5.2 | This study |
| H52: M. marinum lpqZ pSMT3-lpqZ(Mtb)-HA | strain D11 containing pSMT3-lpqZ(Mtb)-HA | pTdTomato-L5, pSMT3-lpqZ(Mtb)-HA | D11: Mycobacterium marinum M lpqZ pTdTomato-L5 | This study |

| | | | | |
|---|---|---|---|---|
| H54: M. marinum fecB pSMT3-fecB(Mtb) | strain C44 containing pSMT3-fecB(Mtb) | pTdTomato-L5, pSMT3-fecB(Mtb) | C44: Mycobacterium marinum M fecB pTdTomato-L5 | This study |

**Supplementary Table 4: plasmids used in this study.** Used plasmids, their characteristics and origin.

| plasmid | characteristics | origin | reference |
|---|---|---|---|
| pTdTomato-L5 | for replacement of pCRISPRx-Sth1-Cas9-L5 | Addgene 140994 | [5] |
| pCRISPRx-Sth1-Cas9-L5 | no gRNA | Addgene 140993 | [5] |
| pCRISPRx-Sth1-Cas9-L5-sgRNA-mmar_5154 | target gRNA: GACACGGTGGCCAACAGCATCAAGAA | pCRISPRx-Sth1 Cas9-L5 (Addgene 140993) | This study |
| pCRISPRx-Sth1-Cas9-L5-sgRNA-mmar_5057 | target gRNA: GCACTTGGGGCACCACGGTGTCCAGAA | pCRISPRx-Sth1 Cas9-L5 (Addgene 140993) | This study |
| pCRISPRx-Sth1-Cas9-L5-sgRNA-mmar_1650 | target gRNA: GGCTGGCTATTGGAACCGTCCGGGAA | pCRISPRx-Sth1 Cas9-L5 (Addgene 140993) | This study |
| pCRISPRx-Sth1-Cas9-L5-sgRNA-mmar_0714 | target gRNA: GTCCTGGCTGGAGCTGTCCGCTGGAA | pCRISPRx-Sth1 Cas9-L5 (Addgene 140993) | This study |
| pCRISPRx-Sth1-Cas9-L5-sgRNA-mmar_2794 | target gRNA: AGCCGTGCCGCCCGTTCGAGCGGGAA | pCRISPRx-Sth1 Cas9-L5 (Addgene 140993) | This study |
| pCRISPRx-Sth1-Cas9-L5-sgRNA-mmar_2769 | target gRNA: GCACGCCCCGCCGCATGCAGTCAGGAG | pCRISPRx-Sth1 Cas9-L5 (Addgene 140993) | This study |
| pCRISPRx-Sth1-Cas9-L5-sgRNA-mmar_0949 | target gRNA: GACTGGGACAACACCGTCGTCAAGAA | pCRISPRx-Sth1 Cas9-L5 (Addgene 140993) | This study |
| pCRISPRx-Sth1-Cas9-L5-sgRNA-mmar_5074 | target gRNA: ACTGGAGCAAGAACATCTCGGACAAGAA | pCRISPRx-Sth1 Cas9-L5 (Addgene 140993) | This study |
| pCRISPRx-Sth1-Cas9-L5-sgRNA-mmar_3021 | target gRNA: GTGGGAAGTTTGCTAGTCGAGGAAGAA | pCRISPRx-Sth1 Cas9-L5 (Addgene 140993) | This study |
| pCRISPRx-Sth1-Cas9-L5-sgRNA-mmar_5467 | target gRNA: ATCGCCTTGGTCATTGCATGACGAGAA | pCRISPRx-Sth1 Cas9-L5 (Addgene 140993) | This study |
| pCRISPRx-Sth1-Cas9-L5-sgRNA-mmar_3092 | target gRNA: GATCCGAACATTCCGGGAGACAAGAA | pCRISPRx-Sth1 Cas9-L5 (Addgene 140993) | This study |
| pCRISPRx-Sth1-Cas9-L5-sgRNA-mmar_3118 | target gRNA: GAACCACACCTGCCGGAGGCTGAGAA | pCRISPRx-Sth1 Cas9-L5 (Addgene 140993) | This study |
| pCRISPRx-Sth1-Cas9-L5-sgRNA-mmar_3206 | target gRNA: GATCATCGCGGCGGCAAAACCGAAGAA | pCRISPRx-Sth1 Cas9-L5 (Addgene 140993) | This study |
| pCRISPRx-Sth1-Cas9-L5-sgRNA-mmar_3366 | target gRNA: ACCACCACTCATCAAGCGGGGCAGGAA | pCRISPRx-Sth1 Cas9-L5 (Addgene 140993) | This study |
| pCRISPRx-Sth1-Cas9-L5-sgRNA-mmar_3638 | target gRNA: ACTACCGCGCCGGATACCTGCAAGAA | pCRISPRx-Sth1 Cas9-L5 (Addgene 140993) | This study |
| pCRISPRx-Sth1-Cas9-L5-sgRNA-mmar_3722 | target gRNA: ACTTGGCCGTGAAGTCCTGGCCGAGAA | pCRISPRx-Sth1 Cas9-L5 (Addgene 140993) | This study |
| pCRISPRx-Sth1-Cas9-L5-sgRNA-mmar_1923 | target gRNA: GCCAGAGCCATCAGCACGGCCAAGAA | pCRISPRx-Sth1 Cas9-L5 (Addgene 140993) | This study |
| pCRISPRx-Sth1-Cas9-L5-sgRNA-mmar_1914 | target gRNA: GCTCACCACCACGCCCAAGACCACCAGAA | pCRISPRx-Sth1 Cas9-L5 (Addgene 140993) | This study |
| pCRISPRx-Sth1-Cas9-L5-sgRNA-mmar_1803 | target gRNA: AACGCATCCAGCGGGCAACCGAGGAA | pCRISPRx-Sth1 Cas9-L5 (Addgene 140993) | This study |
| pCRISPRx-Sth1-Cas9-L5-sgRNA-mmar_1763 | target gRNA: GAGCCTCGGGTTGCGTTTTGTCAGAA | pCRISPRx-Sth1 Cas9-L5 (Addgene 140993) | This study |
| pCRISPRx-Sth1-Cas9-L5-sgRNA-mmar_1706 | target gRNA: GTCAAGGAGGTGTCGGTCAGCGCAGAA | pCRISPRx-Sth1 Cas9-L5 (Addgene 140993) | This study |
| pCRISPRx-Sth1-Cas9-L5-sgRNA-mmar_1691 | target gRNA: GCCCTGGCAAAGACCGCGCCGGTGGAA | pCRISPRx-Sth1 Cas9-L5 (Addgene 140993) | This study |
| pCRISPRx-Sth1-Cas9-L5-sgRNA-mmar_1301 | target gRNA: ATGGATCCCGACGTGCTGCTGCGGGAA | pCRISPRx-Sth1 Cas9-L5 (Addgene 140993) | This study |
| pCRISPRx-Sth1-Cas9-L5-sgRNA-mmar_1236 | target gRNA: GTACCTTGCCCACTAGGGCCACCAGAA | pCRISPRx-Sth1 Cas9-L5 (Addgene 140993) | This study |
| pCRISPRx-Sth1-Cas9-L5-sgRNA-mmar_1154 | target gRNA: GATCATCGCTACCGAGTTGCCTGGAA | pCRISPRx-Sth1 Cas9-L5 (Addgene 140993) | This study |
| pCRISPRx-Sth1-Cas9-L5-sgRNA-mmar_5084 | target gRNA: GTCCACATTGTGGCCGGGACGCAGGAA | pCRISPRx-Sth1 Cas9-L5 (Addgene 140993) | This study |
| pCRISPRx-Sth1-Cas9-L5-sgRNA-mmar_5123 | target gRNA: GCTCATCGCGCCGGTGACGTCCTGAGAA | pCRISPRx-Sth1 Cas9-L5 (Addgene 140993) | This study |
| pCRISPRx-Sth1-Cas9-L5-sgRNA-mmar_5315 | target gRNA: ACGTCAACATCGCGATCGGTGGAGCA | pCRISPRx-Sth1 Cas9-L5 (Addgene 140993) | This study |
| pCRISPRx-Sth1-Cas9-L5-sgRNA-mmar_0499 | target gRNA: GGCAAGTACCGCCAGTGTGCGCGGAA | pCRISPRx-Sth1 Cas9-L5 (Addgene 140993) | This study |
| pCRISPRx-Sth1-Cas9-L5-sgRNA-mmar_0620 | target gRNA: GCGACGTCGTTGGTCGGCGCGTCGGAA | pCRISPRx-Sth1 Cas9-L5 (Addgene 140993) | This study |
| pCRISPRx-Sth1-Cas9-L5-sgRNA-mmar_0696 | target gRNA: ACACCACACCAGCAACTCGCCGCAGAA | pCRISPRx-Sth1 Cas9-L5 (Addgene 140993) | This study |
| pCRISPRx-Sth1-Cas9-L5-sgRNA-mmar_0726 | target gRNA: GTCCACCGATCCCTTTTCGGAATGGAA | pCRISPRx-Sth1 Cas9-L5 (Addgene 140993) | This study |
| pCRISPRx-Sth1-Cas9-L5-sgRNA-mmar_0728 | target gRNA: ATCATGTTGCAGCGGGAGGTGAAGAA | pCRISPRx-Sth1 Cas9-L5 (Addgene 140993) | This study |
| pCRISPRx-Sth1-Cas9-L5-sgRNA-mmar_0950 | target gRNA: ACATCCGGGACAACGACATCCAGGAA | pCRISPRx-Sth1 Cas9-L5 (Addgene 140993) | This study |
| pCRISPRx-Sth1-Cas9-L5-sgRNA-mmar_1000 | target gRNA: ACACCGCCAACATCGATCGTGTGGAA | pCRISPRx-Sth1 Cas9-L5 (Addgene 140993) | This study |
| pCRISPRx-Sth1-Cas9-L5-sgRNA-mmar_1362 | target gRNA: ACGCATTGATCATCAGCGTCGAAGAA | pCRISPRx-Sth1 Cas9-L5 (Addgene 140993) | This study |
| pCRISPRx-Sth1-Cas9-L5-sgRNA-mmar_4800 | target gRNA: GCGCAGGTCGATGATGGCGTCGGGAA | pCRISPRx-Sth1 Cas9-L5 (Addgene 140993) | This study |
| pCRISPRx-Sth1-Cas9-L5-sgRNA-mmar_4769 | target gRNA: ACCGATGTCGCCAAGGGGTGATCGGAA | pCRISPRx-Sth1 Cas9-L5 (Addgene 140993) | This study |
| pCRISPRx-Sth1-Cas9-L5-sgRNA-mmar_4470 | target gRNA: ACCTGGAAAGCATCGGGGTCAGCGGAA | pCRISPRx-Sth1 Cas9-L5 (Addgene 140993) | This study |
| pCRISPRx-Sth1-Cas9-L5-sgRNA-mmar_4463 | target gRNA: GCCGCTCGGGTTGCGGAGGTCGAGAA | pCRISPRx-Sth1 Cas9-L5 (Addgene 140993) | This study |
| pCRISPRx-Sth1-Cas9-L5-sgRNA-mmar_4402 | target gRNA: GTCCCCGCAGAGTCGACCGAAGAGGAA | pCRISPRx-Sth1 Cas9-L5 (Addgene 140993) | This study |
| pCRISPRx-Sth1-Cas9-L5-sgRNA-mmar_4288 | target gRNA: ACCATCTGCCGCCACAAGTACCAGAA | pCRISPRx-Sth1 Cas9-L5 (Addgene 140993) | This study |
| pCRISPRx-Sth1-Cas9-L5-sgRNA-mmar_4206 | target gRNA: GGTCCATTGAAGACCGCCACCTGGAA | pCRISPRx-Sth1 Cas9-L5 (Addgene 140993) | This study |

| | | | |
|---|---|---|---|
| pCRISPRx-Sth1-Cas9-L5-sgRNA-mmar_4196 | target gRNA: ATGGCCCGGTAGACCTCTTTGTCAGAA | pCRISPRx-Sth1 Cas9-L5 (Addgene 140993) | This study |
| pCRISPRx-Sth1-Cas9-L5-sgRNA-mmar_4152 | target gRNA: GCCGAGGCCATGCGCAAGGTCACCGGAA | pCRISPRx-Sth1 Cas9-L5 (Addgene 140993) | This study |
| pCRISPRx-Sth1-Cas9-L5-sgRNA-mmar_4146 | target gRNA: GTCGCACCGGGATTGTTGTCACCGGAA | pCRISPRx-Sth1 Cas9-L5 (Addgene 140993) | This study |
| pCRISPRx-Sth1-Cas9-L5-sgRNA-mmar_4145 | target gRNA: AGCAGTACCCCAACCTGCTCAAAGAA | pCRISPRx-Sth1 Cas9-L5 (Addgene 140993) | This study |
| pCRISPRx-Sth1-Cas9-L5-sgRNA-mmar_4038 | target gRNA: ACTGCAGCGCGTAGCCGAGATTTTGGAA | pCRISPRx-Sth1 Cas9-L5 (Addgene 140993) | This study |
| pCRISPRx-Sth1-Cas9-L5-sgRNA-mmar_4190 | target gRNA: ACTCGATCGTCGCCAAGACCCCAGAA | pCRISPRx-Sth1 Cas9-L5 (Addgene 140993) | This study |
| pCRISPRx-Sth1-Cas9-L5-sgRNA-mmar_2188 | target gRNA: ATTGCGACAGACGTCGGGGTCGAGAA | pCRISPRx-Sth1 Cas9-L5 (Addgene 140993) | This study |
| pCRISPRx-Sth1-Cas9-L5-sgRNA-mmar_2220 | target gRNA: AAGCAAACAACCGAGCTCACCAGGAA | pCRISPRx-Sth1 Cas9-L5 (Addgene 140993) | This study |
| pCRISPRx-Sth1-Cas9-L5-sgRNA-mmar_2225 | target gRNA: GCCCGAATCGCGCGACCGGCAACGGAA | pCRISPRx-Sth1 Cas9-L5 (Addgene 140993) | This study |
| pCRISPRx-Sth1-Cas9-L5-sgRNA-mmar_2495 | target gRNA: AACACCCCGGCGGTCAACGCCAGGAA | pCRISPRx-Sth1 Cas9-L5 (Addgene 140993) | This study |
| pCRISPRx-Sth1-Cas9-L5-sgRNA-mmar_0416 | target gRNA: GTCGATTCGCAACATCCAGCTCAAGAA | pCRISPRx-Sth1 Cas9-L5 (Addgene 140993) | This study |
| pCRISPRx-Sth1-Cas9-L5-sgRNA-mmar_2364 | target gRNA: GAGACGCGTGACCAATGTGCGCAGAA | pCRISPRx-Sth1 Cas9-L5 (Addgene 140993) | This study |
| pCRISPRx-Sth1-Cas9-L5-sgRNA-mmar_2886 | target gRNA: ACGTGACGGTCGGCCATGTGGCAAGAA | pCRISPRx-Sth1 Cas9-L5 (Addgene 140993) | This study |
| pCRISPRx-Sth1-Cas9-L5-sgRNA-mmar_4983 | target gRNA: AAGGCCAATTTGACCGCGGCATAGAA | pCRISPRx-Sth1 Cas9-L5 (Addgene 140993) | This study |
| pCRISPRx-Sth1-Cas9-L5-sgRNA-mmar_0422 | target gRNA: GGCGGGCATCTGATGATCGTCAAGAA | pCRISPRx-Sth1 Cas9-L5 (Addgene 140993) | This study |
| pCRISPRx-Sth1-Cas9-L5-sgRNA-mmar_0809 | target gRNA: AAGGTCGCCAAGCAGGCCGAGCAGAA | pCRISPRx-Sth1 Cas9-L5 (Addgene 140993) | This study |
| pCRISPRx-Sth1-Cas9-L5-sgRNA-mmar_2731 | target gRNA: GTGAGTTGGGTGGCCAACTCGGAGGAA | pCRISPRx-Sth1 Cas9-L5 (Addgene 140993) | This study |
| pCRISPRx-Sth1-Cas9-L5-sgRNA-mmar_1834 | target gRNA: GTTGCTGCAATTGCTGGGTTGTGGAA | pCRISPRx-Sth1 Cas9-L5 (Addgene 140993) | This study |
| pCRISPRx-Sth1-Cas9-L5-sgRNA-mmar_4139 | target gRNA: GTGATCGGCGTGCCATCAGACCAGAA | pCRISPRx-Sth1 Cas9-L5 (Addgene 140993) | This study |
| pCRISPRx-Sth1-Cas9-L5-sgRNA-mmar_5302 | target gRNA: GCTTCGAGGTCGGAATGCGGTTGGGAA | pCRISPRx-Sth1 Cas9-L5 (Addgene 140993) | This study |
| pCRISPRx-Sth1-Cas9-L5-sgRNA-mmar_4576 | target gRNA: GCCGGTGGATTGCGGCGGTAAGAAGAA | pCRISPRx-Sth1 Cas9-L5 (Addgene 140993) | This study |
| pCRISPRx-Sth1-Cas9-L5-sgRNA-mmar_4580 | target gRNA: AGCAACGTGAGCTGCGGGGGGAAGAA | pCRISPRx-Sth1 Cas9-L5 (Addgene 140993) | This study |
| pCRISPRx-Sth1-Cas9-L5-sgRNA-mmar_0525 | target gRNA: GGAGTCGACGCCGAGACCTATCAGAA | pCRISPRx-Sth1 Cas9-L5 (Addgene 140993) | This study |
| pCRISPRx-Sth1-Cas9-L5-sgRNA-mmar_2827 | target gRNA: GCCGAGCATCACCGCAATCACCAGAA | pCRISPRx-Sth1 Cas9-L5 (Addgene 140993) | This study |
| pCRISPRx-Sth1-Cas9-L5-sgRNA-mmar_3014 | target gRNA: AGAACGTCAGGGCGTCATCGTCGGAA | pCRISPRx-Sth1 Cas9-L5 (Addgene 140993) | This study |
| pCRISPRx-Sth1-Cas9-L5-sgRNA-mmar_2122 | target gRNA: AACATCGAATGCACGCCCGGGCAGAA | pCRISPRx-Sth1 Cas9-L5 (Addgene 140993) | This study |
| pCRISPRx-Sth1-Cas9-L5-sgRNA-mmar_1840 | target gRNA: ACCGCTTCACCTGGCACCTTCCCAAGAA | pCRISPRx-Sth1 Cas9-L5 (Addgene 140993) | This study |
| pCRISPRx-Sth1-Cas9-L5-sgRNA-mmar_3718 | target gRNA: GTCGAGCGATGTCCGGTTCGACCGAGAA | pCRISPRx-Sth1 Cas9-L5 (Addgene 140993) | This study |
| pCRISPRx-Sth1-Cas9-L5-sgRNA-mmar_1900 | target gRNA: GATTGACCCGGATGATTGGACCAGAA | pCRISPRx-Sth1 Cas9-L5 (Addgene 140993) | This study |
| pCRISPRx-Sth1-Cas9-L5-sgRNA-mmar_3619 | target gRNA: GCCAGGTAAGCGTTGGACAGCCAGAA | pCRISPRx-Sth1 Cas9-L5 (Addgene 140993) | This study |
| pCRISPRx-Sth1-Cas9-L5-sgRNA-mmar_3872 | target gRNA: ACCCCCACCGCGACCAAGGTGATCGAGAA | pCRISPRx-Sth1 Cas9-L5 (Addgene 140993) | This study |
| pCRISPRx-Sth1-Cas9-L5-sgRNA-mmar_5092 | target gRNA: GAGGTTGTGCTCGACGTTGCAGGGAA | pCRISPRx-Sth1 Cas9-L5 (Addgene 140993) | This study |
| pCRISPRx-Sth1-Cas9-L5-sgRNA-mmar_4479 | target gRNA: GACCGTGACATTGACCGTCGATGGAA | pCRISPRx-Sth1 Cas9-L5 (Addgene 140993) | This study |
| pCRISPRx-Sth1-Cas9-L5-sgRNA-mmar_3713 | target gRNA: GGTGCTGCTGTACTACGACGCCAGAA | pCRISPRx-Sth1 Cas9-L5 (Addgene 140993) | This study |
| pCRISPRx-Sth1-Cas9-L5-sgRNA-mmar_4200 | target gRNA: AGTCCGGGCCGCAGTGCTACCAGGAA | pCRISPRx-Sth1 Cas9-L5 (Addgene 140993) | This study |
| pCRISPRx-Sth1-Cas9-L5-sgRNA-mmar_3050 | target gRNA: ACGACTGGGGGTGTACGTGCCGGGAA | pCRISPRx-Sth1 Cas9-L5 (Addgene 140993) | This study |
| pCRISPRx-Sth1-Cas9-L5-sgRNA-mmar_1803 | target gRNA: ATGCCACCGCCGCCGCGGCGGTCAGCA | pCRISPRx-Sth1 Cas9-L5 (Addgene 140993) | This study |
| pCRISPRx-Sth1-Cas9-L5-sgRNA-mmar_1763 | target gRNA: GCCTCGGGTTGCGTTTTGTCAGAAGAA | pCRISPRx-Sth1 Cas9-L5 (Addgene 140993) | This study |
| pCRISPRx-Sth1-Cas9-L5-sgRNA-mmar_4146 | target gRNA: GCTGACACCGCCGCACTGCGGAAAGAA | pCRISPRx-Sth1 Cas9-L5 (Addgene 140993) | This study |
| pCRISPRx-Sth1-Cas9-L5-sgRNA-mmar_3718 | target gRNA: ACCAGCGTGACCACCGACTCGAACGGAA | pCRISPRx-Sth1 Cas9-L5 (Addgene 140993) | This study |
| pCRISPRx-Sth1-Cas9-L5-sgRNA-mmar_1667 | target gRNA: GGGTGTTCGCTGATGGAGTGGGAGAA | pCRISPRx-Sth1 Cas9-L5 (Addgene 140993) | This study |
| pLJR965-sgRNA-lpqW | target gRNA: GACCATCTGCCGCCACAAGTACCAGAA | pLJR965 (Addgene 115163) | This study |
| pLJR965-sgRNA-aftA | target gRNA: GCGCGTGCAGCTGGTTGGACGACGGAA | pLJR965 (Addgene 115163) | This study |
| pLJR965-sgRNA-aftB | target gRNA: GATCCGGACCCAGGGGTCATACGGAA | pLJR965 (Addgene 115163) | This study |
| pSMT3-empty | used as EV control | - | pSMT3-hsp60, [6] |
| pSMT3-lpqZ-HA | overexpression | pSMT3-lpqZ-HA (Addgene 240159) | This study |

| | | | |
|---|---|---|---|
| pSMT3-fecB-HA | overexpression for co-IP | pSMT3, mmar_1650 genomic DNA | This study |
| pSMT3-fecB | overexpression for complementation | pSMT3-fecB (Addgene 240158) | This study |
| pLJR962-AftA-FLAG | overexpression | pLJR962 (Addgene 115162), mmar_5354 genomic DNA | This study |
| pSMT3-MspA | overexpression of MspA porin of M. smegmatis | - | pSMT3-MspA,[4] |
| pML1357kana-mmar1667-Strep | kanaR, overexpression | pML1357-mmar_0407 [7], mmar_1667 genomic DNA | This study |
| pSMT3-fecB(MTB) | overexpression, M.tuberculosis gene | pSMT3, Rv3044 genomic DNA | This study |
| pSMT3-lpqZ(MTB)-HA | overexpression, M.tuberculosis gene | pSMT3, Rv1244 genomic DNA | This study |

**Supplementary Table 5: oligonucleotides used in this study.** Used oligonucleotides with their sequence and applied usage in this study.

| oligonucleotide name | sequence 5'-> 3' | usage |
|---|---|---|
| mmar_5154_gRNA_Fw | GGGAGACACGGTGGCCAACAGCAT | oligo for target sgRNA for pCRISPRx-Sth1-Cas9 L5 |
| mmar_5057_gRNA_Fw | GGGAGCACTTGGGGCACCACGGTGT | oligo for target sgRNA for pCRISPRx-Sth1-Cas9 L5 |
| mmar_1650_gRNA_Fw | GGGAGGCTGGCTATTGGAACCGTC | oligo for target sgRNA for pCRISPRx-Sth1-Cas9 L5 |
| mmar_0714_gRNA_Fw | GGGAGTCCTGGCTGGAGCTGTCCG | oligo for target sgRNA for pCRISPRx-Sth1-Cas9 L5 |
| mmar_2794_gRNA_Fw | GGGAAGCCGTGCCGCCCGTTCGAG | oligo for target sgRNA for pCRISPRx-Sth1-Cas9 L5 |
| mmar_2769_gRNA_Fw | GGGAGCACGCCCCGCCGCATGCAGT | oligo for target sgRNA for pCRISPRx-Sth1-Cas9 L5 |
| mmar_0949_gRNA_Fw | GGGAGACTGGGACAACACCGTCGT | oligo for target sgRNA for pCRISPRx-Sth1-Cas9 L5 |
| mmar_5074_gRNA_Fw | GGGAACTGGAGCAAGAACATCTCGGA | oligo for target sgRNA for pCRISPRx-Sth1-Cas9 L5 |
| mmar_3021_gRNA_Fw | GGGAGTGGGAAGTTTGCTAGTCGAG | oligo for target sgRNA for pCRISPRx-Sth1-Cas9 L5 |
| mmar_5467_gRNA_Fw | GGGAATCGCCTTGGTCATTGCATGA | oligo for target sgRNA for pCRISPRx-Sth1-Cas9 L5 |
| mmar_3092_gRNA_Fw | GGGAGATCCGAACATTCCGGGAGA | oligo for target sgRNA for pCRISPRx-Sth1-Cas9 L5 |
| mmar_3118_gRNA_Fw | GGGAGAACCACACCTGCCGGAGGC | oligo for target sgRNA for pCRISPRx-Sth1-Cas9 L5 |
| mmar_3206_gRNA_Fw | GGGAGATCATCGCGGCGGCAAAACC | oligo for target sgRNA for pCRISPRx-Sth1-Cas9 L5 |
| mmar_3366_gRNA_Fw | GGGAACCACCACTCATCAAGCGGGG | oligo for target sgRNA for pCRISPRx-Sth1-Cas9 L5 |
| mmar_3638_gRNA_Fw | GGGAACTACCGCGCCGGATACCTG | oligo for target sgRNA for pCRISPRx-Sth1-Cas9 L5 |
| mmar_3722_gRNA_Fw | GGGAACTTGGCCGTGAAGTCCTGGC | oligo for target sgRNA for pCRISPRx-Sth1-Cas9 L5 |
| mmar_1923_gRNA_Fw | GGGAGCCAGAGCCATCAGCACGGC | oligo for target sgRNA for pCRISPRx-Sth1-Cas9 L5 |
| mmar_1914_gRNA_Fw | GGGAGCTCACCACCACGCCCAAGACCA | oligo for target sgRNA for pCRISPRx-Sth1-Cas9 L5 |
| mmar_1803_gRNA_Fw | GGGAAACGCATCCAGCGGGCAACC | oligo for target sgRNA for pCRISPRx-Sth1-Cas9 L5 |
| mmar_1763_gRNA_Fw | GGGAGAGCCTCGGGTTGCGTTTTG | oligo for target sgRNA for pCRISPRx-Sth1-Cas9 L5 |
| mmar_5154_gRNA_Rv | AAACATGCTGTTGGCCACCGTGTC | oligo for target sgRNA for pCRISPRx-Sth1-Cas9 L5 |
| mmar_5057_gRNA_Rv | AAACACACCGTGGTGCCCCAAGTGC | oligo for target sgRNA for pCRISPRx-Sth1-Cas9 L5 |
| mmar_1650_gRNA_Rv | AAACGACGGTTCCAATAGCCAGCC | oligo for target sgRNA for pCRISPRx-Sth1-Cas9 L5 |
| mmar_0714_gRNA_Rv | AAACCGGACAGCTCCAGCCAGGAC | oligo for target sgRNA for pCRISPRx-Sth1-Cas9 L5 |
| mmar_2794_gRNA_Rv | AAACCTCGAACGGGCGGCACGGCT | oligo for target sgRNA for pCRISPRx-Sth1-Cas9 L5 |
| mmar_2769_gRNA_Rv | AAACACTGCATGCGGCGGGGCGTGC | oligo for target sgRNA for pCRISPRx-Sth1-Cas9 L5 |
| mmar_0949_gRNA_Rv | AAACACGACGGTGTTGTCCCAGTC | oligo for target sgRNA for pCRISPRx-Sth1-Cas9 L5 |
| mmar_5074_gRNA_Rv | AAACTCCGAGATGTTCTTGCTCCAGT | oligo for target sgRNA for pCRISPRx-Sth1-Cas9 L5 |
| mmar_3021_gRNA_Rv | AAACCTCGACTAGCAAACTTCCCAC | oligo for target sgRNA for pCRISPRx-Sth1-Cas9 L5 |
| mmar_5467_gRNA_Rv | AAACTCATGCAATGACCAAGGCGAT | oligo for target sgRNA for pCRISPRx-Sth1-Cas9 L5 |
| mmar_3092_gRNA_Rv | AAACTCTCCCGGAATGTTCGGATC | oligo for target sgRNA for pCRISPRx-Sth1-Cas9 L5 |
| mmar_3118_gRNA_Rv | AAACGCCTCCGGCAGGTGTGGTTC | oligo for target sgRNA for pCRISPRx-Sth1-Cas9 L5 |
| mmar_3206_gRNA_Rv | AAACGGTTTTGCCGCCGCGATGATC | oligo for target sgRNA for pCRISPRx-Sth1-Cas9 L5 |
| mmar_3366_gRNA_Rv | AAACCCCCGCTTGATGAGTGGTGGT | oligo for target sgRNA for pCRISPRx-Sth1-Cas9 L5 |
| mmar_3638_gRNA_Rv | AAACCAGGTATCCGGCGCGGTAGT | oligo for target sgRNA for pCRISPRx-Sth1-Cas9 L5 |
| mmar_3722_gRNA_Rv | AAACGCCAGGACTTCACGGCCAAGT | oligo for target sgRNA for pCRISPRx-Sth1-Cas9 L5 |
| mmar_1923_gRNA_Rv | AAACGCCGTGCTGATGGCTCTGGC | oligo for target sgRNA for pCRISPRx-Sth1-Cas9 L5 |
| mmar_1914_gRNA_Rv | AAACTGGTCTTGGGCGTGGTGGTGAGC | oligo for target sgRNA for pCRISPRx-Sth1-Cas9 L5 |
| mmar_1803_gRNA_Rv | AAACGGTTGCCCGCTGGATGCGTT | oligo for target sgRNA for pCRISPRx-Sth1-Cas9 L5 |
| mmar_1763_gRNA_Rv | AAACCAAAACGCAACCCGAGGCTC | oligo for target sgRNA for pCRISPRx-Sth1-Cas9 L5 |
| mmar_1706_gRNA_Fw | GGGAGTCAAGGAGGTGTCGGTCAGC | oligo for target sgRNA for pCRISPRx-Sth1-Cas9 L5 |
| mmar_1691_gRNA_Fw | GGGAGCCCTGGCAAAGACCGCGCCG | oligo for target sgRNA for pCRISPRx-Sth1-Cas9 L5 |
| mmar_1301_gRNA_Fw | GGGAATGGATCCCGACGTGCTGCTG | oligo for target sgRNA for pCRISPRx-Sth1-Cas9 L5 |
| mmar_1236_gRNA_Fw | GGGAGTACCTTGCCCACTAGGGCCA | oligo for target sgRNA for pCRISPRx-Sth1-Cas9 L5 |
| mmar_1154_gRNA_Fw | GGGAGATCATCGCTACCGAGTTGC | oligo for target sgRNA for pCRISPRx-Sth1-Cas9 L5 |

| | | |
|---|---|---|
| mmar_5084_gRNA_Fw | GGGAGTCCACATTGTGGCCGGGACG | oligo for target sgRNA for pCRISPRx-Sth1-Cas9 L5 |
| mmar_5092_gRNA_Fw | GGGAGCCAGGCGGTGATCGTCGGTG | oligo for target sgRNA for pCRISPRx-Sth1-Cas9 L5 |
| mmar_5123_gRNA_Fw | GGGAGCTCATCGCGCCGGTGACGTCC | oligo for target sgRNA for pCRISPRx-Sth1-Cas9 L5 |
| mmar_5315_gRNA_Fw | GGGAACGTCAACATCGCGATCGGT | oligo for target sgRNA for pCRISPRx-Sth1-Cas9 L5 |
| mmar_0499_gRNA_Fw | GGGAGGCAAGTACCGCCAGTGTGC | oligo for target sgRNA for pCRISPRx-Sth1-Cas9 L5 |
| mmar_0620_gRNA_Fw | GGGAGCGACGTCGTTGGTCGGCGCG | oligo for target sgRNA for pCRISPRx-Sth1-Cas9 L5 |
| mmar_0696_gRNA_Fw | GGGAACACCACACCAGCAACTCGCC | oligo for target sgRNA for pCRISPRx-Sth1-Cas9 L5 |
| mmar_0726_gRNA_Fw | GGGAGTCCACCGATCCCTTTTCGGA | oligo for target sgRNA for pCRISPRx-Sth1-Cas9 L5 |
| mmar_0728_gRNA_Fw | GGGAATCATGTTGCAGCGGGAGGT | oligo for target sgRNA for pCRISPRx-Sth1-Cas9 L5 |
| mmar_0950_gRNA_Fw | GGGAACATCCGGGACAACGACATC | oligo for target sgRNA for pCRISPRx-Sth1-Cas9 L5 |
| mmar_1000_gRNA_Fw | GGGAACACCGCCAACATCGATCGT | oligo for target sgRNA for pCRISPRx-Sth1-Cas9 L5 |
| mmar_1362_gRNA_Fw | GGGAACGCATTGATCATCAGCGTC | oligo for target sgRNA for pCRISPRx-Sth1-Cas9 L5 |
| mmar_4800_gRNA_Fw | GGGAGCGCAGGTCGATGATGGCGT | oligo for target sgRNA for pCRISPRx-Sth1-Cas9 L5 |
| mmar_4769_gRNA_Fw | GGGAACCGATGTCGCCAAGGGGTGA | oligo for target sgRNA for pCRISPRx-Sth1-Cas9 L5 |
| mmar_4470_gRNA_Fw | GGGAACCTGGAAAGCATCGGGGTCA | oligo for target sgRNA for pCRISPRx-Sth1-Cas9 L5 |
| mmar_4463_gRNA_Fw | GGGAGCCGCTCGGGTTGCGGAGGT | oligo for target sgRNA for pCRISPRx-Sth1-Cas9 L5 |
| mmar_4402_gRNA_Fw | GGGAGTCCCCGCAGAGTCGACCGAA | oligo for target sgRNA for pCRISPRx-Sth1-Cas9 L5 |
| mmar_4288_gRNA_Fw | GGGAACCATCTGCCGCCACAAGTA | oligo for target sgRNA for pCRISPRx-Sth1-Cas9 L5 |
| mmar_4206_gRNA_Fw | GGGAGGTCCATTGAAGACCGCCAC | oligo for target sgRNA for pCRISPRx-Sth1-Cas9 L5 |
| mmar_4196_gRNA_Fw | GGGAATGGCCCGGTAGACCTCTTTG | oligo for target sgRNA for pCRISPRx-Sth1-Cas9 L5 |
| mmar_4152_gRNA_Fw | GGGAGCCGAGGCCATGCGCAAGGTCA | oligo for target sgRNA for pCRISPRx-Sth1-Cas9 L5 |
| mmar_4146_gRNA_Fw | GGGAGTCGCACCGGGATTGTTGTCA | oligo for target sgRNA for pCRISPRx-Sth1-Cas9 L5 |
| mmar_4145_gRNA_Fw | GGGAAGCAGTACCCCAACCTGCTC | oligo for target sgRNA for pCRISPRx-Sth1-Cas9 L5 |
| mmar_4038_gRNA_Fw | GGGAACTGCAGCGCGTAGCCGAGATT | oligo for target sgRNA for pCRISPRx-Sth1-Cas9 L5 |
| mmar_4190_gRNA_Fw | GGGAACTCGATCGTCGCCAAGACC | oligo for target sgRNA for pCRISPRx-Sth1-Cas9 L5 |
| mmar_1706_gRNA_Rv | AAACGCTGACCGACACCTCCTTGAC | oligo for target sgRNA for pCRISPRx-Sth1-Cas9 L5 |
| mmar_1691_gRNA_Rv | AAACCGGCGCGGTCTTTGCCAGGGC | oligo for target sgRNA for pCRISPRx-Sth1-Cas9 L5 |
| mmar_1301_gRNA_Rv | AAACCAGCAGCACGTCGGGATCCAT | oligo for target sgRNA for pCRISPRx-Sth1-Cas9 L5 |
| mmar_1236_gRNA_Rv | AAACTGGCCCTAGTGGGCAAGGTAC | oligo for target sgRNA for pCRISPRx-Sth1-Cas9 L5 |
| mmar_1154_gRNA_Rv | AAACGCAACTCGGTAGCGATGATC | oligo for target sgRNA for pCRISPRx-Sth1-Cas9 L5 |
| mmar_5084_gRNA_Rv | AAACCGTCCCGGCCACAATGTGGAC | oligo for target sgRNA for pCRISPRx-Sth1-Cas9 L5 |
| mmar_5092_gRNA_Rv | AAACCACCGACGATCACCGCCTGGC | oligo for target sgRNA for pCRISPRx-Sth1-Cas9 L5 |
| mmar_5123_gRNA_Rv | AAACGGACGTCACCGGCGCGATGAGC | oligo for target sgRNA for pCRISPRx-Sth1-Cas9 L5 |
| mmar_5315_gRNA_Rv | AAACACCGATCGCGATGTTGACGT | oligo for target sgRNA for pCRISPRx-Sth1-Cas9 L5 |
| mmar_0499_gRNA_Rv | AAACGCACACTGGCGGTACTTGCC | oligo for target sgRNA for pCRISPRx-Sth1-Cas9 L5 |
| mmar_0620_gRNA_Rv | AAACCGCGCCGACCAACGACGTCGC | oligo for target sgRNA for pCRISPRx-Sth1-Cas9 L5 |
| mmar_0696_gRNA_Rv | AAACGGCGAGTTGCTGGTGTGGTGT | oligo for target sgRNA for pCRISPRx-Sth1-Cas9 L5 |
| mmar_0726_gRNA_Rv | AAACTCCGAAAAGGGATCGGTGGAC | oligo for target sgRNA for pCRISPRx-Sth1-Cas9 L5 |
| mmar_0728_gRNA_Rv | AAACACCTCCCGCTGCAACATGAT | oligo for target sgRNA for pCRISPRx-Sth1-Cas9 L5 |
| mmar_0950_gRNA_Rv | AAACGATGTCGTTGTCCCGGATGT | oligo for target sgRNA for pCRISPRx-Sth1-Cas9 L5 |
| mmar_1000_gRNA_Rv | AAACACGATCGATGTTGGCGGTGT | oligo for target sgRNA for pCRISPRx-Sth1-Cas9 L5 |
| mmar_1362_gRNA_Rv | AAACGACGCTGATGATCAATGCGT | oligo for target sgRNA for pCRISPRx-Sth1-Cas9 L5 |
| mmar_4800_gRNA_Rv | AAACACGCCATCATCGACCTGCGC | oligo for target sgRNA for pCRISPRx-Sth1-Cas9 L5 |
| mmar_4769_gRNA_Rv | AAACTCACCCCTTGGCGACATCGGT | oligo for target sgRNA for pCRISPRx-Sth1-Cas9 L5 |
| mmar_4470_gRNA_Rv | AAACTGACCCCGATGCTTTCCAGGT | oligo for target sgRNA for pCRISPRx-Sth1-Cas9 L5 |
| mmar_4463_gRNA_Rv | AAACACCTCCGCAACCCGAGCGGC | oligo for target sgRNA for pCRISPRx-Sth1-Cas9 L5 |

| | | |
|---|---|---|
| mmar_4402_gRNA_Rv | AAACTTCGGTCGACTCTGCGGGGAC | oligo for target sgRNA for pCRISPRx-Sth1-Cas9 L5 |
| mmar_4288_gRNA_Rv | AAACTACTTGTGGCGGCAGATGGT | oligo for target sgRNA for pCRISPRx-Sth1-Cas9 L5 |
| mmar_4206_gRNA_Rv | AAACGTGGCGGTCTTCAATGGACC | oligo for target sgRNA for pCRISPRx-Sth1-Cas9 L5 |
| mmar_4196_gRNA_Rv | AAACCAAAGAGGTCTACCGGGCCAT | oligo for target sgRNA for pCRISPRx-Sth1-Cas9 L5 |
| mmar_4152_gRNA_Rv | AAACTGACCTTGCGCATGGCCTCGGC | oligo for target sgRNA for pCRISPRx-Sth1-Cas9 L5 |
| mmar_4146_gRNA_Rv | AAACTGACAACAATCCCGGTGCGAC | oligo for target sgRNA for pCRISPRx-Sth1-Cas9 L5 |
| mmar_4145_gRNA_Rv | AAACGAGCAGGTTGGGGTACTGCT | oligo for target sgRNA for pCRISPRx-Sth1-Cas9 L5 |
| mmar_4038_gRNA_Rv | AAACAATCTCGGCTACGCGCTGCAGT | oligo for target sgRNA for pCRISPRx-Sth1-Cas9 L5 |
| mmar_4190_gRNA_Rv | AAACGGTCTTGGCGACGATCGAGT | oligo for target sgRNA for pCRISPRx-Sth1-Cas9 L5 |
| mmar_2188_gRNA_Fw | GGGAATTGCGACAGACGTCGGGGT | oligo for target sgRNA for pCRISPRx-Sth1-Cas9 L5 |
| mmar_2220_gRNA_Fw | GGGAAAGCAAACAACCGAGCTCAC | oligo for target sgRNA for pCRISPRx-Sth1-Cas9 L5 |
| mmar_2225_gRNA_Fw | GGGAGCCCGAATCGCGCGACCGGCA | oligo for target sgRNA for pCRISPRx-Sth1-Cas9 L5 |
| mmar_2495_gRNA_Fw | GGGAAACACCCCGGCGGTCAACGC | oligo for target sgRNA for pCRISPRx-Sth1-Cas9 L5 |
| mmar_0416_gRNA_Fw | GGGAGTCGATTCGCAACATCCAGCT | oligo for target sgRNA for pCRISPRx-Sth1-Cas9 L5 |
| mmar_2364_gRNA_Fw | GGGAGAGACGCGTGACCAATGTGC | oligo for target sgRNA for pCRISPRx-Sth1-Cas9 L5 |
| mmar_2886_gRNA_Fw | GGGAACGTGACGGTCGGCCATGTGG | oligo for target sgRNA for pCRISPRx-Sth1-Cas9 L5 |
| mmar_4983_gRNA_Fw | GGGAAAGGCCAATTTGACCGCGGC | oligo for target sgRNA for pCRISPRx-Sth1-Cas9 L5 |
| mmar_0422_gRNA_Fw | GGGAGGCGGGCATCTGATGATCGT | oligo for target sgRNA for pCRISPRx-Sth1-Cas9 L5 |
| mmar_0809_gRNA_Fw | GGGAAAGGTCGCCAAGCAGGCCGA | oligo for target sgRNA for pCRISPRx-Sth1-Cas9 L5 |
| mmar_2731_gRNA_Fw | GGGAGTGAGTTGGGTGGCCAACTCG | oligo for target sgRNA for pCRISPRx-Sth1-Cas9 L5 |
| mmar_1834_gRNA_Fw | GGGAGTTGCTGCAATTGCTGGGTT | oligo for target sgRNA for pCRISPRx-Sth1-Cas9 L5 |
| mmar_4139_gRNA_Fw | GGGAGTGATCGGCGTGCCATCAGA | oligo for target sgRNA for pCRISPRx-Sth1-Cas9 L5 |
| mmar_5302_gRNA_Fw | GGGAGCTTCGAGGTCGGAATGCGGT | oligo for target sgRNA for pCRISPRx-Sth1-Cas9 L5 |
| mmar_4576_gRNA_Fw | GGGAGCCGGTGGATTGCGGCGGTAA | oligo for target sgRNA for pCRISPRx-Sth1-Cas9 L5 |
| mmar_4580_gRNA_Fw | GGGAAGCAACGTGAGCTGCGGGGG | oligo for target sgRNA for pCRISPRx-Sth1-Cas9 L5 |
| mmar_0525_gRNA_Fw | GGGAGGAGTCGACGCCGAGACCTA | oligo for target sgRNA for pCRISPRx-Sth1-Cas9 L5 |
| mmar_2827_gRNA_Fw | GGGAGCCGAGCATCACCGCAATCA | oligo for target sgRNA for pCRISPRx-Sth1-Cas9 L5 |
| mmar_3014_gRNA_Fw | GGGAAGAACGTCAGGGCGTCATCG | oligo for target sgRNA for pCRISPRx-Sth1-Cas9 L5 |
| mmar_2122_gRNA_Fw | GGGAAACATCGAATGCACGCCCGG | oligo for target sgRNA for pCRISPRx-Sth1-Cas9 L5 |
| mmar_1840_gRNA_Fw | GGGAACCGCTTCACCTGGCACCTTCC | oligo for target sgRNA for pCRISPRx-Sth1-Cas9 L5 |
| mmar_3718_gRNA_Fw | GGGAGTCGAGCGATGTCCGGTTCGAC | oligo for target sgRNA for pCRISPRx-Sth1-Cas9 L5 |
| mmar_1900_gRNA_Fw | GGGAGATTGACCCGGATGATTGGA | oligo for target sgRNA for pCRISPRx-Sth1-Cas9 L5 |
| mmar_3619_gRNA_Fw | GGGAGCCAGGTAAGCGTTGGACAG | oligo for target sgRNA for pCRISPRx-Sth1-Cas9 L5 |
| mmar_3872_gRNA_Fw | GGGAACCCCCACCGCGACCAAGGTGAT | oligo for target sgRNA for pCRISPRx-Sth1-Cas9 L5 |
| mmar_2188_gRNA_Rv | AAACACCCCGACGTCTGTCGCAAT | oligo for target sgRNA for pCRISPRx-Sth1-Cas9 L5 |
| mmar_2220_gRNA_Rv | AAACGTGAGCTCGGTTGTTTGCTT | oligo for target sgRNA for pCRISPRx-Sth1-Cas9 L5 |
| mmar_2225_gRNA_Rv | AAACTGCCGGTCGCGCGATTCGGGC | oligo for target sgRNA for pCRISPRx-Sth1-Cas9 L5 |
| mmar_2495_gRNA_Rv | AAACGCGTTGACCGCCGGGGTGTT | oligo for target sgRNA for pCRISPRx-Sth1-Cas9 L5 |
| mmar_0416_gRNA_Rv | AAACAGCTGGATGTTGCGAATCGAC | oligo for target sgRNA for pCRISPRx-Sth1-Cas9 L5 |
| mmar_2364_gRNA_Rv | AAACGCACATTGGTCACGCGTCTC | oligo for target sgRNA for pCRISPRx-Sth1-Cas9 L5 |
| mmar_2886_gRNA_Rv | AAACCCACATGGCCGACCGTCACGT | oligo for target sgRNA for pCRISPRx-Sth1-Cas9 L5 |
| mmar_4983_gRNA_Rv | AAACGCCGCGGTCAAATTGGCCTT | oligo for target sgRNA for pCRISPRx-Sth1-Cas9 L5 |
| mmar_0422_gRNA_Rv | AAACACGATCATCAGATGCCCGCC | oligo for target sgRNA for pCRISPRx-Sth1-Cas9 L5 |
| mmar_0809_gRNA_Rv | AAACTCGGCCTGCTTGGCGACCTT | oligo for target sgRNA for pCRISPRx-Sth1-Cas9 L5 |
| mmar_2731_gRNA_Rv | AAACCGAGTTGGCCACCCAACTCAC | oligo for target sgRNA for pCRISPRx-Sth1-Cas9 L5 |
| mmar_1834_gRNA_Rv | AAACAACCCAGCAATTGCAGCAAC | oligo for target sgRNA for pCRISPRx-Sth1-Cas9 L5 |

| | | |
|---|---|---|
| mmar_4139_gRNA_Rv | AAACTCTGATGGCACGCCGATCAC | oligo for target sgRNA for pCRISPRx-Sth1-Cas9 L5 |
| mmar_5302_gRNA_Rv | AAACACCGCATTCCGACCTCGAAGC | oligo for target sgRNA for pCRISPRx-Sth1-Cas9 L5 |
| mmar_4576_gRNA_Rv | AAACTTACCGCCGCAATCCACCGGC | oligo for target sgRNA for pCRISPRx-Sth1-Cas9 L5 |
| mmar_4580_gRNA_Rv | AAACCCCCCGCAGCTCACGTTGCT | oligo for target sgRNA for pCRISPRx-Sth1-Cas9 L5 |
| mmar_0525_gRNA_Rv | AAACTAGGTCTCGGCGTCGACTCC | oligo for target sgRNA for pCRISPRx-Sth1-Cas9 L5 |
| mmar_2827_gRNA_Rv | AAACTGATTGCGGTGATGCTCGGC | oligo for target sgRNA for pCRISPRx-Sth1-Cas9 L5 |
| mmar_3014_gRNA_Rv | AAACCGATGACGCCCTGACGTTCT | oligo for target sgRNA for pCRISPRx-Sth1-Cas9 L5 |
| mmar_2122_gRNA_Rv | AAACCCGGGCGTGCATTCGATGTT | oligo for target sgRNA for pCRISPRx-Sth1-Cas9 L5 |
| mmar_1840_gRNA_Rv | AAACGGAAGGTGCCAGGTGAAGCGGT | oligo for target sgRNA for pCRISPRx-Sth1-Cas9 L5 |
| mmar_3718_gRNA_Rv | AAACGTCGAACCGGACATCGCTCGAC | oligo for target sgRNA for pCRISPRx-Sth1-Cas9 L5 |
| mmar_1900_gRNA_Rv | AAACTCCAATCATCCGGGTCAATC | oligo for target sgRNA for pCRISPRx-Sth1-Cas9 L5 |
| mmar_3619_gRNA_Rv | AAACCTGTCCAACGCTTACCTGGC | oligo for target sgRNA for pCRISPRx-Sth1-Cas9 L5 |
| mmar_3872_gRNA_Rv | AAACATCACCTTGGTCGCGGTGGGGGT | oligo for target sgRNA for pCRISPRx-Sth1-Cas9 L5 |
| mmar_4479_gRNA_Fw | GGGAGACCGTGACATTGACCGTCG | oligo for target sgRNA for pCRISPRx-Sth1-Cas9 L5 |
| mmar_3713_gRNA_Fw | GGGAGGTGCTGCTGTACTACGACG | oligo for target sgRNA for pCRISPRx-Sth1-Cas9 L5 |
| mmar_4200_gRNA_Fw | GGGAAGTCCGGGCCGCAGTGCTAC | oligo for target sgRNA for pCRISPRx-Sth1-Cas9 L5 |
| mmar_3050_gRNA_Fw | GGGAACGACTGGGGGTGTACGTGC | oligo for target sgRNA for pCRISPRx-Sth1-Cas9 L5 |
| mmar_4479_gRNA_Rv | AAACCGACGGTCAATGTCACGGTC | oligo for target sgRNA for pCRISPRx-Sth1-Cas9 L5 |
| mmar_3713_gRNA_Rv | AAACCGTCGTAGTACAGCAGCACC | oligo for target sgRNA for pCRISPRx-Sth1-Cas9 L5 |
| mmar_4200_gRNA_Rv | AAACGTAGCACTGCGGCCCGGACT | oligo for target sgRNA for pCRISPRx-Sth1-Cas9 L5 |
| mmar_3050_gRNA_Rv | AAACGCACGTACACCCCCAGTCGT | oligo for target sgRNA for pCRISPRx-Sth1-Cas9 L5 |
| mmar_1803_gRNA_Fw | GGGAATGCCACCGCCGCCGCGGCGG | oligo for target sgRNA for pCRISPRx-Sth1-Cas9 L5 |
| mmar_4146_gRNA_Fw | GGGAGCTGACACCGCCGCACTGCGG | oligo for target sgRNA for pCRISPRx-Sth1-Cas9 L5 |
| mmar_3718_gRNA_Fw | GGGAACCAGCGTGACCACCGACTCGA | oligo for target sgRNA for pCRISPRx-Sth1-Cas9 L5 |
| mmar_1803_gRNA_Rv | AAACCCGCCGCGGCGGCGGTGGCAT | oligo for target sgRNA for pCRISPRx-Sth1-Cas9 L5 |
| mmar_4146_gRNA_Rv | AAACCCGCAGTGCGGCGGTGTCAGC | oligo for target sgRNA for pCRISPRx-Sth1-Cas9 L5 |
| mmar_3718_gRNA_Rv | AAACTCGAGTCGGTGGTCACGCTGGT | oligo for target sgRNA for pCRISPRx-Sth1-Cas9 L5 |
| mmar_1667_gRNA_Fw | GGGAGGGTGTTCGCTGATGGAGTG | oligo for target sgRNA for pCRISPRx-Sth1-Cas9 L5 |
| mmar_1667_gRNA_Rv | AAACCACTCCATCAGCGAACACCC | oligo for target sgRNA for pCRISPRx-Sth1-Cas9 L5 |
| mmar_4288_gRNA_Fw | GGGAGACCATCTGCCGCCACAAGTA | oligo for target sgRNA for pLJR965 |
| mmar_4288_gRNA_Rv | AAACTACTTGTGGCGGCAGATGGTC | oligo for target sgRNA for pLJR965 |
| mmar_5354_gRNA_Fw | GGGAGCGCGTGCAGCTGGTTGGACG | oligo for target sgRNA for pLJR965 |
| mmar_5354_gRNA_Rv | AAACCGTCCAACCAGCTGCACGCGC | oligo for target sgRNA for pLJR965 |
| mmar_5369_gRNA_Fw | GGGAGATCCGGACCCAGGGGTCAT | oligo for target sgRNA for pLJR965 |
| mmar_5369_gRNA_Rv | AAACATGACCCCTGGGTCCGGATC | oligo for target sgRNA for pLJR965 |
| mmar_5154_LEFT | CTACGCCATCGCCTACAAC | genomic sequencing primer for sgRNA target |
| mmar_5057_LEFT | GTTCACCACCAAAACCATTGAT | genomic sequencing primer for sgRNA target |
| mmar_1650_LEFT | GATGTCGCCCAGATCTCC | genomic sequencing primer for sgRNA target |
| mmar_0714_LEFT | GAGCCGCTGAAAGTCGAG | genomic sequencing primer for sgRNA target |
| mmar_2794_LEFT | CCTACCTCGTCCGAGATCAT | genomic sequencing primer for sgRNA target |
| mmar_2769_LEFT | CAGATTCGGTTGGTCACTGTC | genomic sequencing primer for sgRNA target |
| mmar_0949_LEFT | GTCTCTTGAATGGCATGACG | genomic sequencing primer for sgRNA target |
| mmar_5074_LEFT | GACGTCAACCAGATCGACAG | genomic sequencing primer for sgRNA target |
| mmar_3021_LEFT | GTACCACTTGAAATCCCCTCAG | genomic sequencing primer for sgRNA target |
| mmar_5467_LEFT | GGGAGGATGTTCACTGATGAGA | genomic sequencing primer for sgRNA target |
| mmar_3118_LEFT | CGACATTCAGAGGGTAGACGTT | genomic sequencing primer for sgRNA target |

| | | |
|---|---|---|
| mmar_3206_LEFT | CAATGCTGTTGATGCTGGTG | genomic sequencing primer for sgRNA target |
| mmar_3366_LEFT | CTAGCCGATGCTTTTCAGGAG | genomic sequencing primer for sgRNA target |
| mmar_3638_LEFT | ACAGACTTCGCTAAGCTCGC | genomic sequencing primer for sgRNA target |
| mmar_3722_LEFT | TGTTGATCATCAGTCTGGCTTC | genomic sequencing primer for sgRNA target |
| mmar_1923_LEFT | CGGCTGATCAAGTACATCTCC | genomic sequencing primer for sgRNA target |
| mmar_1914_LEFT | CTGCCGGACAATAAGGTCG | genomic sequencing primer for sgRNA target |
| mmar_5154_RIGHT | AAACTCCTGGTTGGAATAGCC | genomic sequencing primer for sgRNA target |
| mmar_5057_RIGHT | GCTGTGTGACACCGAAGTTG | genomic sequencing primer for sgRNA target |
| mmar_1650_RIGHT | GCATACAAATTCGGTGTCAGC | genomic sequencing primer for sgRNA target |
| mmar_0714_RIGHT | AAACTGAACAGGTTGCTGCC | genomic sequencing primer for sgRNA target |
| mmar_2794_RIGHT | TAGAGCGTTACCTAGCTCAGCC | genomic sequencing primer for sgRNA target |
| mmar_2769_RIGHT | GTCAATATTGCGGATCTTCACC | genomic sequencing primer for sgRNA target |
| mmar_0949_RIGHT | GTGTTGTGCCAATCCTGGTT | genomic sequencing primer for sgRNA target |
| mmar_5074_RIGHT | TACCGAGGAGTTGTAGAAAGCG | genomic sequencing primer for sgRNA target |
| mmar_3021_RIGHT | CTCTATCCAGGCAATGACCTC | genomic sequencing primer for sgRNA target |
| mmar_5467_RIGHT | GCGATCTGTTGAAAGACCGTAT | genomic sequencing primer for sgRNA target |
| mmar_3118_RIGHT | CTCCTGAGTGAACACGCTGAT | genomic sequencing primer for sgRNA target |
| mmar_3206_RIGHT | GTCGCTGTCGTAGTTGGAGAC | genomic sequencing primer for sgRNA target |
| mmar_3366_RIGHT | CAGGTCAATCCATTTCTGTGTG | genomic sequencing primer for sgRNA target |
| mmar_3638_RIGHT | GGATTATCTGCACCCAGTACAAG | genomic sequencing primer for sgRNA target |
| mmar_3722_RIGHT | CTGATAGGCGCTCTTGGC | genomic sequencing primer for sgRNA target |
| mmar_1923_RIGHT | CAACGACCTTGAAGTTGGAGTC | genomic sequencing primer for sgRNA target |
| mmar_1914_RIGHT | CGCGGTACTGATCCTTGG | genomic sequencing primer for sgRNA target |
| mmar_1706_LEFT | CACCGTTCCCCAAGGAAT | genomic sequencing primer for sgRNA target |
| mmar_1691_LEFT | CCTGCTATTGGACCAAACCC | genomic sequencing primer for sgRNA target |
| mmar_1301_LEFT | GAGTTCATCAGCTCCACAAGC | genomic sequencing primer for sgRNA target |
| mmar_1236_LEFT | AACAACTTGCTGGTGGTGTATC | genomic sequencing primer for sgRNA target |
| mmar_1154_LEFT | CAAAGTCCTAGCTCTACTCGCC | genomic sequencing primer for sgRNA target |
| mmar_5084_LEFT | AACAGGCTCACCATCAACAAC | genomic sequencing primer for sgRNA target |
| mmar_5092_LEFT | GCACCTGACCTCCTCTTACG | genomic sequencing primer for sgRNA target |
| mmar_5123_LEFT | CACGTCAGGTGACCGTGTT | genomic sequencing primer for sgRNA target |
| mmar_5315_LEFT | CGGCACAAAGGTCCTCAT | genomic sequencing primer for sgRNA target |
| mmar_0499_LEFT | ATACAGCTCGTCCCTCTAGCC | genomic sequencing primer for sgRNA target |
| mmar_0620_LEFT | GATTTCTCGGGGAGTAACGC | genomic sequencing primer for sgRNA target |
| mmar_0696_LEFT | CCAAGGACTTGGGTATCTGAAT | genomic sequencing primer for sgRNA target |
| mmar_0726_LEFT | GACTATGTGGTCAACATCCTGC | genomic sequencing primer for sgRNA target |
| mmar_0728_LEFT | CAGGACTACTGGAAAGCCAACT | genomic sequencing primer for sgRNA target |
| mmar_0950_LEFT | GTGACAACAAGACCTCCACATC | genomic sequencing primer for sgRNA target |
| mmar_1000_LEFT | CCATTTGTTCACGTTACTGGC | genomic sequencing primer for sgRNA target |
| mmar_1362_LEFT | GGCTTTCATCGTCTGCATTG | genomic sequencing primer for sgRNA target |
| mmar_4800_LEFT | GTGTGTGGACGAGTCCCG | genomic sequencing primer for sgRNA target |
| mmar_4769_LEFT | ATATAGTGAGCCGCATGAAAGG | genomic sequencing primer for sgRNA target |
| mmar_4470_LEFT | CTGACTACAAATCCATCTGGACC | genomic sequencing primer for sgRNA target |
| mmar_4463_LEFT | GCAGCTCAGCTTGTTCATCC | genomic sequencing primer for sgRNA target |
| mmar_4402_LEFT | GTGTGCGCACCACTCATC | genomic sequencing primer for sgRNA target |
| mmar_4288_LEFT | TCACGGTCACCTACAAGATCC | genomic sequencing primer for sgRNA target |

| | | |
|---|---|---|
| mmar_4206_LEFT | GCTTGGACGTGATGTGGAC | genomic sequencing primer for sgRNA target |
| mmar_4196_LEFT | AATTGGACTCCGGTGACTTTAC | genomic sequencing primer for sgRNA target |
| mmar_4152_LEFT | CCAACATAAGGAGAATTCCGAT | genomic sequencing primer for sgRNA target |
| mmar_4145_LEFT | ATTGCGAACTCGAAATGAGC | genomic sequencing primer for sgRNA target |
| mmar_4038_LEFT | GGTACCGTCAAGGGGTGTC | genomic sequencing primer for sgRNA target |
| mmar_4190_LEFT | GTGTGAAGTCGCTCCCGT | genomic sequencing primer for sgRNA target |
| mmar_1706_RIGHT | GAGAGCACGATATCCATCAAGC | genomic sequencing primer for sgRNA target |
| mmar_1691_RIGHT | GATCTTGGAAGGTGGTCTTGTG | genomic sequencing primer for sgRNA target |
| mmar_1301_RIGHT | GGGTTTCCACGAAGACCAC | genomic sequencing primer for sgRNA target |
| mmar_1236_RIGHT | AGGCGGTTGGACATGAATC | genomic sequencing primer for sgRNA target |
| mmar_1154_RIGHT | CGATGTGTAGATGCTGTCGAAT | genomic sequencing primer for sgRNA target |
| mmar_5084_RIGHT | CGATTCATTCGAGTCCAGC | genomic sequencing primer for sgRNA target |
| mmar_5092_RIGHT | GTAGAGGTTGGTTGGTGTTGGT | genomic sequencing primer for sgRNA target |
| mmar_5123_RIGHT | TCTTGACTTCGATCGCATTATC | genomic sequencing primer for sgRNA target |
| mmar_5315_RIGHT | CGCTGATCTTGTAGCTGCTG | genomic sequencing primer for sgRNA target |
| mmar_0499_RIGHT | GAGGTCGTGGTCGAGGAG | genomic sequencing primer for sgRNA target |
| mmar_0620_RIGHT | GATGAAGACATAACCGTTGTCG | genomic sequencing primer for sgRNA target |
| mmar_0696_RIGHT | ACCTTCATCACCGCATCTG | genomic sequencing primer for sgRNA target |
| mmar_0726_RIGHT | GTTGTCGTAGTCGGAGGGAGT | genomic sequencing primer for sgRNA target |
| mmar_0728_RIGHT | CACCGAGATCTCACCGAAAT | genomic sequencing primer for sgRNA target |
| mmar_0950_RIGHT | AGCCTTCGGGTAGCAGTTG | genomic sequencing primer for sgRNA target |
| mmar_1000_RIGHT | ATAGACGACCACGAACTTCTCC | genomic sequencing primer for sgRNA target |
| mmar_1362_RIGHT | CAAAGGTGAGGTCACTGGTTC | genomic sequencing primer for sgRNA target |
| mmar_4800_RIGHT | ATTCGTGCACCAGACATCG | genomic sequencing primer for sgRNA target |
| mmar_4769_RIGHT | ATGGAGGGGTCGAATTGTC | genomic sequencing primer for sgRNA target |
| mmar_4470_RIGHT | CTTGGAGATGATCACCGTTTC | genomic sequencing primer for sgRNA target |
| mmar_4463_RIGHT | GATAGGTGCCGTTGTGACTTTC | genomic sequencing primer for sgRNA target |
| mmar_4402_RIGHT | CATGGCCAGGTAGGGCTC | genomic sequencing primer for sgRNA target |
| mmar_4288_RIGHT | GCAGGTTGTCAAACAGCTCTC | genomic sequencing primer for sgRNA target |
| mmar_4206_RIGHT | CGATCCAGGTTGGCTGAC | genomic sequencing primer for sgRNA target |
| mmar_4196_RIGHT | ATCAGTGCGGGTTTGTCTTC | genomic sequencing primer for sgRNA target |
| mmar_4152_RIGHT | AGTTGGAAATGTCGCCCTC | genomic sequencing primer for sgRNA target |
| mmar_4145_RIGHT | GACGAACGTGCTCTGAATGTC | genomic sequencing primer for sgRNA target |
| mmar_4038_RIGHT | GAATCTCTGTTACCGCTGCTG | genomic sequencing primer for sgRNA target |
| mmar_4190_RIGHT | GGTAATTCGGAAACAACCTTTG | genomic sequencing primer for sgRNA target |
| mmar_2188_LEFT | CGGATAACTCAGCAAAGGTTG | genomic sequencing primer for sgRNA target |
| mmar_2220_LEFT | CTACGATGCAGGGTATGCAG | genomic sequencing primer for sgRNA target |
| mmar_2225_LEFT | CTCCCTGCTGACCCAATATC | genomic sequencing primer for sgRNA target |
| mmar_2495_LEFT | GACTGTCGTTGAGCACTGTTGT | genomic sequencing primer for sgRNA target |
| mmar_0416_LEFT | GGGGTCTTACACCGTTTATGTG | genomic sequencing primer for sgRNA target |
| mmar_2364_LEFT | CTCGACACCGCCTACCAG | genomic sequencing primer for sgRNA target |
| mmar_2886_LEFT | GTCTGAATTCGCTACCACTGC | genomic sequencing primer for sgRNA target |
| mmar_4983_LEFT | GTATTCGGTCACGGTGGAAAT | genomic sequencing primer for sgRNA target |
| mmar_0422_LEFT | GGGCTCCTATCTGGTCTACAAC | genomic sequencing primer for sgRNA target |
| mmar_0809_LEFT | GTATCTTGGAATCCCGTGAACA | genomic sequencing primer for sgRNA target |
| mmar_2361_LEFT | CGGTGGTGATAACGGTGAG | genomic sequencing primer for sgRNA target |

| | | |
|---|---|---|
| mmar_2731_LEFT | CTCGCTCGATCACGATCTTT | genomic sequencing primer for sgRNA target |
| mmar_1834_LEFT | ATTTTTCTCACAACACCGATCC | genomic sequencing primer for sgRNA target |
| mmar_4139_LEFT | GGTCGACCACGACTACTTCAC | genomic sequencing primer for sgRNA target |
| mmar_5302_LEFT | GGCTGTGACGTCAACTCGAT | genomic sequencing primer for sgRNA target |
| mmar_4576_LEFT | AGAGGAGTTTTTGCGTGAAGTC | genomic sequencing primer for sgRNA target |
| mmar_4580_LEFT | AACTGAATTGAAACTCAACCGAG | genomic sequencing primer for sgRNA target |
| mmar_0525_LEFT | ACAGCCGGTGGTGTTGAG | genomic sequencing primer for sgRNA target |
| mmar_2827_LEFT | GAGTCTTTATGTGGTGAAGGGG | genomic sequencing primer for sgRNA target |
| mmar_3014_LEFT | GAGCGAAATGTTCGACAAGC | genomic sequencing primer for sgRNA target |
| mmar_2122_LEFT | AAGCAGGTGACTTGTGACGAC | genomic sequencing primer for sgRNA target |
| mmar_1840_LEFT | CTCAGCGACAACCCCAAC | genomic sequencing primer for sgRNA target |
| mmar_1900_LEFT | GTTAAGGGGTAGTGGGCCAT | genomic sequencing primer for sgRNA target |
| mmar_3619_LEFT | AGTGCACACGAATCTGGTCTC | genomic sequencing primer for sgRNA target |
| mmar_3872_LEFT | TATCTGGAAGGTCACTTGATGG | genomic sequencing primer for sgRNA target |
| mmar_2188_RIGHT | GTCGCCTTGGAACTGTCTTG | genomic sequencing primer for sgRNA target |
| mmar_2220_RIGHT | GAGCTCGACGTTTCCCTTC | genomic sequencing primer for sgRNA target |
| mmar_2225_RIGHT | CGCTGGTGAAATAGGTTTCG | genomic sequencing primer for sgRNA target |
| mmar_2495_RIGHT | GGCTGACTGTAGGAGATGCC | genomic sequencing primer for sgRNA target |
| mmar_0416_RIGHT | GTTCTTGAGTAGCTGCGACGAC | genomic sequencing primer for sgRNA target |
| mmar_2364_RIGHT | GTAGAACTGTGCAGTCAATGGG | genomic sequencing primer for sgRNA target |
| mmar_2886_RIGHT | CAGCTCGATGTGATAAGAGCC | genomic sequencing primer for sgRNA target |
| mmar_4983_RIGHT | CCAGTTCGATGTGCAGTGAC | genomic sequencing primer for sgRNA target |
| mmar_0422_RIGHT | GTAGATCTCCGACGCCTGC | genomic sequencing primer for sgRNA target |
| mmar_0809_RIGHT | GACGGATTCTTCAGCACGAC | genomic sequencing primer for sgRNA target |
| mmar_2361_RIGHT | CACCTTCGTCACAACGTCC | genomic sequencing primer for sgRNA target |
| mmar_2731_RIGHT | GTGACGATGACCAGTGTGTTG | genomic sequencing primer for sgRNA target |
| mmar_1834_RIGHT | GGCACTTGCTGTGCGTAG | genomic sequencing primer for sgRNA target |
| mmar_4139_RIGHT | CTTGACGATCATCGACTCCC | genomic sequencing primer for sgRNA target |
| mmar_5302_RIGHT | CGAAGTACAGCAAGAGGTTTCC | genomic sequencing primer for sgRNA target |
| mmar_4576_RIGHT | ATCCATTTGCGTTGTAGTCCAG | genomic sequencing primer for sgRNA target |
| mmar_4580_RIGHT | CGTGTAGTTGAGGGTCTGTCCT | genomic sequencing primer for sgRNA target |
| mmar_0525_RIGHT | CGATCAGGGACTTCATCTGC | genomic sequencing primer for sgRNA target |
| mmar_2827_RIGHT | GGTTCATCGGGATTGACG | genomic sequencing primer for sgRNA target |
| mmar_3014_RIGHT | CATCAATACATCCGGACACTTG | genomic sequencing primer for sgRNA target |
| mmar_2122_RIGHT | GAGCCTTGTTGTCCAGCAG | genomic sequencing primer for sgRNA target |
| mmar_1840_RIGHT | AGGGAAAACGTCTGGTGCTC | genomic sequencing primer for sgRNA target |
| mmar_1900_RIGHT | ACTTCGTCGTAGTCCATGCC | genomic sequencing primer for sgRNA target |
| mmar_3619_RIGHT | CCAGTAGATCCGCGTTGTAGA | genomic sequencing primer for sgRNA target |
| mmar_3872_RIGHT | CTTGCCGCTCTCGTTGAC | genomic sequencing primer for sgRNA target |
| mmar_5092_LEFT | GCACCTGACCTCCTCTTACG | genomic sequencing primer for sgRNA target |
| mmar_5092_RIGHT | GTAGAGGTTGGTTGGTGTTGGT | genomic sequencing primer for sgRNA target |
| mmar_4479_LEFT | GGATAGACGGACGTTGAATCTG | genomic sequencing primer for sgRNA target |
| mmar_3713_LEFT | CAATCCACTGGCCACTCAG | genomic sequencing primer for sgRNA target |
| mmar_4200_LEFT | GACCCTGCGGTATTTGAATC | genomic sequencing primer for sgRNA target |
| mmar_3050_LEFT | GTAAACCGGCGTCAGTTGCTA | genomic sequencing primer for sgRNA target |
| mmar_4479_RIGHT | ACACCTTCTTGACCTGGTGG | genomic sequencing primer for sgRNA target |

| | | |
|---|---|---|
| mmar_3713_RIGHT | GCTCGTTGTGCACCATCTC | genomic sequencing primer for sgRNA target |
| mmar_4200_RIGHT | CTGTTGCTTGTCCAGGAAGTC | genomic sequencing primer for sgRNA target |
| mmar_3050_RIGHT | GGATATCGGCACTGTTGTAGGT | genomic sequencing primer for sgRNA target |
| mmar_1803_LEFT | TACATATGACAGGTGCCAAACC | genomic sequencing primer for sgRNA target |
| mmar_1763_LEFT | CAATACAGCGAATATCAGCGAG | genomic sequencing primer for sgRNA target |
| mmar_1803_RIGHT | CAGTTCGATCAGCTGTTGCT | genomic sequencing primer for sgRNA target |
| mmar_1763_RIGHT | GTTGTAGGTGCAGTCACCCTT | genomic sequencing primer for sgRNA target |
| mmar_4800_LEFT | AATCCGTTCGGCCCACATT | genomic sequencing primer for sgRNA target |
| mmar_4800_RIGHT | TTGTCACAGATTCCTCGGCG | genomic sequencing primer for sgRNA target |
| mmar_4146_LEFT | CAGCAGTAGAGAGGTAAGCGGT | genomic sequencing primer for sgRNA target |
| mmar_3718_LEFT | GATCTGGTGAATTTCTCGGTC | genomic sequencing primer for sgRNA target |
| mmar_4146_RIGHT | GTACCAGGAGAAGGAGAAGTGC | genomic sequencing primer for sgRNA target |
| mmar_3718_RIGHT | CGTTCTCGTAGCTGATCAAGAC | genomic sequencing primer for sgRNA target |
| mmar_3092_LEFT | TTCACCGTCACCGGACAAC | genomic sequencing primer for sgRNA target |
| mmar_3092_RIGHT | CGTTGGCAGCGAAGTTCATC | genomic sequencing primer for sgRNA target |
| mmar_1667_LEFT | CAACAGCAGCTACACCACCAC | genomic sequencing primer for sgRNA target |
| mmar_1667_RIGHT | CGACGTAGAGGTTGTCGAATC | genomic sequencing primer for sgRNA target |
| CRISPRi-seq | TTCCTGTGAAGAGCCATTGATAATG | sequencing primer for pCRISPRx-Sth1-Cas9-L5 and pCRISPRi sgRNA |
| pSMT3-fecB.Fw | GAGGAATCACGCTAGCgtgcaattcggccgg | cloning of pSMT3-fecB |
| pSMT3-fecB.Rv | TGGCGGCCGCTCTAGAtcagttgatcggtgcgttcac | cloning of pSMT3-fecB |
| pLJR962-M5354-FLAG.Fw | gagaAGGcGGTATCGATatgcgtaatgcactggccac | cloning of pLJR962-aftA-FLAG |
| pLJR962-M5354-FLAG.Rv | AGCTAATCAGCGGCCGCTCACTTATCGTCGTCATCCTTGTAGT | cloning of pLJR962-aftA-FLAG |
| pSMT3-fecB-HA.Fw | GAGGAATCACGCTAGCgtgcaattcggccggatg | cloning of pSMT3-fecB-HA |
| pSMT3-fecB-HA.Rv | CATACGGATAGGATCCgttgatcggtgcgttcaccc | cloning of pSMT3-fecB-HA |
| pSMT3-lpqZ-HA.Fw | GAGGAATCACGCTAGCgtgaaaatcggcaggctggc | cloning of pSMT3-lpqZ-HA |
| pSMT3-lpqZ-HA.Rv | CATACGGATAGGATCCacgccccagcggatg | cloning of pSMT3-lpqZ-HA |
| pML1357kana.Fw | gatcatcattccgttaacAACGCAAAAAGCCCCCC | cloning of pML1357kana intermediate |
| pML1357kana.Rv | atctagcttagtcaatgcatTTAGAAAAACTCATCGAGCATCAAATGAAACT | cloning of pML1357kana intermediate |
| mmar1667-Strep.Rv | gaggaatcacgctagcttgggaaccctgcgaaatcatcc | cloning of pML1357kana-mmar1667-Strep |
| mmar1667-Strep.Fw | taggtcggcgacgcgtaagcttTTATTACTTCTCGAACTGCGGGTGGCTCCAGCCTGCAGGggccggcgcgaagctgtatac | cloning of pML1357kana-mmar1667-Strep |
| fecB(MTB).Fw | GAGGAATCACGCTAGCatgcgatccactgttgctgt | cloning of pSMT3-fecB(Mtb) |
| fecB(MTB).Rv | TGGCGGCCGCTCTAGActagttgatcggcgcgtcg | cloning of pSMT3-fecB(Mtb) |
| lpqZ(MTB).Fw | GAGGAATCACGCTAGCgtgagaatcaccaggatcctcgc | cloning of pSMT3-lpqZ(Mtb)-HA |
| lpqZ(MTB).Rv | CATACGGATAGGATCCacgtcccagcgggt | cloning of pSMT3-lpqZ(Mtb)-HA |
